# Targeting Anti-HLA Class I and II Antibodies with CAR-B Cell Therapy: A Novel Strategy to Mitigate Graft Rejection and Platelet Refractoriness

**DOI:** 10.1101/2025.02.20.639042

**Authors:** Varun Kesherwani

## Abstract

Alloimmune responses mediated by anti-HLA (human leukocyte antigen) antibodies are significant barriers to successful transplantation and transfusion therapies. Current immunosuppressive strategies target broad components of the immune system, often leading to non-specific effects, such as increased susceptibility to infections and malignancies. Here, we propose a novel therapeutic approach using Chimeric Antigen Receptor (CAR)-B cells engineered to target anti-HLA Class I and II antibodies. By specifically neutralizing these antibodies, CAR-B cell therapy has the potential to mitigate immune responses in allogeneic transplantation and transfusion, reducing the incidence of graft rejection and transfusion reactions. It will also neutralize the T cell response by binding to reactive T cells against donor HLA molecules. In this article, we discuss the rationale, design, and anticipated outcomes of this innovative therapy.

## Introduction

Human leukocyte antigen (HLA) mismatches between donor and recipient can trigger a cascade of immune responses, including the production of alloantibodies that target HLA molecules. Anti-HLA Class I and Class II antibodies are critical mediators of transplant rejection and transfusion complications such as hemolytic transfusion reactions (HTR). These alloantibodies, primarily produced by B cells, bind to foreign HLA antigens on donor tissues or transfused blood cells, leading to complement activation, antibody-dependent cellular cytotoxicity (ADCC), and eventually tissue destruction [1–5].

The current management of alloimmune responses relies heavily on broad immunosuppressants, including calcineurin inhibitors and corticosteroids, which suppress both T and B cell activity [6, 7]. While these therapies reduce immune rejection, they also compromise the patient’s ability to fight infections and malignancies. The advent of chimeric antigen receptor (CAR) technology has revolutionized the treatment of hematologic cancers, particularly through CAR-T cell therapies that target specific tumor antigens [8, 9]. However, CAR-B cells, with their natural ability to modulate humoral immunity, offer an untapped potential for use in transplantation and transfusion medicine.

This article explores the development of CAR-B cells engineered to target anti-HLA Class I and II antibodies. By neutralizing these antibodies, CAR-B cells could provide a targeted, less toxic alternative to broad immunosuppression, improving outcomes for transplant recipients and transfusion patients alike. CAR-B also minimize the cost associated with alternative approach that involve direct administration of fused molecules to neutralize anti-HLA antibodies [10, 11]. Also, as the with natural B cell activation, these CAR-B cells will only be activated when anti-HLA antibodies will be present the system.

The hypothesis driving this idea is that CAR-B cells engineered (in vitro or in vivo) to target and neutralize anti-HLA Class I and II antibodies can prevent alloantibody-mediated immune responses in transplantation and transfusion settings. It is anticipated that these engineered CAR-B cells will also prevent T cell-mediated injuries to the grafted organ or tissue.

The goal of this study to fill a critical gap in the current landscape of transplantation and transfusion medicine, providing a novel therapeutic strategy that directly targets the underlying cause of alloimmune responses.

### Construct Design and CAR-B Cell Engineering

CAR-B cells are designed to express a chimeric antigen receptor that specifically recognizes and binds to anti-HLA antibodies. The CAR construct (Fig. 1) is composed of:

**Fig. 1.**
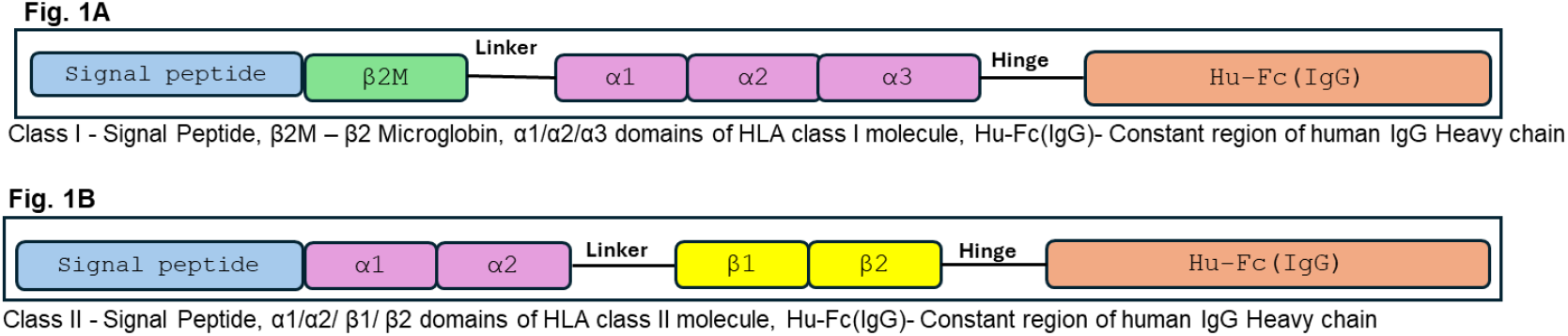

#### Extracellular HLA Domain

Derived from an HLA molecule that recognizes anti-HLA Class I and II antibodies. This domain is crucial for the specific binding of the CAR-B cells to the target alloantibodies.

#### Transmembrane Domain

Anchors the receptor to the B cell membrane and supports signal transduction.

#### Intracellular Signaling Domain

Promotes B cell activation upon engagement with the target anti-HLA antibodies, enhancing the cell’s immune-regulatory functions.

In an alternative design (Fig. 2), HLA class I and class II peptides were incorporated into the same peptide chain using a linker. This approach has been suggested in previous studies to enhance the stability of peptides in secreted molecules [8].

**Fig. 2.**
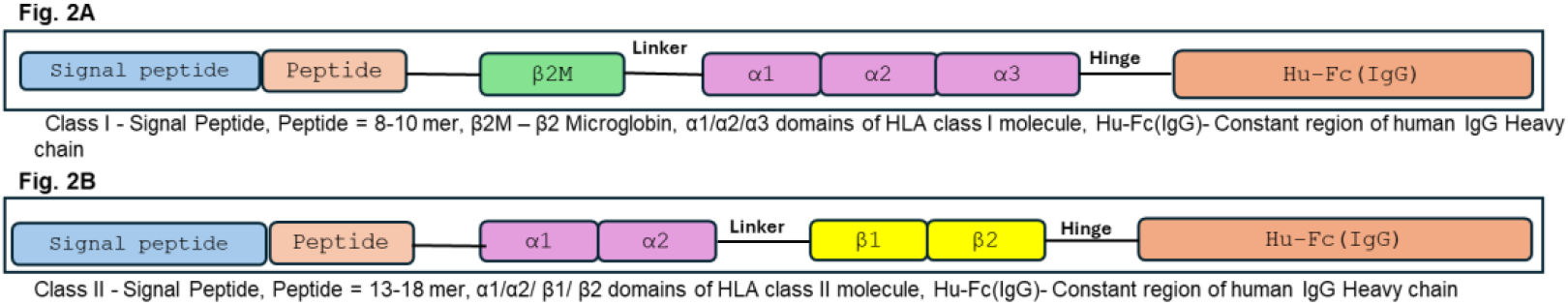

The Fig. 3 represents pictorial representation of the membrane bound and secreted fused HLA-IgG molecules.

**Fig. 3.**
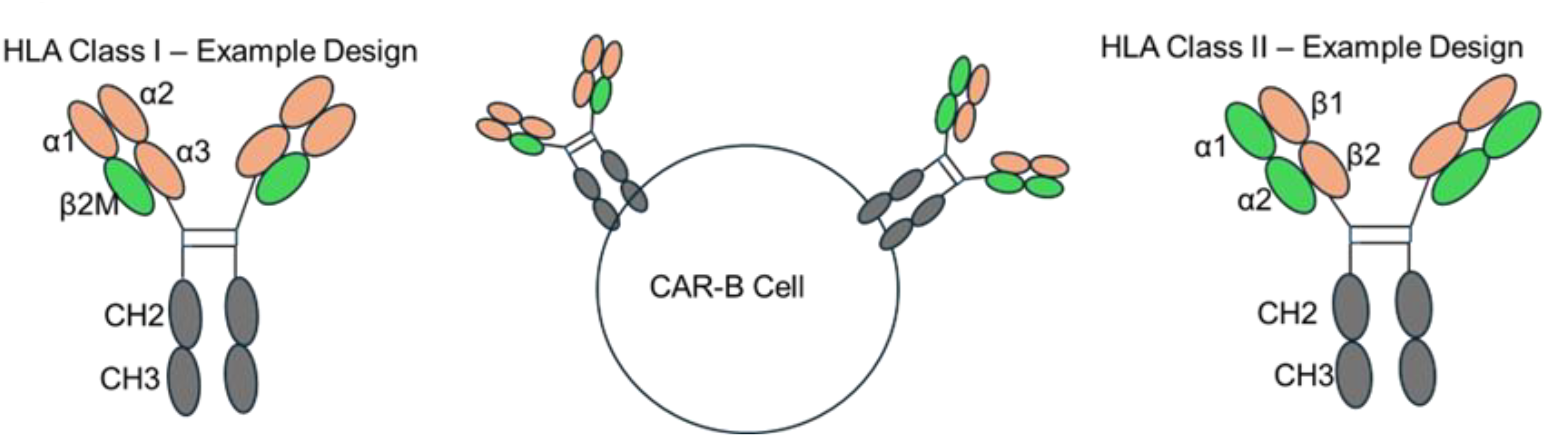

### Structural prediction using alpha fold-2 and stability of fused HLA-IgG molecules

The sequences of the CAR-B constructs used for 3D structure prediction with AlphaFold2 are listed in the supplementary file. As a proof of concept, one class I HLA-A*02:01 and class II HLA-DRB1*04:01/DRA101:01 protein extracellular domains were included. Both approaches were employed: (1) single-chain constructs for all components and (2) a separate chain for the HLA peptide in structural predictions. For class I HLA molecules, beta-2 microglobulin was incorporated, while for class II HLA molecules, the alpha chain was included within the same construct.

**Fig. 4A**: The CAR-B A*02:01P-Fc (IgG) structure was generated using the Fig. 1A approach, keeping the binding peptide in the one chain. **Fig. 4B**: This structure was generated using the binding peptide as a separate chain with AlphaFold2. **Fig. 5A**: The CAR-B DRB1*04:01P/DRA1*01:01P-Fc (IgG) structure was generated using the Fig. 1B approach, keeping the 15 mer binding peptide in the same chain. **Fig. 5B**: This structure was generated using the binding peptide as a separate chain with AlphaFold2 using approach of Fig. 2B. The rationale was generating these structural variations were to testify the proper folding and stability of different 3D structure of fused proteins.

**Fig. 4.**
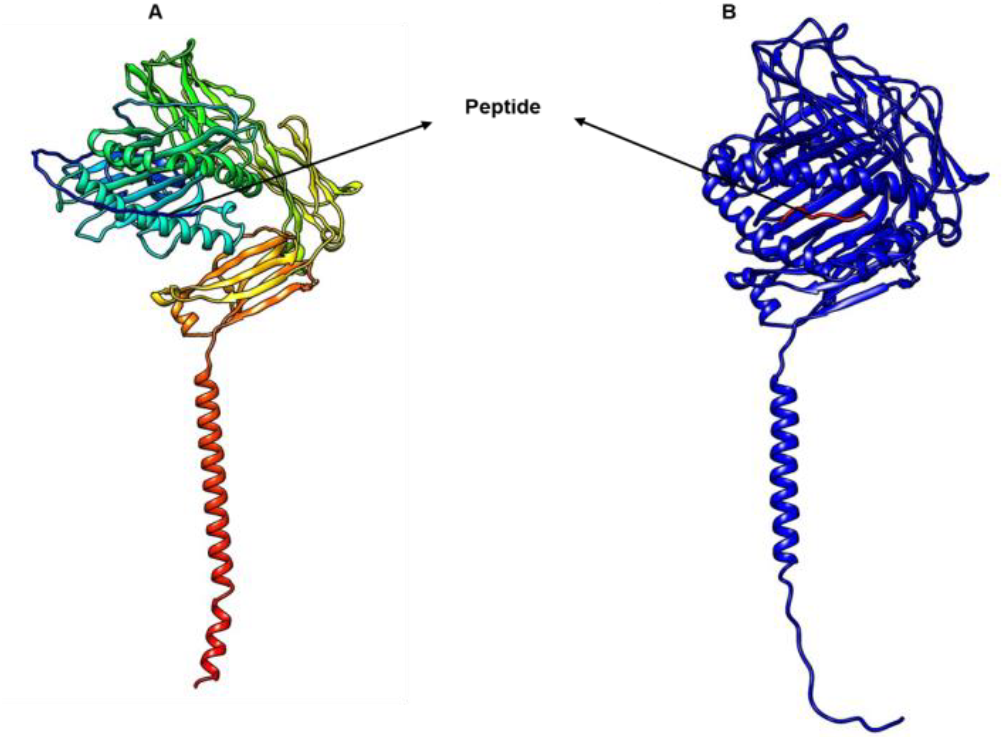
A. CAR-B A*02:01P-Fc(IgG) generated using Alphafold2 with binding peptide in one chain. B. CAR-B A*02:01P-Fc(IgG) generated using Alphafold2 with binding peptide as a separate chain.

**Fig. 5:**
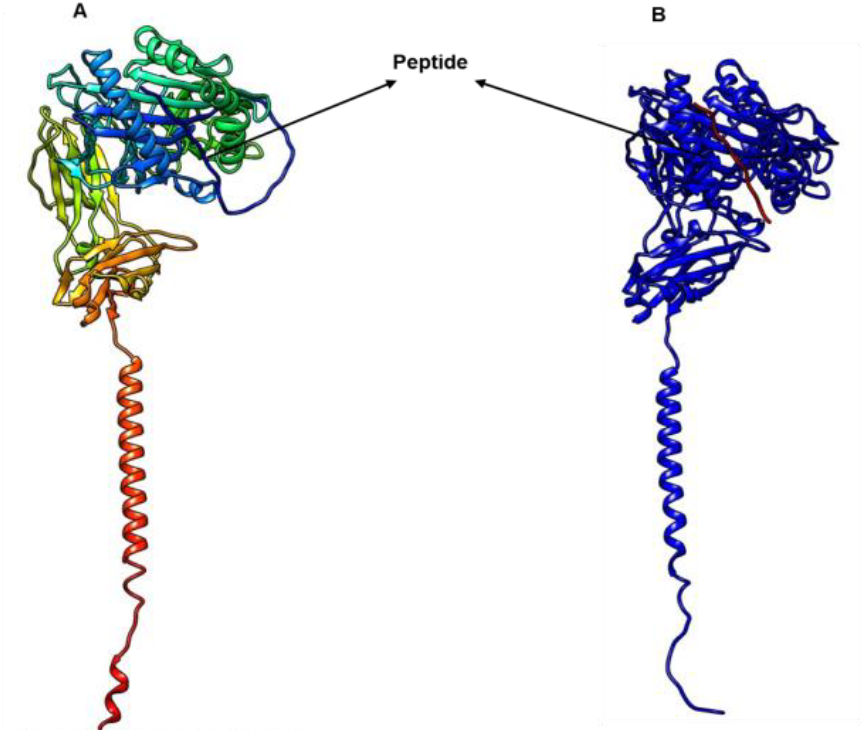
A. CAR-B DRB1*04:01P/DRA1*01:01P-Fc(IgG) generated using Alphafold2 with binding peptide in one chain. B. CAR-B DRB1*04.01P/DRA1*01:01P-Fc(IgG) generated using Alphafold2with binding peptide as a separate chain.

### Function interactions between Fused HLA-IgG molecules and anti HLA class I and class II antibodies using alpha-fold2

To find the functional interactions between fused HLA-IgG molecules and anti-HLA antibodies, two antibodies with available PDB structures in the RCSB database were used. RCSB 7TLO (W6/32), which binds to all class I HLA molecules, and 8EUQ, which binds to the HLA class II DRB1 molecule, were selected. The interaction was studied using AlphaFold2, and the interacting hydrogen bonds were identified using UCSF Chimera with default parameters.

**Fig. 6**: The CAR-B A*02:01P-Fc (IgG) with peptide in same chain and anti-HLA class I Fab antibody fragment (7t0I: W6/32) was molecularly docked using alphafold2. This structure shown below represent the pose where CAR-B A*02:01P-Fc (IgG) interacts with W6/32 extracellular domains. The number of hydrogen bonds between CAR-B and W6/32 was determined using UCSF Chimera.

**Fig. 6:**
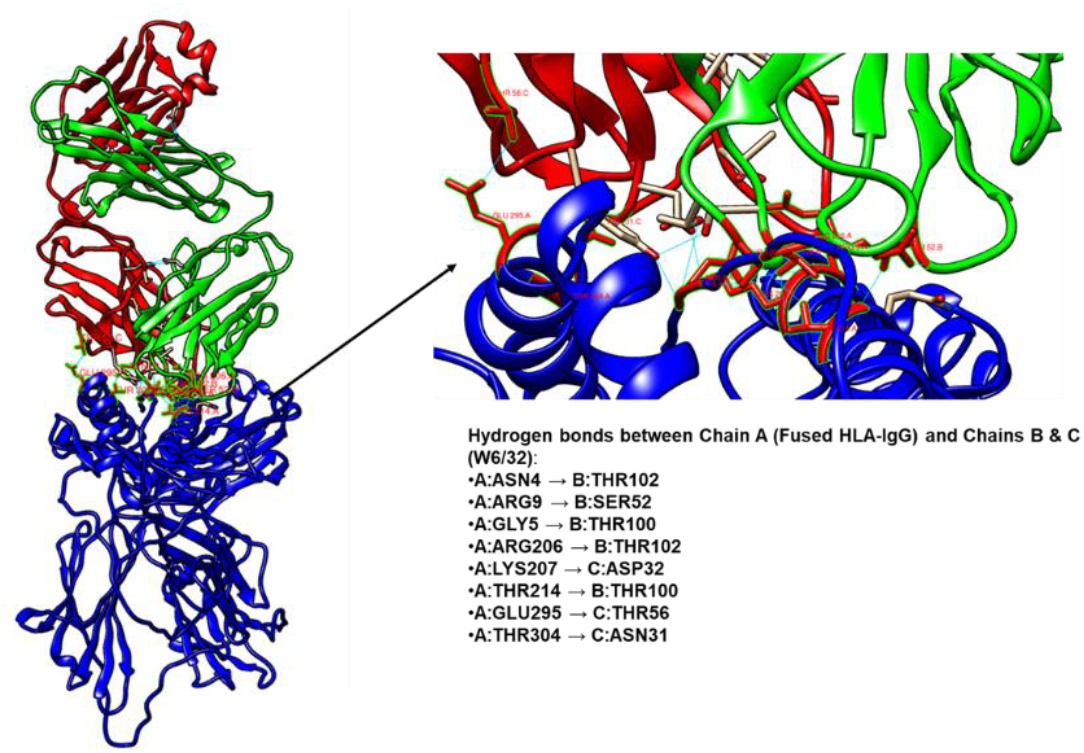
Intermolecular interactions between CAR-B A*02:0lP-Fc(IgG) and anti HLA class I antibody (RCSB-7TOL W6/32) generated using Alphafold2 Interacting H-bonds between CAR-B and W632 were identified using UCSF Chimera.

**Fig. 7**: The CAR-B DRB1*04:01P/DRA1*01:01P-Fc (IgG) with peptide in same chain and anti-HLA class II Fab antibody (8EUQ) was molecularly docked using alphafold2. This structure shown below represent the pose where CAR-B DRB1*04:01P/DRA1*01:01P-Fc (IgG) interacts with 8EUQ (anti DRB1*04:01P/DRA1*01:01P) extracellular domains. The number of hydrogen bonds between CAR-B and 8EUQ was determined using UCSF Chimera.

**Fig 7:**
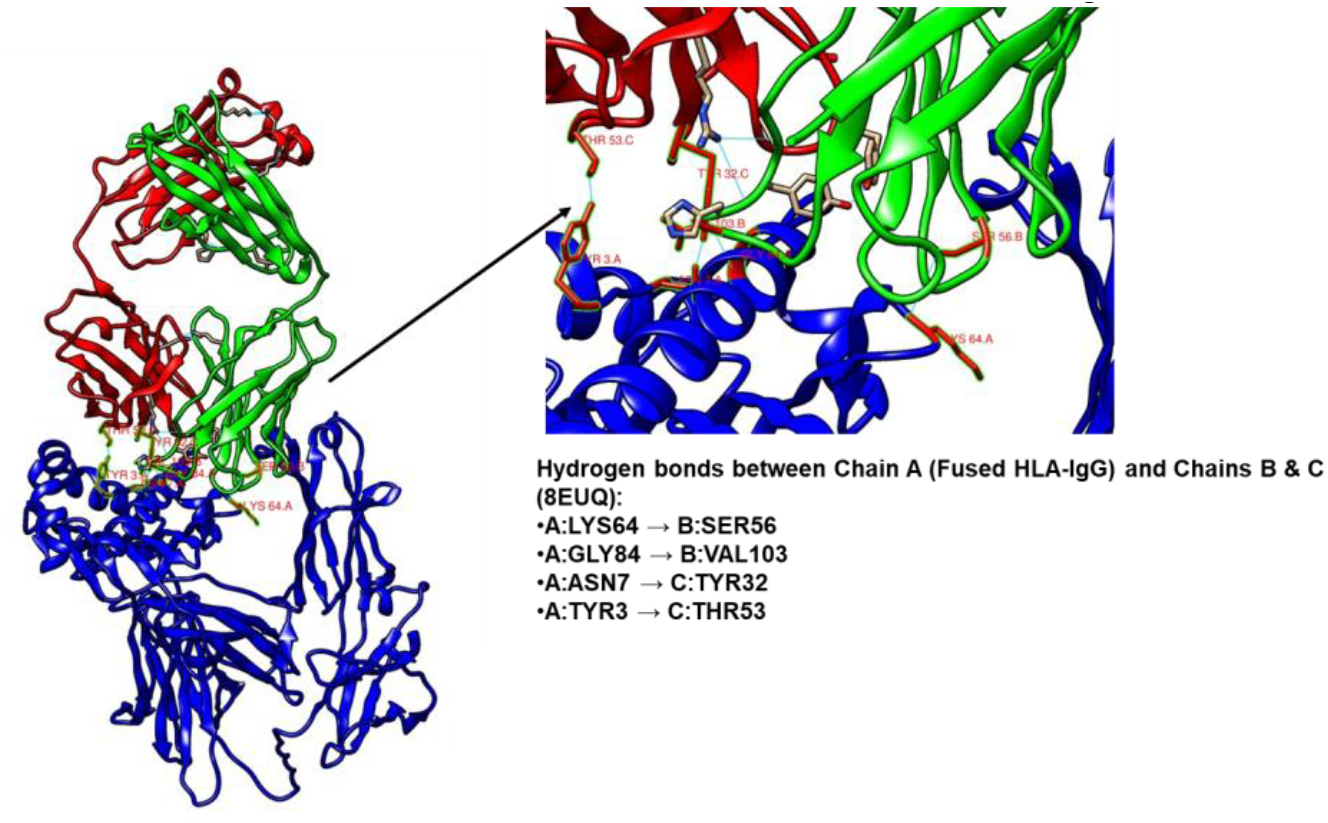
Intermolecular interactions between CAR-B DRB1*04:01 P/DRA1*01:01P-Fc(IgG) and anti HLA class II antibody (RCSB 8EUQ) generated using Alphafold2. Interacting H-bonds between CAR-B and 8EUQ were identified using UCSF Chimera.

### Future Directions

To further advance CAR-B cell therapy, comprehensive in vitro and in vivo preclinical evaluations are necessary. In vitro, CAR-B cells should be characterized through binding assays to confirm their specificity for anti-HLA antibodies, neutralization assays to assess their ability to mitigate antibody-mediated responses, and cytokine production analysis to validate their immune-modulatory potential. In vivo, efficacy should be tested in humanized mouse models of transplantation and transfusion. In the transplantation model, CAR-B cells can be infused following human graft transplantation to assess their impact on graft survival and immune tolerance, with key metrics including immune cell infiltration and anti-HLA antibody levels. In the transfusion model, CAR-B cells can be evaluated for their ability to prevent alloantibody formation and hemolysis in HLA-mismatched RBC transfusions. These studies will provide critical insights into the therapeutic potential of CAR-B cells in controlling alloimmune responses. If successful, this approach could represent a paradigm shift in transplantation and transfusion medicine by offering targeted and durable immunosuppression, reducing dependence on broad-spectrum immunosuppressive drugs and minimizing associated side effects. Earlier in vivo CAR-B engineering has been done to neutralize HIV antibodies [12]. Once thought to be difficult to engineer, several approaches have been proposed and have been used successfully to engineered B cells and transform them into plasma cells for antibody production [13–17]. The biotech company Be Biopharma has successfully advanced engineered B cell therapy to clinical trials for Hemophilia B.

This preclinical study will lay the groundwork for further exploration into the use of CAR-B cells in clinical settings. Future studies will focus on optimizing CAR constructs, improving in vivo persistence, and evaluating potential off-target effects. Additionally, the combination of CAR-B cell therapy with other immunomodulatory strategies can be explored to further enhance immune tolerance and graft acceptance.

In clinical settings, CAR-B cells will be generated by isolating peripheral blood mononuclear cells (PBMCs) from healthy donors, followed by the purification of B cells. The B cells will be then transduced with lentiviral vectors encoding the CAR constructs (Fig. 8). Following transduction, CAR expression is verified using flow cytometry, and functional assays are conducted to assess their ability to neutralize anti-HLA antibodies in vitro.

**Fig. 8:**
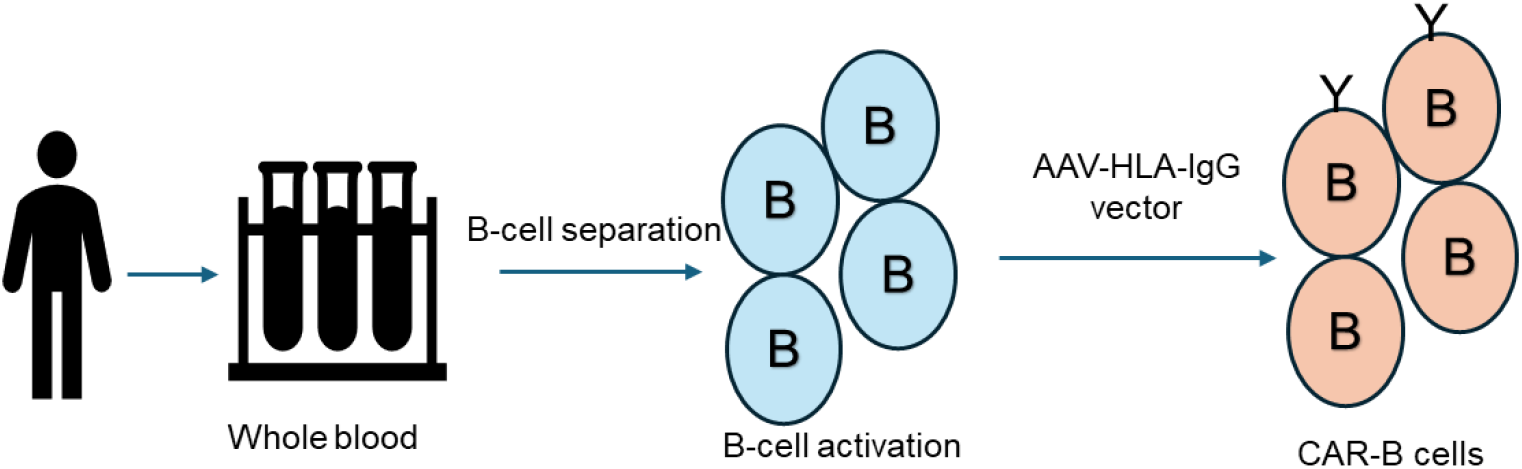
Clinical workflowfor CAR-B cells treatment

Given the potential of CAR-B cells to neutralize specific alloantibodies, this approach could be expanded to other areas of immunotherapy, including autoimmune diseases and antibody-mediated disorders beyond regular transplantation and transfusion [18–20].

## Conclusion

The development of CAR-B cells targeting anti-HLA Class I and II antibodies offers a novel and targeted approach to address the challenges of alloimmune responses in transplantation and transfusion medicine. This therapeutic strategy promises to reduce graft rejection, prevent transfusion complications, and ultimately improve patient outcomes. With further development, CAR-B cell therapy could revolutionize the field of immunotherapy, providing patients with more precise and effective treatments.

## Supplementary File

The sequence data of HLA-IgG used for AlphaFold2 in this article, along with additional HLA-IgG molecules, are provided in the supplementary file.

## Notes

### Competing Interest Statement

The authors have declared no competing interest.

