## Supplementary File for "Targeting Anti-HLA Class I and II Antibodies with CAR-B Cell Therapy: A Novel Strategy to Mitigate Graft Rejection and Platelet Refractoriness"

Example Fusion proteins sequences used in alphafold2 modelling for 3D structures shown in different figures. The full sequences of all HLAs are mentioned in annex 1.

Fig. 4A: A*02:01P

FRYNGLIHR**GGGGSGGGGSGGGGS**IQRTPKIQVYSRHPAENGKSNFLNCYVSGFHPSDIEVDLLKNGERI
EKVEHSDLSFSKDWSFYLLYYTEFTPTEKDEYACRVNHVTLSQPKIVKWGKSYILL**GGGGSGGGGSGGGG**
**S**GSHSMRYFFTSVSRPGRGEPRFIAVGYVDDTQFVRFDSDAASQRMEPRAPWIEQEGPEYWDGETRKVKA
HSQTHRVDLGTLRGYYNQSEAGSHTVQRMYGCDVGSDWRFLRGYHQYAYDGKDYIALKEDLRSWTAADMA
AQTTKHKWEAAHVAEQLRAYLEGTCVEWLRRYLENGKETLQRTDAPKTHMTHHAVSDHEATLRCWALSFY
PAEITLTWQRDGEDQTQDTELVETRPAGDGTFQKWAAVVVPSGQEQRYTCHVQHEGLPKPLTLRWEPKSC
DKTHTCPPCPAPELLGGPSVFLFPPKPKDTLMISRTPEVTCVVVDVSHEDPEVKFNWYVDGVEVHNAKTK
PREEQYNSTYRVVSVLTVLHQDWLNGKEYKCKVSNKGLPSSIEKTISKAKGQPREPQVYTLPPSRDELTK
NQVSLTCLVKGFYPSDIAVEWESNGQPENNYKTTPPVLDSDGSFFLYSKLTVDKSRWQQGNVFSCSVMHE
ALHNHYTQKSLSLSPGK**ITIFITLFLLSVCYSATVTFFKVKWIFSSVVDLKQTIIPDYRNMIGQGA**

Fig. 4B: A*02:01P

FRYNGLIHR:IQRTPKIQVYSRHPAENGKSNFLNCYVSGFHPSDIEVDLLKNGERIEKVEHSDLSFSKDW
SFYLLYYTEFTPTEKDEYACRVNHVTLSQPKIVKWGKSYILL**GGGGSGGGGSGGGGS**GSHSMRYFFTSVS
RPGRGEPRFIAVGYVDDTQFVRFDSDAASQRMEPRAPWIEQEGPEYWDGETRKVKAHSQTHRVDLGTLRG
YYNQSEAGSHTVQRMYGCDVGSDWRFLRGYHQYAYDGKDYIALKEDLRSWTAADMAAQTTKHKWEAAHVA
EQLRAYLEGTCVEWLRRYLENGKETLQRTDAPKTHMTHHAVSDHEATLRCWALSFYPAEITLTWQRDGED
QTQDTELVETRPAGDGTFQKWAAVVVPSGQEQRYTCHVQHEGLPKPLTLRWEPKSCDKTHTCPPCPAPEL
LGGPSVFLFPPKPKDTLMISRTPEVTCVVVDVSHEDPEVKFNWYVDGVEVHNAKTKPREEQYNSTYRVVS
VLTVLHQDWLNGKEYKCKVSNKGLPSSIEKTISKAKGQPREPQVYTLPPSRDELTKNQVSLTCLVKGFYP
SDIAVEWESNGQPENNYKTTPPVLDSDGSFFLYSKLTVDKSRWQQGNVFSCSVMHEALHNHYTQKSLSLS
PGK**ITIFITLFLLSVCYSATVTFFKVKWIFSSVVDLKQTIIPDYRNMIGQGA**

Fig. 5A-DRB1*04:01P/DRA1*01:01P

PKYVKQNTLKLAT**GGGGSGGGGSGGGGS**EEHVIIQAEFYLNPDQSGEFMFDFDGDEIFHVDMAKKETVWR
LEEFGRFASFEAQGALANIAVDKANLEIMTKRSNYTPITNVPPEVTVLTNSPVELREPNVLICFIDKFTP
PVVNVTWLRNGKPVTTGVSETVFLPREDHLFRKFHYLPFLPSTEDVYDCRVEHWGLDEPLLKHW**GGGGSG**
**GGGSGGGGS**PRFLEQVKHECHFFNGTERVRFLDRYFYHQEEYVRFDSDVGEYRAVTELGRPDAEYWNSQK
DLLEQKRAAVDTYCRHNYGVGESFTVQRRVYPEVTVYPAKTQPLQHHNLLVCSVNGFYPGSIEVRWFRNG
QEEKTGVVSTGLIQNGDWTFQTLVMLETVPRSGEVYTCQVEHPSLTSPLTVEWEPKSCDKTHTCPPCPAP
ELLGGPSVFLFPPKPKDTLMISRTPEVTCVVVDVSHEDPEVKFNWYVDGVEVHNAKTKPREEQYNSTYRV
VSVLTVLHQDWLNGKEYKCKVSNKGLPSSIEKTISKAKGQPREPQVYTLPPSRDELTKNQVSLTCLVKGF
YPSDIAVEWESNGQPENNYKTTPPVLDSDGSFFLYSKLTVDKSRWQQGNVFSCSVMHEALHNHYTQKSLS
LSPGK**ITIFITLFLLSVCYSATVTFFKVKWIFSSVVDLKQTIIPDYRNMIGQGA**

Fig. 5B-DRB1*04:01P/DRA1*01:01P

PKYVKQNTLKLAT:EEHVIIQAEFYLNPDQSGEFMFDFDGDEIFHVDMAKKETVWRLEEFGRFASFEAQG
ALANIAVDKANLEIMTKRSNYTPITNVPPEVTVLTNSPVELREPNVLICFIDKFTPPVVNVTWLRNGKPV
TTGVSETVFLPREDHLFRKFHYLPFLPSTEDVYDCRVEHWGLDEPLLKHW**GGGGSGGGGSGGGGS**PRFLE
QVKHECHFFNGTERVRFLDRYFYHQEEYVRFDSDVGEYRAVTELGRPDAEYWNSQKDLLEQKRAAVDTYC
RHNYGVGESFTVQRRVYPEVTVYPAKTQPLQHHNLLVCSVNGFYPGSIEVRWFRNGQEEKTGVVSTGLIQ
NGDWTFQTLVMLETVPRSGEVYTCQVEHPSLTSPLTVEWEPKSCDKTHTCPPCPAPELLGGPSVFLFPPK
PKDTLMISRTPEVTCVVVDVSHEDPEVKFNWYVDGVEVHNAKTKPREEQYNSTYRVVSVLTVLHQDWLNG
KEYKCKVSNKGLPSSIEKTISKAKGQPREPQVYTLPPSRDELTKNQVSLTCLVKGFYPSDIAVEWESNGQ
PENNYKTTPPVLDSDGSFFLYSKLTVDKSRWQQGNVFSCSVMHEALHNHYTQKSLSLSPGK**ITIFITLFL
LSVCYSATVTFFKVKWIFSSVVDLKQTIIPDYRNMIGQGA**

Fig. 6: A*02:01P: W6/32

FRYNGLIHR**GGGGSGGGGSGGGGS**IQRTPKIQVYSRHPAENGKSNFLNCYVSGFHPSDIEVDLLKNGERI
EKVEHSDLSFSKDWSFYLLYYTEFTPTEKDEYACRVNHVTLSQPKIVKWGKSYILL**GGGGSGGGGSGGGG
S**GSHSMRYFFTSVSRPGRGEPRFIAVGYVDDTQFVRFDSDAASQRMEPRAPWIEQEGPEYWDGETRKVKA
HSQTHRVDLGTLRGYYNQSEAGSHTVQRMYGCDVGSDWRFLRGYHQYAYDGKDYIALKEDLRSWTAADMA
AQTTKHKWEAAHVAEQLRAYLEGTCVEWLRRYLENGKETLQRTDAPKTHMTHHAVSDHEATLRCWALSFY
PAEITLTWQRDGEDQTQDTELVETRPAGDGTFQKWAAVVVPSGQEQRYTCHVQHEGLPKPLTLRWEPKSC
DKTHTCPPCPAPELLGGPSVFLFPPKPKDTLMISRTPEVTCVVVDVSHEDPEVKFNWYVDGVEVHNAKTK
PREEQYNSTYRVVSVLTVLHQDWLNGKEYKCKVSNKGLPSSIEKTISKAKGQPREPQVYTLPPSRDELTK
NQVSLTCLVKGFYPSDIAVEWESNGQPENNYKTTPPVLDSDGSFFLYSKLTVDKSRWQQGNVFSCSVMHE
ALHNHYTQKSLSLSPGK:VQLKQSGPGLVQPSQSLSLTCTVSGFSLTSYGVHWVRQPPGKGLEWLGVIWS
GGSTDYNAAFISRLSIRKDNSKSQVFFKMNSLQADDTAIYYCARTFTTSTSAWFAYWGQGTLVTVSAAKT
TAPSVYPLAPVCGDTTGSSVTLGCLVKGYFPEPVTLTWNSGSLSSGVHTFPAVLQSDLYTLSSSVTVTSS
TWPSQSITCNVAHPASSTKVDKKIEP:SIVMTQTPKFLLVSAGDRVTITCKASQSVSNDVAWYQQKPGQS
PKLLIYYASNRYTGVPDRFTGSGYGTDFTFTISTVQAEDLAVYFCQQDYSSPPWTFGGGTKLEIRRADAA
PTVSIFPPSSEQLTSGGASVVCFLNNFYPKDINVKWKIDGSERQNGVLNSWTDQDSKDSTYSMSSTLTLT
KDEYERHNSYTCEATHKTSTSPIVKSFN

Fig. 7-DRB1*04:01P/DRA1*01:01P:8EUQ

PKYVKQNTLKLAT**GGGGSGGGGSGGGGS**EEHVIIQAEFYLNPDQSGEFMFDFDGDEIFHVDMAKKETVWR
LEEFGRFASFEAQGALANIAVDKANLEIMTKRSNYTPITNVPPEVTVLTNSPVELREPNVLICFIDKFTP
PVVNVTWLRNGKPVTTGVSETVFLPREDHLFRKFHYLPFLPSTEDVYDCRVEHWGLDEPLLKHW**GGGGSG
GGGSGGGGS**PRFLEQVKHECHFFNGTERVRFLDRYFYHQEEYVRFDSDVGEYRAVTELGRPDAEYWNSQK
DLLEQKRAAVDTYCRHNYGVGESFTVQRRVYPEVTVYPAKTQPLQHHNLLVCSVNGFYPGSIEVRWFRNG
QEEKTGVVSTGLIQNGDWTFQTLVMLETVPRSGEVYTCQVEHPSLTSPLTVEWEPKSCDKTHTCPPCPAP
ELLGGPSVFLFPPKPKDTLMISRTPEVTCVVVDVSHEDPEVKFNWYVDGVEVHNAKTKPREEQYNSTYRV
VSVLTVLHQDWLNGKEYKCKVSNKGLPSSIEKTISKAKGQPREPQVYTLPPSRDELTKNQVSLTCLVKGF
YPSDIAVEWESNGQPENNYKTTPPVLDSDGSFFLYSKLTVDKSRWQQGNVFSCSVMHEALHNHYTQKSLS
LSPGK:DIQMTQSPSSLSASLGQRVSLTCRASQEISGYLTWLQQKPDGTIKRLVYAASTLDSGVPKRFSG
SRSGSDYSLTISSLESEDFADYYCLQYTNYPLTFGAGTKLELKRTVAAPSVFIFPPSDEQLKSGTASVVC
LLNNFYPREAKVQWKVDNALQSGNSQESVTEQDSKDSTYSLSSTLTLSKADYEKHKVYACEVTHQGLSSP
VTKSFNRGEC:QVQLKESGPGLVAPSQSLSITCTVSGFSLTSYGVHWVRQPPGKGLEWLGVIWAGGSINY
NSALMSRLSISKDNFKSQVFLKMSSLQTDDTAMYYCARAYGDYVHYAMDYWGQGTSVTASSASTKGPSVF

PLAPSSKSTSGGTAALGCLVKDYFPEPVTVSWNSGALTSGVHTFPAVLQSSGLYSLSSVVTVPSSSLGTQ
TYICNVNHKPSNTKVDKKVEPKSC

Annex 1:

The binding peptides can be added in a single chain or in the separate chain. For class II few amino acids from exon 1 have been excluded but can be added if required. Alternate Linker and transmembrane sequences will be used during further validation.

>A0101P-Final-298-Final
**MSRSVALAVLALLSLSGLEA**IQRTPKIQVYSRHPAENGKSNFLNCYVSGFHPSDIEVDLLKNGERIEKVE
HSDLSFSKDWSFYLLYYTEFTPTEKDEYACRVNHVTLSQPKIVKWGKSYILL**GGGGSGGGGSGGGGS**GSH
SMRYFFTSVSRPGRGEPRFIAVGYVDDTQFVRFDSDAASQKMEPRAPWIEQEGPEYWDQETRNMKAHSQT
DRANLGTLRGYYNQSEDGSHTIQIMYGCDVGPDGRFLRGYRQDAYDGKDYIALNEDLRSWTAADMAAQIT
KRKWEAVHAAEQRRVYLEGRCVDGLRRYLENGKETLQRTDPPKTHMTHHPISDHEATLRCWALGFYPAEI
TLTWQRDGEDQTQDTELVETRPAGDGTFQKWAAVVVPSGEEQRYTCHVQHEGLPKPLTLRWEPKSCDKTH
TCPPCPAPELLGGPSVFLFPPKPKDTLMISRTPEVTCVVVDVSHEDPEVKFNWYVDGVEVHNAKTKPREE
QYNSTYRVVSVLTVLHQDWLNGKEYKCKVSNKGLPSSIEKTISKAKGQPREPQVYTLPPSRDELTKNQVS
LTCLVKGFYPSDIAVEWESNGQPENNYKTTPPVLDSDGSFFLYSKLTVDKSRWQQGNVFSCSVMHEALHN
HYTQKSLSLSPGK**ITIFITLFLLSVCYSATVTFFKVKWIFSSVVDLKQTIIPDYRNMIGQGA**

>A02:01P-Final

**MSRSVALAVLALLSLSGLEA**IQRTPKIQVYSRHPAENGKSNFLNCYVSGFHPSDIEVDLLKNGERIEKVE
HSDLSFSKDWSFYLLYYTEFTPTEKDEYACRVNHVTLSQPKIVKWGKSYILL**GGGGSGGGGSGGGGS**GSH
SMRYFFTSVSRPGRGEPRFIAVGYVDDTQFVRFDSDAASQRMEPRAPWIEQEGPEYWDGETRKVKAHSQT
HRVDLGTLRGYYNQSEAGSHTVQRMYGCDVGSDWRFLRGYHQYAYDGKDYIALKEDLRSWTAADMAAQTT
KHKWEAAHVAEQLRAYLEGTCVEWLRRYLENGKETLQRTDAPKTHMTHHAVSDHEATLRCWALSFYPAEI
TLTWQRDGEDQTQDTELVETRPAGDGTFQKWAAVVVPSGQEQRYTCHVQHEGLPKPLTLRWEPKSCDKTH
TCPPCPAPELLGGPSVFLFPPKPKDTLMISRTPEVTCVVVDVSHEDPEVKFNWYVDGVEVHNAKTKPREE
QYNSTYRVVSVLTVLHQDWLNGKEYKCKVSNKGLPSSIEKTISKAKGQPREPQVYTLPPSRDELTKNQVS
LTCLVKGFYPSDIAVEWESNGQPENNYKTTPPVLDSDGSFFLYSKLTVDKSRWQQGNVFSCSVMHEALHN
HYTQKSLSLSPGK**ITIFITLFLLSVCYSATVTFFKVKWIFSSVVDLKQTIIPDYRNMIGQGA**

>A*02:03P-Final

**MSRSVALAVLALLSLSGLEA**IQRTPKIQVYSRHPAENGKSNFLNCYVSGFHPSDIEVDLLKNGERIEKVE
HSDLSFSKDWSFYLLYYTEFTPTEKDEYACRVNHVTLSQPKIVKWGKSYILL**GGGGSGGGGSGGGGS**GSH
SMRYFFTSVSRPGRGEPRFIAVGYVDDTQFVRFDSDAASQRMEPRAPWIEQEGPEYWDGETRKVKAHSQT
HRVDLGTLRGYYNQSEAGSHTVQRMYGCDVGSDWRFLRGYHQYAYDGKDYIALKEDLRSWTAADMAAQTT
KHKWETAHEAEQWRAYLEGTCVEWLRRYLENGKETLQRTDAPKTHMTHHAVSDHEATLRCWALSFYPAEI

TLTWQRDGEDQTQDTELVETRPAGDGTFQKWAAVVVPSGQEQRYTCHVQHEGLPKPLTLRWEPKSCDKTH
TCPPCPAPELLGGPSVFLFPPKPKDTLMISRTPEVTCVVVDVSHEDPEVKFNWYVDGVEVHNAKTKPREE
QYNSTYRVVSVLTVLHQDWLNGKEYKCKVSNKGLPSSIEKTISKAKGQPREPQVYTLPPSRDELTKNQVS
LTCLVKGFYPSDIAVEWESNGQPENNYKTTPPVLDSDGSFFLYSKLTVDKSRWQQGNVFSCSVMHEALHN
HYTQKSLSLSPGK**ITIFITLFLLSVCYSATVTFFKVKWIFSSVVDLKQTIIPDYRNMIGQGA**

>A*02:06P-Final

**MSRSVALAVLALLSLSGLEA**IQRTPKIQVYSRHPAENGKSNFLNCYVSGFHPSDIEVDLLKNGERIEKVE
HSDLSFSKDWSFYLLYYTEFTPTEKDEYACRVNHVTLSQPKIVKWGKSYILL**GGGGSGGGGSGGGGS**GSH
SMRYFYTSVSRPGRGEPRFIAVGYVDDTQFVRFDSDAASQRMEPRAPWIEQEGPEYWDGETRKVKAHSQT
HRVDLGTLRGYYNQSEAGSHTVQRMYGCDVGSDWRFLRGYHQYAYDGKDYIALKEDLRSWTAADMAAQTT
KHKWEAAHVAEQLRAYLEGTCVEWLRRYLENGKETLQRTDAPKTHMTHHAVSDHEATLRCWALSFYPAEI
TLTWQRDGEDQTQDTELVETRPAGDGTFQKWAAVVVPSGQEQRYTCHVQHEGLPKPLTLRWEPKSCDKTH
TCPPCPAPELLGGPSVFLFPPKPKDTLMISRTPEVTCVVVDVSHEDPEVKFNWYVDGVEVHNAKTKPREE
QYNSTYRVVSVLTVLHQDWLNGKEYKCKVSNKGLPSSIEKTISKAKGQPREPQVYTLPPSRDELTKNQVS
LTCLVKGFYPSDIAVEWESNGQPENNYKTTPPVLDSDGSFFLYSKLTVDKSRWQQGNVFSCSVMHEALHN
HYTQKSLSLSPGK**ITIFITLFLLSVCYSATVTFFKVKWIFSSVVDLKQTIIPDYRNMIGQGA**

>A03:01P-Final

**MSRSVALAVLALLSLSGLEA**IQRTPKIQVYSRHPAENGKSNFLNCYVSGFHPSDIEVDLLKNGERIEKVE
HSDLSFSKDWSFYLLYYTEFTPTEKDEYACRVNHVTLSQPKIVKWGKSYILL**GGGGSGGGGSGGGGS**GSH
SMRYFFTSVSRPGRGEPRFIAVGYVDDTQFVRFDSDAASQRMEPRAPWIEQEGPEYWDQETRNVKAQSQT
DRVDLGTLRGYYNQSEAGSHTIQIMYGCDVGSDGRFLRGYRQDAYDGKDYIALNEDLRSWTAADMAAQIT
KRKWEAAHEAEQLRAYLDGTCVEWLRRYLENGKETLQRTDPPKTHMTHHPISDHEATLRCWALGFYPAEI
TLTWQRDGEDQTQDTELVETRPAGDGTFQKWAAVVVPSGEEQRYTCHVQHEGLPKPLTLRWEPKSCDKTH
TCPPCPAPELLGGPSVFLFPPKPKDTLMISRTPEVTCVVVDVSHEDPEVKFNWYVDGVEVHNAKTKPREE
QYNSTYRVVSVLTVLHQDWLNGKEYKCKVSNKGLPSSIEKTISKAKGQPREPQVYTLPPSRDELTKNQVS
LTCLVKGFYPSDIAVEWESNGQPENNYKTTPPVLDSDGSFFLYSKLTVDKSRWQQGNVFSCSVMHEALHN
HYTQKSLSLSPGK**ITIFITLFLLSVCYSATVTFFKVKWIFSSVVDLKQTIIPDYRNMIGQGA**

> A11:01P-Final

**MSRSVALAVLALLSLSGLEA**IQRTPKIQVYSRHPAENGKSNFLNCYVSGFHPSDIEVDLLKNGERIEKVE
HSDLSFSKDWSFYLLYYTEFTPTEKDEYACRVNHVTLSQPKIVKWGKSYILL**GGGGSGGGGSGGGGS**GSH
SMRYFYTSVSRPGRGEPRFIAVGYVDDTQFVRFDSDAASQRMEPRAPWIEQEGPEYWDQETRNVKAQSQT
DRVDLGTLRGYYNQSEDGSHTIQIMYGCDVGPDGRFLRGYRQDAYDGKDYIALNEDLRSWTAADMAAQIT
KRKWEAAHAAEQQRAYLEGRCVEWLRRYLENGKETLQRTDPPKTHMTHHPISDHEATLRCWALGFYPAEI
TLTWQRDGEDQTQDTELVETRPAGDGTFQKWAAVVVPSGEEQRYTCHVQHEGLPKPLTLRWEPKSCDKTH
TCPPCPAPELLGGPSVFLFPPKPKDTLMISRTPEVTCVVVDVSHEDPEVKFNWYVDGVEVHNAKTKPREE
QYNSTYRVVSVLTVLHQDWLNGKEYKCKVSNKGLPSSIEKTISKAKGQPREPQVYTLPPSRDELTKNQVS
LTCLVKGFYPSDIAVEWESNGQPENNYKTTPPVLDSDGSFFLYSKLTVDKSRWQQGNVFSCSVMHEALHN
HYTQKSLSLSPGK**ITIFITLFLLSVCYSATVTFFKVKWIFSSVVDLKQTIIPDYRNMIGQGA**

>A*11:02P-Final

**MSRSVALAVLALLSLSGLEA**IQRTPKIQVYSRHPAENGKSNFLNCYVSGFHPSDIEVDLLKNGERIEKVE
HSDLSFSKDWSFYLLYYTEFTPTEKDEYACRVNHVTLSQPKIVKWGKSYILL**GGGGSGGGGSGGGGS**GSH
SMRYFYTSVSRPGRGKPRFIAVGYVDDTQFVRFDSDAASQRMEPRAPWIEQEGPEYWDQETRNVKAQSQT
DRVDLGTLRGYYNQSEDGSHTIQIMYGCDVGPDGRFLRGYRQDAYDGKDYIALNEDLRSWTAADMAAQIT
KRKWEAAHAAEQQRAYLEGRCVEWLRRYLENGKETLQRTDPPKTHMTHHPISDHEATLRCWALGFYPAEI
TLTWQRDGEDQTQDTELVETRPAGDGTFQKWAAVVVPSGEEQRYTCHVQHEGLPKPLTLRWEPKSCDKTH
TCPPCPAPELLGGPSVFLFPPKPKDTLMISRTPEVTCVVVDVSHEDPEVKFNWYVDGVEVHNAKTKPREE
QYNSTYRVVSVLTVLHQDWLNGKEYKCKVSNKGLPSSIEKTISKAKGQPREPQVYTLPPSRDELTKNQVS
LTCLVKGFYPSDIAVEWESNGQPENNYKTTPPVLDSDGSFFLYSKLTVDKSRWQQGNVFSCSVMHEALHN
HYTQKSLSLSPGK**ITIFITLFLLSVCYSATVTFFKVKWIFSSVVDLKQTIIPDYRNMIGQGA**

>A*23:01P-Final

**MSRSVALAVLALLSLSGLEA**IQRTPKIQVYSRHPAENGKSNFLNCYVSGFHPSDIEVDLLKNGERIEKVE
HSDLSFSKDWSFYLLYYTEFTPTEKDEYACRVNHVTLSQPKIVKWGKSYILL**GGGGSGGGGSGGGGS**GSH
SMRYFSTSVSRPGRGEPRFIAVGYVDDTQFVRFDSDAASQRMEPRAPWIEQEGPEYWDEETGKVKAHSQT
DRENLRIALRYYNQSEAGSHTLQMMFGCDVGSDGRFLRGYHQYAYDGKDYIALKEDLRSWTAADMAAQIT
QRKWEAARVAEQLRAYLEGTCVDGLRRYLENGKETLQRTDPPKTHMTHHPISDHEATLRCWALGFYPAEI
TLTWQRDGEDQTQDTELVETRPAGDGTFQKWAAVVVPSGEEQRYTCHVQHEGLPKPLTLRWEPKSCDKTH
TCPPCPAPELLGGPSVFLFPPKPKDTLMISRTPEVTCVVVDVSHEDPEVKFNWYVDGVEVHNAKTKPREE
QYNSTYRVVSVLTVLHQDWLNGKEYKCKVSNKGLPSSIEKTISKAKGQPREPQVYTLPPSRDELTKNQVS
LTCLVKGFYPSDIAVEWESNGQPENNYKTTPPVLDSDGSFFLYSKLTVDKSRWQQGNVFSCSVMHEALHN
HYTQKSLSLSPGK**ITIFITLFLLSVCYSATVTFFKVKWIFSSVVDLKQTIIPDYRNMIGQGA**

>A24:02P-Final

**MSRSVALAVLALLSLSGLEA**IQRTPKIQVYSRHPAENGKSNFLNCYVSGFHPSDIEVDLLKNGERIEKVE
HSDLSFSKDWSFYLLYYTEFTPTEKDEYACRVNHVTLSQPKIVKWGKSYILL**GGGGSGGGGSGGGGS**GSH
SMRYFSTSVSRPGRGEPRFIAVGYVDDTQFVRFDSDAASQRMEPRAPWIEQEGPEYWDEETGKVKAHSQT
DRENLRIALRYYNQSEAGSHTLQMMFGCDVGSDGRFLRGYHQYAYDGKDYIALKEDLRSWTAADMAAQIT
KRKWEAAHVAEQQRAYLEGTCVDGLRRYLENGKETLQRTDPPKTHMTHHPISDHEATLRCWALGFYPAEI
TLTWQRDGEDQTQDTELVETRPAGDGTFQKWAAVVVPSGEEQRYTCHVQHEGLPKPLTLRWEPKSCDKTH
TCPPCPAPELLGGPSVFLFPPKPKDTLMISRTPEVTCVVVDVSHEDPEVKFNWYVDGVEVHNAKTKPREE
QYNSTYRVVSVLTVLHQDWLNGKEYKCKVSNKGLPSSIEKTISKAKGQPREPQVYTLPPSRDELTKNQVS
LTCLVKGFYPSDIAVEWESNGQPENNYKTTPPVLDSDGSFFLYSKLTVDKSRWQQGNVFSCSVMHEALHN
HYTQKSLSLSPGK**ITIFITLFLLSVCYSATVTFFKVKWIFSSVVDLKQTIIPDYRNMIGQGA**

>A*24:03P-Final

**MSRSVALAVLALLSLSGLEA**IQRTPKIQVYSRHPAENGKSNFLNCYVSGFHPSDIEVDLLKNGERIEKVE
HSDLSFSKDWSFYLLYYTEFTPTEKDEYACRVNHVTLSQPKIVKWGKSYILL**GGGGSGGGGSGGGGS**GSH
SMRYFSTSVSRPGRGEPRFIAVGYVDDTQFVRFDSDAASQRMEPRAPWIEQEGPEYWDEETGKVKAHSQT

DRENLRIALRYYNQSEAGSHTLQMMFGCDVGSDGRFLRGYHQYAYDGKDYIALKEDLRSWTAADMAAQIT
KRKWEAAHVAEQQRAYLEGTCVEWLRRYLENGKETLQRTDPPKTHMTHHPISDHEATLRCWALGFYPAEI
TLTWQRDGEDQTQDTELVETRPAGDGTFQKWAAVVVPSGEEQRYTCHVQHEGLPKPLTLRWEPKSCDKTH
TCPPCPAPELLGGPSVFLFPPKPKDTLMISRTPEVTCVVVDVSHEDPEVKFNWYVDGVEVHNAKTKPREE
QYNSTYRVVSVLTVLHQDWLNGKEYKCKVSNKGLPSSIEKTISKAKGQPREPQVYTLPPSRDELTKNQVS
LTCLVKGFYPSDIAVEWESNGQPENNYKTTPPVLDSDGSFFLYSKLTVDKSRWQQGNVFSCSVMHEALHN
HYTQKSLSLSPGK**ITIFITLFLLSVCYSATVTFFKVKWIFSSVVDLKQTIIPDYRNMIGQGA**

> A*25:01P-Final

**MSRSVALAVLALLSLSGLEA**IQRTPKIQVYSRHPAENGKSNFLNCYVSGFHPSDIEVDLLKNGERIEKVE
HSDLSFSKDWSFYLLYYTEFTPTEKDEYACRVNHVTLSQPKIVKWGKSYILL**GGGGSGGGGSGGGGS**GSH
SMRYFYTSVSRPGRGEPRFIAVGYVDDTQFVRFDSDAASQRMEPRAPWIEQEGPEYWDRNTRNVKAHSQT
DRESLRIALRYYNQSEDGSHTIQRMYGCDVGPDGRFLRGYQQDAYDGKDYIALNEDLRSWTAADMAAQIT
QRKWETAHEAEQWRAYLEGRCVEWLRRYLENGKETLQRTDAPKTHMTHHAVSDHEATLRCWALSFYPAEI
TLTWQRDGEDQTQDTELVETRPAGDGTFQKWASVVVPSGQEQRYTCHVQHEGLPKPLTLRWEPKSCDKTH
TCPPCPAPELLGGPSVFLFPPKPKDTLMISRTPEVTCVVVDVSHEDPEVKFNWYVDGVEVHNAKTKPREE
QYNSTYRVVSVLTVLHQDWLNGKEYKCKVSNKGLPSSIEKTISKAKGQPREPQVYTLPPSRDELTKNQVS
LTCLVKGFYPSDIAVEWESNGQPENNYKTTPPVLDSDGSFFLYSKLTVDKSRWQQGNVFSCSVMHEALHN
HYTQKSLSLSPGK**ITIFITLFLLSVCYSATVTFFKVKWIFSSVVDLKQTIIPDYRNMIGQGA**

>A*26:01P-Final

**MSRSVALAVLALLSLSGLEA**IQRTPKIQVYSRHPAENGKSNFLNCYVSGFHPSDIEVDLLKNGERIEKVE
HSDLSFSKDWSFYLLYYTEFTPTEKDEYACRVNHVTLSQPKIVKWGKSYILL**GGGGSGGGGSGGGGS**GSH
SMRYFYTSVSRPGRGEPRFIAVGYVDDTQFVRFDSDAASQRMEPRAPWIEQEGPEYWDRNTRNVKAHSQT
DRANLGTLRGYYNQSEDGSHTIQRMYGCDVGPDGRFLRGYQQDAYDGKDYIALNEDLRSWTAADMAAQIT
QRKWETAHEAEQWRAYLEGRCVEWLRRYLENGKETLQRTDAPKTHMTHHAVSDHEATLRCWALSFYPAEI
TLTWQRDGEDQTQDTELVETRPAGDGTFQKWASVVVPSGQEQRYTCHVQHEGLPKPLTLRWEPKSCDKTH
TCPPCPAPELLGGPSVFLFPPKPKDTLMISRTPEVTCVVVDVSHEDPEVKFNWYVDGVEVHNAKTKPREE
QYNSTYRVVSVLTVLHQDWLNGKEYKCKVSNKGLPSSIEKTISKAKGQPREPQVYTLPPSRDELTKNQVS
LTCLVKGFYPSDIAVEWESNGQPENNYKTTPPVLDSDGSFFLYSKLTVDKSRWQQGNVFSCSVMHEALHN
HYTQKSLSLSPGK**ITIFITLFLLSVCYSATVTFFKVKWIFSSVVDLKQTIIPDYRNMIGQGA**

> A*29:01P-Final

**MSRSVALAVLALLSLSGLEA**IQRTPKIQVYSRHPAENGKSNFLNCYVSGFHPSDIEVDLLKNGERIEKVE
HSDLSFSKDWSFYLLYYTEFTPTEKDEYACRVNHVTLSQPKIVKWGKSYILL**GGGGSGGGGSGGGGS**GSH
SMRYFTTSVSRPGRGEPRFIAVGYVDDTQFVRFDSDAASQRMEPRAPWIEQEGPEYWDLQTRNVKAQSQT
DRANLGTLRGYYNQSEAGSHTIQMMYGCHVGSDGRFLRGYRQDAYDGKDYIALNEDLRSWTAADMAAQIT
QRKWEAARVAEQLRAYLEGTCVEWLRRYLENGKETLQRTDAPKTHMTHHAVSDHEATLRCWALSFYPAEI
TLTWQRDGEDQTQDTELVETRPAGDGTFQKWASVVVPSGQEQRYTCHVQHEGLPKPLTLRWEPKSCDKTH
TCPPCPAPELLGGPSVFLFPPKPKDTLMISRTPEVTCVVVDVSHEDPEVKFNWYVDGVEVHNAKTKPREE
QYNSTYRVVSVLTVLHQDWLNGKEYKCKVSNKGLPSSIEKTISKAKGQPREPQVYTLPPSRDELTKNQVS
LTCLVKGFYPSDIAVEWESNGQPENNYKTTPPVLDSDGSFFLYSKLTVDKSRWQQGNVFSCSVMHEALHN
HYTQKSLSLSPGK**ITIFITLFLLSVCYSATVTFFKVKWIFSSVVDLKQTIIPDYRNMIGQGA**

> A*29:02P-Final

**MSRSVALAVLALLSLSGLEA**IQRTPKIQVYSRHPAENGKSNFLNCYVSGFHPSDIEVDLLKNGERIEKVE
HSDLSFSKDWSFYLLYYTEFTPTEKDEYACRVNHVTLSQPKIVKWGKSYILL**GGGGSGGGGSGGGGS**GSH
SMRYFTTSVSRPGRGEPRFIAVGYVDDTQFVRFDSDAASQRMEPRAPWIEQEGPEYWDLQTRNVKAQSQT
DRANLGTLRGYYNQSEAGSHTIQMMYGCDVGSDGRFLRGYRQDAYDGKDYIALNEDLRSWTAADMAAQIT
QRKWEAARVAEQLRAYLEGTCVEWLRRYLENGKETLQRTDAPKTHMTHHAVSDHEATLRCWALSFYPAEI
TLTWQRDGEDQTQDTELVETRPAGDGTFQKWASVVVPSGQEQRYTCHVQHEGLPKPLTLRWEPKSCDKTH
TCPPCPAPELLGGPSVFLFPPKPKDTLMISRTPEVTCVVVDVSHEDPEVKFNWYVDGVEVHNAKTKPREE
QYNSTYRVVSVLTVLHQDWLNGKEYKCKVSNKGLPSSIEKTISKAKGQPREPQVYTLPPSRDELTKNQVS
LTCLVKGFYPSDIAVEWESNGQPENNYKTTPPVLDSDGSFFLYSKLTVDKSRWQQGNVFSCSVMHEALHN
HYTQKSLSLSPGK**ITIFITLFLLSVCYSATVTFFKVKWIFSSVVDLKQTIIPDYRNMIGQGA**

> A*30:01P-Final

**MSRSVALAVLALLSLSGLEA**IQRTPKIQVYSRHPAENGKSNFLNCYVSGFHPSDIEVDLLKNGERIEKVE
HSDLSFSKDWSFYLLYYTEFTPTEKDEYACRVNHVTLSQPKIVKWGKSYILL**GGGGSGGGGSGGGGS**GSH
SMRYFSTSVSRPGSGEPRFIAVGYVDDTQFVRFDSDAASQRMEPRAPWIEQERPEYWDQETRNVKAQSQT
DRVDLGTLRGYYNQSEAGSHTIQIMYGCDVGSDGRFLRGYEQHAYDGKDYIALNEDLRSWTAADMAAQIT
QRKWEAARWAEQLRAYLEGTCVEWLRRYLENGKETLQRTDPPKTHMTHHPISDHEATLRCWALGFYPAEI
TLTWQRDGEDQTQDTELVETRPAGDGTFQKWAAVVVPSGEEQRYTCHVQHEGLPKPLTLRWEPKSCDKTH
TCPPCPAPELLGGPSVFLFPPKPKDTLMISRTPEVTCVVVDVSHEDPEVKFNWYVDGVEVHNAKTKPREE
QYNSTYRVVSVLTVLHQDWLNGKEYKCKVSNKGLPSSIEKTISKAKGQPREPQVYTLPPSRDELTKNQVS
LTCLVKGFYPSDIAVEWESNGQPENNYKTTPPVLDSDGSFFLYSKLTVDKSRWQQGNVFSCSVMHEALHN
HYTQKSLSLSPGK**ITIFITLFLLSVCYSATVTFFKVKWIFSSVVDLKQTIIPDYRNMIGQGA**

> A*30:02P-Final

**MSRSVALAVLALLSLSGLEA**IQRTPKIQVYSRHPAENGKSNFLNCYVSGFHPSDIEVDLLKNGERIEKVE
HSDLSFSKDWSFYLLYYTEFTPTEKDEYACRVNHVTLSQPKIVKWGKSYILL**GGGGSGGGGSGGGGS**GSH
SMRYFSTSVSRPGSGEPRFIAVGYVDDTQFVRFDSDAASQRMEPRAPWIEQERPEYWDQETRNVKAHSQT
DRENLGTLRGYYNQSEAGSHTIQIMYGCDVGSDGRFLRGYEQHAYDGKDYIALNEDLRSWTAADMAAQIT
QRKWEAARRAEQLRAYLEGTCVEWLRRYLENGKETLQRTDPPKTHMTHHPISDHEATLRCWALGFYPAEI
TLTWQRDGEDQTQDTELVETRPAGDGTFQKWAAVVVPSGEEQRYTCHVQHEGLPKPLTLRWEPKSCDKTH
TCPPCPAPELLGGPSVFLFPPKPKDTLMISRTPEVTCVVVDVSHEDPEVKFNWYVDGVEVHNAKTKPREE
QYNSTYRVVSVLTVLHQDWLNGKEYKCKVSNKGLPSSIEKTISKAKGQPREPQVYTLPPSRDELTKNQVS
LTCLVKGFYPSDIAVEWESNGQPENNYKTTPPVLDSDGSFFLYSKLTVDKSRWQQGNVFSCSVMHEALHN
HYTQKSLSLSPGK**ITIFITLFLLSVCYSATVTFFKVKWIFSSVVDLKQTIIPDYRNMIGQGA**

> A*31:01P-Final

**MSRSVALAVLALLSLSGLEA**IQRTPKIQVYSRHPAENGKSNFLNCYVSGFHPSDIEVDLLKNGERIEKVE
HSDLSFSKDWSFYLLYYTEFTPTEKDEYACRVNHVTLSQPKIVKWGKSYILL**GGGGSGGGGSGGGGS**GSH
SMRYFTTSVSRPGRGEPRFIAVGYVDDTQFVRFDSDAASQRMEPRAPWIEQERPEYWDQETRNVKAHSQI
DRVDLGTLRGYYNQSEAGSHTIQMMYGCDVGSDGRFLRGYQQDAYDGKDYIALNEDLRSWTAADMAAQIT
QRKWEAARVAEQLRAYLEGTCVEWLRRYLENGKETLQRTDPPKTHMTHHAVSDHEATLRCWALSFYPAEI

TLTWQRDGEDQTQDTELVETRPAGDGTFQKWASVVVPSGQEQRYTCHVQHEGLPKPLTLRWEPKSCDKTH
TCPPCPAPELLGGPSVFLFPPKPKDTLMISRTPEVTCVVVDVSHEDPEVKFNWYVDGVEVHNAKTKPREE
QYNSTYRVVSVLTVLHQDWLNGKEYKCKVSNKGLPSSIEKTISKAKGQPREPQVYTLPPSRDELTKNQVS
LTCLVKGFYPSDIAVEWESNGQPENNYKTTPPVLDSDGSFFLYSKLTVDKSRWQQGNVFSCSVMHEALHN
HYTQKSLSLSPGK**ITIFITLFLLSVCYSATVTFFKVKWIFSSVVDLKQTIIPDYRNMIGQGA**

> A*32:01P-Final

**MSRSVALAVLALLSLSGLEA**IQRTPKIQVYSRHPAENGKSNFLNCYVSGFHPSDIEVDLLKNGERIEKVE
HSDLSFSKDWSFYLLYYTEFTPTEKDEYACRVNHVTLSQPKIVKWGKSYILL**GGGGSGGGGSGGGGS**GSH
SMRYFFTSVSRPGRGEPRFIAVGYVDDTQFVRFDSDAASQRMEPRAPWIEQEGPEYWDQETRNVKAHSQT
DRESLRIALRYYNQSEAGSHTIQMMYGCDVGPDGRLLRGYQQDAYDGKDYIALNEDLRSWTAADMAAQIT
QRKWEAARVAEQLRAYLEGTCVEWLRRYLENGKETLQRTDAPKTHMTHHAVSDHEATLRCWALSFYPAEI
TLTWQRDGEDQTQDTELVETRPAGDGTFQKWASVVVPSGQEQRYTCHVQHEGLPKPLTLRWEPKSCDKTH
TCPPCPAPELLGGPSVFLFPPKPKDTLMISRTPEVTCVVVDVSHEDPEVKFNWYVDGVEVHNAKTKPREE
QYNSTYRVVSVLTVLHQDWLNGKEYKCKVSNKGLPSSIEKTISKAKGQPREPQVYTLPPSRDELTKNQVS
LTCLVKGFYPSDIAVEWESNGQPENNYKTTPPVLDSDGSFFLYSKLTVDKSRWQQGNVFSCSVMHEALHN
HYTQKSLSLSPGK**ITIFITLFLLSVCYSATVTFFKVKWIFSSVVDLKQTIIPDYRNMIGQGA**

> A*33:01P-Final

**MSRSVALAVLALLSLSGLEA**IQRTPKIQVYSRHPAENGKSNFLNCYVSGFHPSDIEVDLLKNGERIEKVE
HSDLSFSKDWSFYLLYYTEFTPTEKDEYACRVNHVTLSQPKIVKWGKSYILL**GGGGSGGGGSGGGGS**GSH
SMRYFTTSVSRPGRGEPRFIAVGYVDDTQFVRFDSDAASQRMEPRAPWIEQEGPEYWDRNTRNVKAHSQI
DRVDLGTLRGYYNQSEAGSHTIQMMYGCDVGSDGRFLRGYQQDAYDGKDYIALNEDLRSWTAADMAAQIT
QRKWEAARVAEQLRAYLEGTCVEWLRRHLENGKETLQRTDPPRTHMTHHAVSDHEATLRCWALSFYPAEI
TLTWQRDGEDQTQDTELVETRPAGDGTFQKWASVVVPSGQEQRYTCHVQHEGLPKPLTLRWEPKSCDKTH
TCPPCPAPELLGGPSVFLFPPKPKDTLMISRTPEVTCVVVDVSHEDPEVKFNWYVDGVEVHNAKTKPREE
QYNSTYRVVSVLTVLHQDWLNGKEYKCKVSNKGLPSSIEKTISKAKGQPREPQVYTLPPSRDELTKNQVS
LTCLVKGFYPSDIAVEWESNGQPENNYKTTPPVLDSDGSFFLYSKLTVDKSRWQQGNVFSCSVMHEALHN
HYTQKSLSLSPGK**ITIFITLFLLSVCYSATVTFFKVKWIFSSVVDLKQTIIPDYRNMIGQGA**

> A*33:03P-Final

**MSRSVALAVLALLSLSGLEA**IQRTPKIQVYSRHPAENGKSNFLNCYVSGFHPSDIEVDLLKNGERIEKVE
HSDLSFSKDWSFYLLYYTEFTPTEKDEYACRVNHVTLSQPKIVKWGKSYILL**GGGGSGGGGSGGGGS**GSH
SMRYFTTSVSRPGRGEPRFIAVGYVDDTQFVRFDSDAASQRMEPRAPWIEQEGPEYWDRNTRNVKAHSQI
DRVDLGTLRGYYNQSEAGSHTIQMMYGCDVGSDGRFLRGYQQDAYDGKDYIALNEDLRSWTAADMAAQIT
QRKWEAARVAEQLRAYLEGTCVEWLRRYLENGKETLQRTDPPKTHMTHHAVSDHEATLRCWALSFYPAEI
TLTWQRDGEDQTQDTELVETRPAGDGTFQKWASVVVPSGQEQRYTCHVQHEGLPKPLTLRWEPKSCDKTH
TCPPCPAPELLGGPSVFLFPPKPKDTLMISRTPEVTCVVVDVSHEDPEVKFNWYVDGVEVHNAKTKPREE
QYNSTYRVVSVLTVLHQDWLNGKEYKCKVSNKGLPSSIEKTISKAKGQPREPQVYTLPPSRDELTKNQVS
LTCLVKGFYPSDIAVEWESNGQPENNYKTTPPVLDSDGSFFLYSKLTVDKSRWQQGNVFSCSVMHEALHN
HYTQKSLSLSPGK**ITIFITLFLLSVCYSATVTFFKVKWIFSSVVDLKQTIIPDYRNMIGQGA**

> A*34:01P-Final

**MSRSVALAVLALLSLSGLEA**IQRTPKIQVYSRHPAENGKSNFLNCYVSGFHPSDIEVDLLKNGERIEKVE
HSDLSFSKDWSFYLLYYTEFTPTEKDEYACRVNHVTLSQPKIVKWGKSYILL**GGGGSGGGGSGGGGS**GSH
SMRYFYTSVSRPGRGEPRFIAVGYVDDTQFVRFDSDAASQRMEPRAPWIEQEGPEYWDRNTRKVKAQSQT
DRVDLGTLRGYYNQSEDGSHTIQRMYGCDVGPDGRFLRGYQQDAYDGKDYIALNEDLRSWTAADMAAQIT
QRKWETAHEAEQWRAYLEGTCVEWLRRYLENGKETLQRTDAPKTHMTHHAVSDHEATLRCWALSFYPAEI
TLTWQRDGEDQTQDTELVETRPAGDGTFQKWASVVVPSGQEQRYTCHVQHEGLPKPLTLRWEPKSCDKTH
TCPPCPAPELLGGPSVFLFPPKPKDTLMISRTPEVTCVVVDVSHEDPEVKFNWYVDGVEVHNAKTKPREE
QYNSTYRVVSVLTVLHQDWLNGKEYKCKVSNKGLPSSIEKTISKAKGQPREPQVYTLPPSRDELTKNQVS
LTCLVKGFYPSDIAVEWESNGQPENNYKTTPPVLDSDGSFFLYSKLTVDKSRWQQGNVFSCSVMHEALHN
HYTQKSLSLSPGK**ITIFITLFLLSVCYSATVTFFKVKWIFSSVVDLKQTIIPDYRNMIGQGA**

> A*34:02P-Final

**MSRSVALAVLALLSLSGLEA**IQRTPKIQVYSRHPAENGKSNFLNCYVSGFHPSDIEVDLLKNGERIEKVE
HSDLSFSKDWSFYLLYYTEFTPTEKDEYACRVNHVTLSQPKIVKWGKSYILL**GGGGSGGGGSGGGGS**GSH
SMRYFYTSVSRPGRGEPRFIAVGYVDDTQFVRFDSDAASQRMEPRAPWIEQEGPEYWDRNTRNVKAQSQT
DRVDLGTLRGYYNQSEDGSHTIQIMYGCDVGSDGRFLRGYRQDAYDGKDYIALNEDLRSWTAADMAAQIT
QRKWETAHEAEQLRAYLEGTCVEWLRRYLENGKETLQRTDAPKTHMTHHAVSDHEATLRCWALSFYPAEI
TLTWQRDGEDQTQDTELVETRPAGDGTFQKWASVVVPSGQEQRYTCHVQHEGLPKPLTLRWEPKSCDKTH
TCPPCPAPELLGGPSVFLFPPKPKDTLMISRTPEVTCVVVDVSHEDPEVKFNWYVDGVEVHNAKTKPREE
QYNSTYRVVSVLTVLHQDWLNGKEYKCKVSNKGLPSSIEKTISKAKGQPREPQVYTLPPSRDELTKNQVS
LTCLVKGFYPSDIAVEWESNGQPENNYKTTPPVLDSDGSFFLYSKLTVDKSRWQQGNVFSCSVMHEALHN
HYTQKSLSLSPGK**ITIFITLFLLSVCYSATVTFFKVKWIFSSVVDLKQTIIPDYRNMIGQGA**

> A*36:01P-Final

**MSRSVALAVLALLSLSGLEA**IQRTPKIQVYSRHPAENGKSNFLNCYVSGFHPSDIEVDLLKNGERIEKVE
HSDLSFSKDWSFYLLYYTEFTPTEKDEYACRVNHVTLSQPKIVKWGKSYILL**GGGGSGGGGSGGGGS**GSH
SMRYFFTSVSRPGRGEPRFIAVGYVDDTQFVRFDSDAASQKMEPRAPWIEQEGPEYWDQETRNMKAHSQT
DRANLGTLRGYYNQSEDGSHTIQIMYGCDVGPDGRFLRGYRQDAYDGKDYIALNEDLRSWTAADMAAQIT
KRKWEAVHAAEQRRVYLEGTCVEWLRRYLENGKETLQRTDPPKTHMTHHPISDHEATLRCWALGFYPAEI
TLTWQRDGEDQTQDTELVETRPAGDGTFQKWAAVVVPSGEEQRYTCHVQHEGLPKPLTLRWEPKSCDKTH
TCPPCPAPELLGGPSVFLFPPKPKDTLMISRTPEVTCVVVDVSHEDPEVKFNWYVDGVEVHNAKTKPREE
QYNSTYRVVSVLTVLHQDWLNGKEYKCKVSNKGLPSSIEKTISKAKGQPREPQVYTLPPSRDELTKNQVS
LTCLVKGFYPSDIAVEWESNGQPENNYKTTPPVLDSDGSFFLYSKLTVDKSRWQQGNVFSCSVMHEALHN
HYTQKSLSLSPGK**ITIFITLFLLSVCYSATVTFFKVKWIFSSVVDLKQTIIPDYRNMIGQGA**

> A*43:01-Final

**MSRSVALAVLALLSLSGLEA**IQRTPKIQVYSRHPAENGKSNFLNCYVSGFHPSDIEVDLLKNGERIEKVE
HSDLSFSKDWSFYLLYYTEFTPTEKDEYACRVNHVTLSQPKIVKWGKSYILL**GGGGSGGGGSGGGGS**GSH
SMRYFYTSVSRPGRGEPRFIAVGYVDDTQFVRFDSDAASQRMEPRAPWIEQEGPEYWDLQTRNVKAHSQT
DRANLGTLRGYYNQSEDGSHTIQRMYGCDVGPDGRFLRGYQQDAYDGKDYIALNEDLRSWTAADMAAQIT
QRKWETAHEAEQWRAYLEGRCVEWLRRYLENGKETLQRTDAPKTHMTHHAVSDHEATLRCWALSFYPAEI
TLTWQRDGEDQTQDTELVETRPAGDGTFQKWASVVVPSGQEQRYTCHVQHEGLPKPLTLRWEPKSCDKTH

TCPPCPAPELLGGPSVFLFPPKPKDTLMISRTPEVTCVVVDVSHEDPEVKFNWYVDGVEVHNAKTKPREE
QYNSTYRVVSVLTVLHQDWLNGKEYKCKVSNKGLPSSIEKTISKAKGQPREPQVYTLPPSRDELTKNQVS
LTCLVKGFYPSDIAVEWESNGQPENNYKTTPPVLDSDGSFFLYSKLTVDKSRWQQGNVFSCSVMHEALHN
HYTQKSLSLSPGK**ITIFITLFLLSVCYSATVTFFKVKWIFSSVVDLKQTIIPDYRNMIGQGA**

> A*66:01P-Final

**MSRSVALAVLALLSLSGLEA**IQRTPKIQVYSRHPAENGKSNFLNCYVSGFHPSDIEVDLLKNGERIEKVE
HSDLSFSKDWSFYLLYYTEFTPTEKDEYACRVNHVTLSQPKIVKWGKSYILL**GGGGSGGGGSGGGGS**GSH
SMRYFYTSVSRPGRGEPRFIAVGYVDDTQFVRFDSDAASQRMEPRAPWIEQEGPEYWDRNTRNVKAQSQT
DRVDLGTLRGYYNQSEDGSHTIQRMYGCDVGPDGRFLRGYQQDAYDGKDYIALNEDLRSWTAADMAAQIT
QRKWETAHEAEQWRAYLEGRCVEWLRRYLENGKETLQRTDAPKTHMTHHAVSDHEATLRCWALSFYPAEI
TLTWQRDGEDQTQDTELVETRPAGDGTFQKWASVVVPSGQEQRYTCHVQHEGLPKPLTLRWEPKSCDKTH
TCPPCPAPELLGGPSVFLFPPKPKDTLMISRTPEVTCVVVDVSHEDPEVKFNWYVDGVEVHNAKTKPREE
QYNSTYRVVSVLTVLHQDWLNGKEYKCKVSNKGLPSSIEKTISKAKGQPREPQVYTLPPSRDELTKNQVS
LTCLVKGFYPSDIAVEWESNGQPENNYKTTPPVLDSDGSFFLYSKLTVDKSRWQQGNVFSCSVMHEALHN
HYTQKSLSLSPGK**ITIFITLFLLSVCYSATVTFFKVKWIFSSVVDLKQTIIPDYRNMIGQGA**

> A*66:02P-Final

**MSRSVALAVLALLSLSGLEA**IQRTPKIQVYSRHPAENGKSNFLNCYVSGFHPSDIEVDLLKNGERIEKVE
HSDLSFSKDWSFYLLYYTEFTPTEKDEYACRVNHVTLSQPKIVKWGKSYILL**GGGGSGGGGSGGGGS**GSH
SMRYFYTSVSRPGRGEPRFIAVGYVDDTQFVRFDSDAASQRMEPRAPWIEQEGPEYWDRNTRNVKAQSQT
DRVDLGTLRGYYNQSEAGSHTIQRMYGCDVGPDGRFLRGYQQDAYDGKDYIALNEDLRSWTAADMAAQIT
QRKWETAHEAEQWRAYLEGECVEWLRRYLENGKETLQRTDAPKTHMTHHAVSDHEATLRCWALSFYPAEI
TLTWQRDGEDQTQDTELVETRPAGDGTFQKWASVVVPSGQEQRYTCHVQHEGLPKPLTLRWEPKSCDKTH
TCPPCPAPELLGGPSVFLFPPKPKDTLMISRTPEVTCVVVDVSHEDPEVKFNWYVDGVEVHNAKTKPREE
QYNSTYRVVSVLTVLHQDWLNGKEYKCKVSNKGLPSSIEKTISKAKGQPREPQVYTLPPSRDELTKNQVS
LTCLVKGFYPSDIAVEWESNGQPENNYKTTPPVLDSDGSFFLYSKLTVDKSRWQQGNVFSCSVMHEALHN
HYTQKSLSLSPGK**ITIFITLFLLSVCYSATVTFFKVKWIFSSVVDLKQTIIPDYRNMIGQGA**

> A*68:01P-Final

**MSRSVALAVLALLSLSGLEA**IQRTPKIQVYSRHPAENGKSNFLNCYVSGFHPSDIEVDLLKNGERIEKVE
HSDLSFSKDWSFYLLYYTEFTPTEKDEYACRVNHVTLSQPKIVKWGKSYILL**GGGGSGGGGSGGGGS**GSH
SMRYFYTSVSRPGRGEPRFIAVGYVDDTQFVRFDSDAASQRMEPRAPWIEQEGPEYWDRNTRNVKAQSQT
DRVDLGTLRGYYNQSEAGSHTIQMMYGCDVGSDGRFLRGYRQDAYDGKDYIALKEDLRSWTAADMAAQTT
KHKWEAAHVAEQWRAYLEGTCVEWLRRYLENGKETLQRTDAPKTHMTHHAVSDHEATLRCWALSFYPAEI
TLTWQRDGEDQTQDTELVETRPAGDGTFQKWVAVVVPSGQEQRYTCHVQHEGLPKPLTLRWEPKSCDKTH
TCPPCPAPELLGGPSVFLFPPKPKDTLMISRTPEVTCVVVDVSHEDPEVKFNWYVDGVEVHNAKTKPREE
QYNSTYRVVSVLTVLHQDWLNGKEYKCKVSNKGLPSSIEKTISKAKGQPREPQVYTLPPSRDELTKNQVS

LTCLVKGFYPSDIAVEWESNGQPENNYKTTPPVLDSDGSFFLYSKLTVDKSRWQQGNVFSCSVMHEALHN
HYTQKSLSLSPGK**ITIFITLFLLSVCYSATVTFFKVKWIFSSVVDLKQTIIPDYRNMIGQGA**

> A*68:02P-Final

**MSRSVALAVLALLSLSGLEA**IQRTPKIQVYSRHPAENGKSNFLNCYVSGFHPSDIEVDLLKNGERIEKVE
HSDLSFSKDWSFYLLYYTEFTPTEKDEYACRVNHVTLSQPKIVKWGKSYILL**GGGGSGGGGSGGGGS**GSH
SMRYFYTSMSRPGRGEPRFIAVGYVDDTQFVRFDSDAASQRMEPRAPWIEQEGPEYWDRNTRNVKAQSQT
DRVDLGTLRGYYNQSEAGSHTIQRMYGCDVGPDGRFLRGYHQYAYDGKDYIALKEDLRSWTAADMAAQTT
KHKWEAAHVAEQWRAYLEGTCVEWLRRYLENGKETLQRTDAPKTHMTHHAVSDHEATLRCWALSFYPAEI
TLTWQRDGEDQTQDTELVETRPAGDGTFQKWVAVVVPSGQEQRYTCHVQHEGLPKPLTLRWEPKSCDKTH
TCPPCPAPELLGGPSVFLFPPKPKDTLMISRTPEVTCVVVDVSHEDPEVKFNWYVDGVEVHNAKTKPREE
QYNSTYRVVSVLTVLHQDWLNGKEYKCKVSNKGLPSSIEKTISKAKGQPREPQVYTLPPSRDELTKNQVS
LTCLVKGFYPSDIAVEWESNGQPENNYKTTPPVLDSDGSFFLYSKLTVDKSRWQQGNVFSCSVMHEALHN
HYTQKSLSLSPGK**ITIFITLFLLSVCYSATVTFFKVKWIFSSVVDLKQTIIPDYRNMIGQGA**

> A*69:01P-Final

**MSRSVALAVLALLSLSGLEA**IQRTPKIQVYSRHPAENGKSNFLNCYVSGFHPSDIEVDLLKNGERIEKVE
HSDLSFSKDWSFYLLYYTEFTPTEKDEYACRVNHVTLSQPKIVKWGKSYILL**GGGGSGGGGSGGGGS**GSH
SMRYFYTSVSRPGRGEPRFIAVGYVDDTQFVRFDSDAASQRMEPRAPWIEQEGPEYWDRNTRNVKAQSQT
DRVDLGTLRGYYNQSEAGSHTVQRMYGCDVGSDWRFLRGYHQYAYDGKDYIALKEDLRSWTAADMAAQTT
KHKWEAAHVAEQLRAYLEGTCVEWLRRYLENGKETLQRTDAPKTHMTHHAVSDHEATLRCWALSFYPAEI
TLTWQRDGEDQTQDTELVETRPAGDGTFQKWAAVVVPSGQEQRYTCHVQHEGLPKPLTLRWEPKSCDKTH
TCPPCPAPELLGGPSVFLFPPKPKDTLMISRTPEVTCVVVDVSHEDPEVKFNWYVDGVEVHNAKTKPREE
QYNSTYRVVSVLTVLHQDWLNGKEYKCKVSNKGLPSSIEKTISKAKGQPREPQVYTLPPSRDELTKNQVS
LTCLVKGFYPSDIAVEWESNGQPENNYKTTPPVLDSDGSFFLYSKLTVDKSRWQQGNVFSCSVMHEALHN
HYTQKSLSLSPGK**ITIFITLFLLSVCYSATVTFFKVKWIFSSVVDLKQTIIPDYRNMIGQGA**

> A*74:01P-Final

**MSRSVALAVLALLSLSGLEA**IQRTPKIQVYSRHPAENGKSNFLNCYVSGFHPSDIEVDLLKNGERIEKVE
HSDLSFSKDWSFYLLYYTEFTPTEKDEYACRVNHVTLSQPKIVKWGKSYILL**GGGGSGGGGSGGGGS**GSH
SMRYFFTSVSRPGRGEPRFIAVGYVDDTQFVRFDSDAASQRMEPRAPWIEQEGPEYWDQETRNVKAHSQT
DRVDLGTLRGYYNQSEAGSHTIQMMYGCDVGPDGRLLRGYQQDAYDGKDYIALNEDLRSWTAADMAAQIT
QRKWEAARVAEQLRAYLEGTCVEWLRRYLENGKETLQRTDAPKTHMTHHAVSDHEATLRCWALSFYPAEI
TLTWQRDGEDQTQDTELVETRPAGDGTFQKWASVVVPSGQEQRYTCHVQHEGLPKPLTLRWEPKSCDKTH
TCPPCPAPELLGGPSVFLFPPKPKDTLMISRTPEVTCVVVDVSHEDPEVKFNWYVDGVEVHNAKTKPREE
QYNSTYRVVSVLTVLHQDWLNGKEYKCKVSNKGLPSSIEKTISKAKGQPREPQVYTLPPSRDELTKNQVS
LTCLVKGFYPSDIAVEWESNGQPENNYKTTPPVLDSDGSFFLYSKLTVDKSRWQQGNVFSCSVMHEALHN
HYTQKSLSLSPGK**ITIFITLFLLSVCYSATVTFFKVKWIFSSVVDLKQTIIPDYRNMIGQGA**

> A*80:01P-Final

**MSRSVALAVLALLSLSGLEA**IQRTPKIQVYSRHPAENGKSNFLNCYVSGFHPSDIEVDLLKNGERIEKVE
HSDLSFSKDWSFYLLYYTEFTPTEKDEYACRVNHVTLSQPKIVKWGKSYILL**GGGGSGGGGSGGGGS**GSH

SMRYFFTSVSRPGRGEPRFIAVGYVDDSQFVQFDSDAASQRMEPRAPWIEQEEPEYWDEETRNVKAHSQT
NRANLGTLRGYYNQSEDGSHTIQIMYGCDVGSDGRFLRGYRQDAYDGKDYIALNEDLRSWTAADMAAQIT
KRKWEAARRAEQLRAYLEGECVDGLRRYLENGKETLQRTDPPKTHMTHHPISDHEATLRCWALSFYPAEI
TLTWQRDGEDQTQDTELVETRPAGDGTFQKWAAVVVPSGKEKRYTCHVQHEGLPEPLTLRWEPKSCDKTH
TCPPCPAPELLGGPSVFLFPPKPKDTLMISRTPEVTCVVVDVSHEDPEVKFNWYVDGVEVHNAKTKPREE
QYNSTYRVVSVLTVLHQDWLNGKEYKCKVSNKGLPSSIEKTISKAKGQPREPQVYTLPPSRDELTKNQVS
LTCLVKGFYPSDIAVEWESNGQPENNYKTTPPVLDSDGSFFLYSKLTVDKSRWQQGNVFSCSVMHEALHN
HYTQKSLSLSPGK**ITIFITLFLLSVCYSATVTFFKVKWIFSSVVDLKQTIIPDYRNMIGQGA**

HLA B

> B*07:02P-Final

**MSRSVALAVLALLSLSGLEA**IQRTPKIQVYSRHPAENGKSNFLNCYVSGFHPSDIEVDLLKNGERIEKVE
HSDLSFSKDWSFYLLYYTEFTPTEKDEYACRVNHVTLSQPKIVKWGKSYILL**GGGGSGGGGSGGGGS**GSH
SMRYFYTSVSRPGRGEPRFISVGYVDDTQFVRFDSDAASPREEPRAPWIEQEGPEYWDRNTQIYKAQAQT
DRESLRNLRGYYNQSEAGSHTLQSMYGCDVGPDGRLLRGHDQYAYDGKDYIALNEDLRSWTAADTAAQIT
QRKWEAAREAEQRRAYLEGECVEWLRRYLENGKDKLERADPPKTHVTHHPISDHEATLRCWALGFYPAEI
TLTWQRDGEDQTQDTELVETRPAGDRTFQKWAAVVVPSGEEQRYTCHVQHEGLPKPLTLRWEPKSCDKTH
TCPPCPAPELLGGPSVFLFPPKPKDTLMISRTPEVTCVVVDVSHEDPEVKFNWYVDGVEVHNAKTKPREE
QYNSTYRVVSVLTVLHQDWLNGKEYKCKVSNKGLPSSIEKTISKAKGQPREPQVYTLPPSRDELTKNQVS
LTCLVKGFYPSDIAVEWESNGQPENNYKTTPPVLDSDGSFFLYSKLTVDKSRWQQGNVFSCSVMHEALHN
HYTQKSLSLSPGK**ITIFITLFLLSVCYSATVTFFKVKWIFSSVVDLKQTIIPDYRNMIGQGA**

> B*08:01P-Final

**MSRSVALAVLALLSLSGLEA**IQRTPKIQVYSRHPAENGKSNFLNCYVSGFHPSDIEVDLLKNGERIEKVE
HSDLSFSKDWSFYLLYYTEFTPTEKDEYACRVNHVTLSQPKIVKWGKSYILL**GGGGSGGGGSGGGGS**GSH
SMRYFDTAMSRPGRGEPRFISVGYVDDTQFVRFDSDAASPREEPRAPWIEQEGPEYWDRNTQIFKTNTQT
DRESLRNLRGYYNQSEAGSHTLQSMYGCDVGPDGRLLRGHNQYAYDGKDYIALNEDLRSWTAADTAAQIT
QRKWEAARVAEQDRAYLEGTCVEWLRRYLENGKDTLERADPPKTHVTHHPISDHEATLRCWALGFYPAEI
TLTWQRDGEDQTQDTELVETRPAGDRTFQKWAAVVVPSGEEQRYTCHVQHEGLPKPLTLRWEPKSCDKTH
TCPPCPAPELLGGPSVFLFPPKPKDTLMISRTPEVTCVVVDVSHEDPEVKFNWYVDGVEVHNAKTKPREE
QYNSTYRVVSVLTVLHQDWLNGKEYKCKVSNKGLPSSIEKTISKAKGQPREPQVYTLPPSRDELTKNQVS
LTCLVKGFYPSDIAVEWESNGQPENNYKTTPPVLDSDGSFFLYSKLTVDKSRWQQGNVFSCSVMHEALHN
HYTQKSLSLSPGK**ITIFITLFLLSVCYSATVTFFKVKWIFSSVVDLKQTIIPDYRNMIGQGA**

> B*13:01P-Final

**MSRSVALAVLALLSLSGLEA**IQRTPKIQVYSRHPAENGKSNFLNCYVSGFHPSDIEVDLLKNGERIEKVE
HSDLSFSKDWSFYLLYYTEFTPTEKDEYACRVNHVTLSQPKIVKWGKSYILL**GGGGSGGGGSGGGGS**GSH
SMRYFYTAMSRPGRGEPRFITVGYVDDTQFVRFDSDATSPRMAPRAPWIEQEGPEYWDRETQISKTNTQT
YRENLRTALRYYNQSEAGSHIIQRMYGCDLGPDGRLLRGHNQLAYDGKDYIALNEDLSSWTAADTAAQIT
QLKWEAARVAEQLRAYLEGECVEWLRRYLENGKETLQRADPPKTHVTHHPISDHEATLRCWALGFYPAEI
TLTWQRDGEDQTQDTELVETRPAGDRTFQKWAAVVVPSGEEQRYTCHVQHEGLPKPLTLRWEPKSCDKTH
TCPPCPAPELLGGPSVFLFPPKPKDTLMISRTPEVTCVVVDVSHEDPEVKFNWYVDGVEVHNAKTKPREE

QYNSTYRVVSVLTVLHQDWLNGKEYKCKVSNKGLPSSIEKTISKAKGQPREPQVYTLPPSRDELTKNQVS
LTCLVKGFYPSDIAVEWESNGQPENNYKTTPPVLDSDGSFFLYSKLTVDKSRWQQGNVFSCSVMHEALHN
HYTQKSLSLSPGK**ITIFITLFLLSVCYSATVTFFKVKWIFSSVVDLKQTIIPDYRNMIGQGA**

> B*13:02P-Final

**MSRSVALAVLALLSLSGLEA**IQRTPKIQVYSRHPAENGKSNFLNCYVSGFHPSDIEVDLLKNGERIEKVE
HSDLSFSKDWSFYLLYYTEFTPTEKDEYACRVNHVTLSQPKIVKWGKSYILL**GGGGSGGGGSGGGGS**GSH
SMRYFYTAMSRPGRGEPRFITVGYVDDTQFVRFDSDATSPRMAPRAPWIEQEGPEYWDRETQISKTNTQT
YRENLRTALRYYNQSEAGSHTWQTMYGCDLGPDGRLLRGHNQLAYDGKDYIALNEDLSSWTAADTAAQIT
QLKWEAARVAEQLRAYLEGECVEWLRRYLENGKETLQRADPPKTHVTHHPISDHEATLRCWALGFYPAEI
TLTWQRDGEDQTQDTELVETRPAGDRTFQKWAAVVVPSGEEQRYTCHVQHEGLPKPLTLRWEPKSCDKTH
TCPPCPAPELLGGPSVFLFPPKPKDTLMISRTPEVTCVVVDVSHEDPEVKFNWYVDGVEVHNAKTKPREE
QYNSTYRVVSVLTVLHQDWLNGKEYKCKVSNKGLPSSIEKTISKAKGQPREPQVYTLPPSRDELTKNQVS
LTCLVKGFYPSDIAVEWESNGQPENNYKTTPPVLDSDGSFFLYSKLTVDKSRWQQGNVFSCSVMHEALHN
HYTQKSLSLSPGK**ITIFITLFLLSVCYSATVTFFKVKWIFSSVVDLKQTIIPDYRNMIGQGA**

>B*18:01P-Final

**MSRSVALAVLALLSLSGLEA**IQRTPKIQVYSRHPAENGKSNFLNCYVSGFHPSDIEVDLLKNGERIEKVE
HSDLSFSKDWSFYLLYYTEFTPTEKDEYACRVNHVTLSQPKIVKWGKSYILL**GGGGSGGGGSGGGGS**GSH
SMRYFHTSVSRPGRGEPRFISVGYVDGTQFVRFDSDAASPRTEPRAPWIEQEGPEYWDRNTQISKTNTQT
YRESLRNLRGYYNQSEAGSHTLQRMYGCDVGPDGRLLRGHDQSAYDGKDYIALNEDLSSWTAADTAAQIT
QRKWEAARVAEQLRAYLEGTCVEWLRRHLENGKETLQRADPPKTHVTHHPISDHEATLRCWALGFYPAEI
TLTWQRDGEDQTQDTELVETRPAGDRTFQKWAAVVVPSGEEQRYTCHVQHEGLPKPLTLRWEPKSCDKTH
TCPPCPAPELLGGPSVFLFPPKPKDTLMISRTPEVTCVVVDVSHEDPEVKFNWYVDGVEVHNAKTKPREE
QYNSTYRVVSVLTVLHQDWLNGKEYKCKVSNKGLPSSIEKTISKAKGQPREPQVYTLPPSRDELTKNQVS
LTCLVKGFYPSDIAVEWESNGQPENNYKTTPPVLDSDGSFFLYSKLTVDKSRWQQGNVFSCSVMHEALHN
HYTQKSLSLSPGK**ITIFITLFLLSVCYSATVTFFKVKWIFSSVVDLKQTIIPDYRNMIGQGA**

>B*27:05P-Final

**MSRSVALAVLALLSLSGLEA**IQRTPKIQVYSRHPAENGKSNFLNCYVSGFHPSDIEVDLLKNGERIEKVE
HSDLSFSKDWSFYLLYYTEFTPTEKDEYACRVNHVTLSQPKIVKWGKSYILL**GGGGSGGGGSGGGGS**GSH
SMRYFHTSVSRPGRGEPRFITVGYVDDTLFVRFDSDAASPREEPRAPWIEQEGPEYWDRETQICKAKAQT
DREDLRTLLRYYNQSEAGSHTLQNMYGCDVGPDGRLLRGYHQDAYDGKDYIALNEDLSSWTAADTAAQIT
QRKWEAARVAEQLRAYLEGECVEWLRRYLENGKETLQRADPPKTHVTHHPISDHEATLRCWALGFYPAEI
TLTWQRDGEDQTQDTELVETRPAGDRTFQKWAAVVVPSGEEQRYTCHVQHEGLPKPLTLRWEPKSCDKTH
TCPPCPAPELLGGPSVFLFPPKPKDTLMISRTPEVTCVVVDVSHEDPEVKFNWYVDGVEVHNAKTKPREE
QYNSTYRVVSVLTVLHQDWLNGKEYKCKVSNKGLPSSIEKTISKAKGQPREPQVYTLPPSRDELTKNQVS
LTCLVKGFYPSDIAVEWESNGQPENNYKTTPPVLDSDGSFFLYSKLTVDKSRWQQGNVFSCSVMHEALHN
HYTQKSLSLSPGK**ITIFITLFLLSVCYSATVTFFKVKWIFSSVVDLKQTIIPDYRNMIGQGA**

>B*27:08-Final

**MSRSVALAVLALLSLSGLEA**IQRTPKIQVYSRHPAENGKSNFLNCYVSGFHPSDIEVDLLKNGERIEKVE
HSDLSFSKDWSFYLLYYTEFTPTEKDEYACRVNHVTLSQPKIVKWGKSYILL**GGGGSGGGGSGGGGS**GSH
SMRYFHTSVSRPGRGEPRFITVGYVDDTLFVRFDSDAASPREEPRAPWIEQEGPEYWDRETQICKAKAQT
DRESLRNLRGYYNQSEAGSHTLQNMYGCDVGPDGRLLRGYHQDAYDGKDYIALNEDLSSWTAADTAAQIT
QRKWEAARVAEQLRAYLEGECVEWLRRYLENGKETLQRADPPKTHVTHHPISDHEATLRCWALGFYPAEI
TLTWQRDGEDQTQDTELVETRPAGDRTFQKWAAVVVPSGEEQRYTCHVQHEGLPKPLTLRWEPKSCDKTH
TCPPCPAPELLGGPSVFLFPPKPKDTLMISRTPEVTCVVVDVSHEDPEVKFNWYVDGVEVHNAKTKPREE
QYNSTYRVVSVLTVLHQDWLNGKEYKCKVSNKGLPSSIEKTISKAKGQPREPQVYTLPPSRDELTKNQVS
LTCLVKGFYPSDIAVEWESNGQPENNYKTTPPVLDSDGSFFLYSKLTVDKSRWQQGNVFSCSVMHEALHN
HYTQKSLSLSPGK**ITIFITLFLLSVCYSATVTFFKVKWIFSSVVDLKQTIIPDYRNMIGQGA**

>B*35:01-Final

**MSRSVALAVLALLSLSGLEA**IQRTPKIQVYSRHPAENGKSNFLNCYVSGFHPSDIEVDLLKNGERIEKVE
HSDLSFSKDWSFYLLYYTEFTPTEKDEYACRVNHVTLSQPKIVKWGKSYILL**GGGGSGGGGSGGGGS**GSH
SMRYFYTAMSRPGRGEPRFIAVGYVDDTQFVRFDSDAASPRTEPRAPWIEQEGPEYWDRNTQIFKTNTQT
YRESLRNLRGYYNQSEAGSHIIQRMYGCDLGPDGRLLRGHDQSAYDGKDYIALNEDLSSWTAADTAAQIT
QRKWEAARVAEQLRAYLEGLCVEWLRRYLENGKETLQRADPPKTHVTHHPVSDHEATLRCWALGFYPAEI
TLTWQRDGEDQTQDTELVETRPAGDRTFQKWAAVVVPSGEEQRYTCHVQHEGLPKPLTLRWEPKSCDKTH
TCPPCPAPELLGGPSVFLFPPKPKDTLMISRTPEVTCVVVDVSHEDPEVKFNWYVDGVEVHNAKTKPREE
QYNSTYRVVSVLTVLHQDWLNGKEYKCKVSNKGLPSSIEKTISKAKGQPREPQVYTLPPSRDELTKNQVS
LTCLVKGFYPSDIAVEWESNGQPENNYKTTPPVLDSDGSFFLYSKLTVDKSRWQQGNVFSCSVMHEALHN
HYTQKSLSLSPGK**ITIFITLFLLSVCYSATVTFFKVKWIFSSVVDLKQTIIPDYRNMIGQGA**

>B*37:01-Final

**MSRSVALAVLALLSLSGLEA**IQRTPKIQVYSRHPAENGKSNFLNCYVSGFHPSDIEVDLLKNGERIEKVE
HSDLSFSKDWSFYLLYYTEFTPTEKDEYACRVNHVTLSQPKIVKWGKSYILL**GGGGSGGGGSGGGGS**GSH
SMRYFHTSVSRPGRGEPRFISVGYVDDTQFVRFDSDAASPRTEPRAPWIEQEGPEYWDRETQISKTNTQT
YREDLRTLLRYYNQSEAGSHTIQRMSGCDVGPDGRLLRGYNQFAYDGKDYIALNEDLSSWTAADTAAQIT
QRKWEAARVAEQDRAYLEGTCVEWLRRYLENGKETLQRADPPKTHVTHHPISDHEATLRCWALGFYPAEI
TLTWQRDGEDQTQDTELVETRPAGDRTFQKWAAVVVPSGEEQRYTCHVQHEGLPKPLTLRWEPKSCDKTH
TCPPCPAPELLGGPSVFLFPPKPKDTLMISRTPEVTCVVVDVSHEDPEVKFNWYVDGVEVHNAKTKPREE
QYNSTYRVVSVLTVLHQDWLNGKEYKCKVSNKGLPSSIEKTISKAKGQPREPQVYTLPPSRDELTKNQVS
LTCLVKGFYPSDIAVEWESNGQPENNYKTTPPVLDSDGSFFLYSKLTVDKSRWQQGNVFSCSVMHEALHN
HYTQKSLSLSPGK**ITIFITLFLLSVCYSATVTFFKVKWIFSSVVDLKQTIIPDYRNMIGQGA**

>B*38:01-Final

**MSRSVALAVLALLSLSGLEA**IQRTPKIQVYSRHPAENGKSNFLNCYVSGFHPSDIEVDLLKNGERIEKVE
HSDLSFSKDWSFYLLYYTEFTPTEKDEYACRVNHVTLSQPKIVKWGKSYILL**GGGGSGGGGSGGGGS**GSH
SMRYFYTSVSRPGRGEPRFISVGYVDDTQFVRFDSDAASPREEPRAPWIEQEGPEYWDRNTQICKTNTQT
YRENLRIALRYYNQSEAGSHTLQRMYGCDVGPDGRLLRGHNQFAYDGKDYIALNEDLSSWTAADTAAQIT
QRKWEAARVAEQLRTYLEGTCVEWLRRYLENGKETLQRADPPKTHVTHHPISDHEATLRCWALGFYPAEI
TLTWQRDGEDQTQDTELVETRPAGDRTFQKWAAVVVPSGEEQRYTCHVQHEGLPKPLTLRWEPKSCDKTH

TCPPCPAPELLGGPSVFLFPPKPKDTLMISRTPEVTCVVVDVSHEDPEVKFNWYVDGVEVHNAKTKPREE
QYNSTYRVVSVLTVLHQDWLNGKEYKCKVSNKGLPSSIEKTISKAKGQPREPQVYTLPPSRDELTKNQVS
LTCLVKGFYPSDIAVEWESNGQPENNYKTTPPVLDSDGSFFLYSKLTVDKSRWQQGNVFSCSVMHEALHN
HYTQKSLSLSPGK**ITIFITLFLLSVCYSATVTFFKVKWIFSSVVDLKQTIIPDYRNMIGQGA**

>B*39:01-Final

**MSRSVALAVLALLSLSGLEA**IQRTPKIQVYSRHPAENGKSNFLNCYVSGFHPSDIEVDLLKNGERIEKVE
HSDLSFSKDWSFYLLYYTEFTPTEKDEYACRVNHVTLSQPKIVKWGKSYILL**GGGGSGGGGSGGGGS**GSH
SMRYFYTSVSRPGRGEPRFISVGYVDDTQFVRFDSDAASPREEPRAPWIEQEGPEYWDRNTQICKTNTQT
DRESLRNLRGYYNQSEAGSHTLQRMYGCDVGPDGRLLRGHNQFAYDGKDYIALNEDLSSWTAADTAAQIT
QRKWEAARVAEQLRTYLEGTCVEWLRRYLENGKETLQRADPPKTHVTHHPISDHEATLRCWALGFYPAEI
TLTWQRDGEDQTQDTELVETRPAGDRTFQKWAAVVVPSGEEQRYTCHVQHEGLPKPLTLRWEPKSCDKTH
TCPPCPAPELLGGPSVFLFPPKPKDTLMISRTPEVTCVVVDVSHEDPEVKFNWYVDGVEVHNAKTKPREE
QYNSTYRVVSVLTVLHQDWLNGKEYKCKVSNKGLPSSIEKTISKAKGQPREPQVYTLPPSRDELTKNQVS
LTCLVKGFYPSDIAVEWESNGQPENNYKTTPPVLDSDGSFFLYSKLTVDKSRWQQGNVFSCSVMHEALHN
HYTQKSLSLSPGK**ITIFITLFLLSVCYSATVTFFKVKWIFSSVVDLKQTIIPDYRNMIGQGA**

>B*41:01-Final

**MSRSVALAVLALLSLSGLEA**IQRTPKIQVYSRHPAENGKSNFLNCYVSGFHPSDIEVDLLKNGERIEKVE
HSDLSFSKDWSFYLLYYTEFTPTEKDEYACRVNHVTLSQPKIVKWGKSYILL**GGGGSGGGGSGGGGS**GSH
SMRYFHTAMSRPGRGEPRFITVGYVDDTLFVRFDSDATSPRKEPRAPWIEQEGPEYWDRETQISKTNTQT
YRESLRNLRGYYNQSEAGSHTWQRMYGCDVGPDGRLLRGHNQYAYDGKDYIALNEDLRSWTAADTAAQIT
QRKWEAARVAEQDRAYLEGTCVEWLRRYLENGKDTLERADPPKTHVTHHPISDHEATLRCWALGFYPAEI
TLTWQRDGEDQTQDTELVETRPAGDRTFQKWAAVVVPSGEEQRYTCHVQHEGLPKPLTLRWEPKSCDKTH
TCPPCPAPELLGGPSVFLFPPKPKDTLMISRTPEVTCVVVDVSHEDPEVKFNWYVDGVEVHNAKTKPREE
QYNSTYRVVSVLTVLHQDWLNGKEYKCKVSNKGLPSSIEKTISKAKGQPREPQVYTLPPSRDELTKNQVS
LTCLVKGFYPSDIAVEWESNGQPENNYKTTPPVLDSDGSFFLYSKLTVDKSRWQQGNVFSCSVMHEALHN
HYTQKSLSLSPGK**ITIFITLFLLSVCYSATVTFFKVKWIFSSVVDLKQTIIPDYRNMIGQGA**

>B*42:01-Final

**MSRSVALAVLALLSLSGLEA**IQRTPKIQVYSRHPAENGKSNFLNCYVSGFHPSDIEVDLLKNGERIEKVE
HSDLSFSKDWSFYLLYYTEFTPTEKDEYACRVNHVTLSQPKIVKWGKSYILL**GGGGSGGGGSGGGGS**GSH
SMRYFYTSVSRPGRGEPRFISVGYVDDTQFVRFDSDAASPREEPRAPWIEQEGPEYWDRNTQIYKAQAQT
DRESLRNLRGYYNQSEAGSHTLQSMYGCDVGPDGRLLRGHNQYAYDGKDYIALNEDLRSWTAADTAAQIT
QRKWEAARVAEQDRAYLEGTCVEWLRRYLENGKDTLERADPPKTHVTHHPISDHEATLRCWALGFYPAEI
TLTWQRDGEDQTQDTELVETRPAGDRTFQKWAAVVVPSGEEQRYTCHVQHEGLPKPLTLRWEPKSCDKTH
TCPPCPAPELLGGPSVFLFPPKPKDTLMISRTPEVTCVVVDVSHEDPEVKFNWYVDGVEVHNAKTKPREE
QYNSTYRVVSVLTVLHQDWLNGKEYKCKVSNKGLPSSIEKTISKAKGQPREPQVYTLPPSRDELTKNQVS
LTCLVKGFYPSDIAVEWESNGQPENNYKTTPPVLDSDGSFFLYSKLTVDKSRWQQGNVFSCSVMHEALHN
HYTQKSLSLSPGK**ITIFITLFLLSVCYSATVTFFKVKWIFSSVVDLKQTIIPDYRNMIGQGA**

>B*44:02-Final

**MSRSVALAVLALLSLSGLEA**IQRTPKIQVYSRHPAENGKSNFLNCYVSGFHPSDIEVDLLKNGERIEKVE
HSDLSFSKDWSFYLLYYTEFTPTEKDEYACRVNHVTLSQPKIVKWGKSYILL**GGGGSGGGGSGGGGS**GSH
SMRYFYTAMSRPGRGEPRFITVGYVDDTLFVRFDSDATSPRKEPRAPWIEQEGPEYWDRETQISKTNTQT
YRENLRTALRYYNQSEAGSHIIQRMYGCDVGPDGRLLRGYDQDAYDGKDYIALNEDLSSWTAADTAAQIT
QRKWEAARVAEQDRAYLEGLCVESLRRYLENGKETLQRADPPKTHVTHHPISDHEVTLRCWALGFYPAEI
TLTWQRDGEDQTQDTELVETRPAGDRTFQKWAAVVVPSGEEQRYTCHVQHEGLPKPLTLRWEPKSCDKTH
TCPPCPAPELLGGPSVFLFPPKPKDTLMISRTPEVTCVVVDVSHEDPEVKFNWYVDGVEVHNAKTKPREE
QYNSTYRVVSVLTVLHQDWLNGKEYKCKVSNKGLPSSIEKTISKAKGQPREPQVYTLPPSRDELTKNQVS
LTCLVKGFYPSDIAVEWESNGQPENNYKTTPPVLDSDGSFFLYSKLTVDKSRWQQGNVFSCSVMHEALHN
HYTQKSLSLSPGK**ITIFITLFLLSVCYSATVTFFKVKWIFSSVVDLKQTIIPDYRNMIGQGA**

>B*44:03-Final

**MSRSVALAVLALLSLSGLEA**IQRTPKIQVYSRHPAENGKSNFLNCYVSGFHPSDIEVDLLKNGERIEKVE
HSDLSFSKDWSFYLLYYTEFTPTEKDEYACRVNHVTLSQPKIVKWGKSYILL**GGGGSGGGGSGGGGS**GSH
SMRYFYTAMSRPGRGEPRFITVGYVDDTLFVRFDSDATSPRKEPRAPWIEQEGPEYWDRETQISKTNTQT
YRENLRTALRYYNQSEAGSHIIQRMYGCDVGPDGRLLRGYDQDAYDGKDYIALNEDLSSWTAADTAAQIT
QRKWEAARVAEQLRAYLEGLCVESLRRYLENGKETLQRADPPKTHVTHHPISDHEVTLRCWALGFYPAEI
TLTWQRDGEDQTQDTELVETRPAGDRTFQKWAAVVVPSGEEQRYTCHVQHEGLPKPLTLRWEPKSCDKTH
TCPPCPAPELLGGPSVFLFPPKPKDTLMISRTPEVTCVVVDVSHEDPEVKFNWYVDGVEVHNAKTKPREE
QYNSTYRVVSVLTVLHQDWLNGKEYKCKVSNKGLPSSIEKTISKAKGQPREPQVYTLPPSRDELTKNQVS
LTCLVKGFYPSDIAVEWESNGQPENNYKTTPPVLDSDGSFFLYSKLTVDKSRWQQGNVFSCSVMHEALHN
HYTQKSLSLSPGK**ITIFITLFLLSVCYSATVTFFKVKWIFSSVVDLKQTIIPDYRNMIGQGA**

>B*45:01-Final

**MSRSVALAVLALLSLSGLEA**IQRTPKIQVYSRHPAENGKSNFLNCYVSGFHPSDIEVDLLKNGERIEKVE
HSDLSFSKDWSFYLLYYTEFTPTEKDEYACRVNHVTLSQPKIVKWGKSYILL**GGGGSGGGGSGGGGS**GSH
SMRYFHTAMSRPGRGEPRFITVGYVDDTLFVRFDSDATSPRKEPRAPWIEQEGPEYWDRETQISKTNTQT
YRESLRNLRGYYNQSEAGSHTWQRMYGCDLGPDGRLLRGYNQLAYDGKDYIALNEDLSSWTAADTAAQIT
QRKWEAARVAEQDRAYLEGLCVESLRRYLENGKETLQRADPPKTHVTHHPISDHEATLRCWALGFYPAEI
TLTWQRDGEDQTQDTELVETRPAGDRTFQKWAAVVVPSGEEQRYTCHVQHEGLPKPLTLRWEPKSCDKTH
TCPPCPAPELLGGPSVFLFPPKPKDTLMISRTPEVTCVVVDVSHEDPEVKFNWYVDGVEVHNAKTKPREE
QYNSTYRVVSVLTVLHQDWLNGKEYKCKVSNKGLPSSIEKTISKAKGQPREPQVYTLPPSRDELTKNQVS
LTCLVKGFYPSDIAVEWESNGQPENNYKTTPPVLDSDGSFFLYSKLTVDKSRWQQGNVFSCSVMHEALHN
HYTQKSLSLSPGK**ITIFITLFLLSVCYSATVTFFKVKWIFSSVVDLKQTIIPDYRNMIGQGA**

>B*46:01-Final

**MSRSVALAVLALLSLSGLEA**IQRTPKIQVYSRHPAENGKSNFLNCYVSGFHPSDIEVDLLKNGERIEKVE
HSDLSFSKDWSFYLLYYTEFTPTEKDEYACRVNHVTLSQPKIVKWGKSYILL**GGGGSGGGGSGGGGS**GSH
SMRYFYTAMSRPGRGEPRFIAVGYVDDTQFVRFDSDAASPRMAPRAPWIEQEGPEYWDRETQKYKRQAQT
DRVSLRNLRGYYNQSEAGSHTLQRMYGCDVGPDGRLLRGHDQSAYDGKDYIALNEDLSSWTAADTAAQIT
QRKWEAAREAEQWRAYLEGLCVEWLRRYLENGKETLQRADPPKTHVTHHPISDHEATLRCWALGFYPAEI
TLTWQRDGEDQTQDTELVETRPAGDRTFQKWAAVVVPSGEEQRYTCHVQHEGLPKPLTLRWEPKSCDKTH

TCPPCPAPELLGGPSVFLFPPKPKDTLMISRTPEVTCVVVDVSHEDPEVKFNWYVDGVEVHNAKTKPREE
QYNSTYRVVSVLTVLHQDWLNGKEYKCKVSNKGLPSSIEKTISKAKGQPREPQVYTLPPSRDELTKNQVS
LTCLVKGFYPSDIAVEWESNGQPENNYKTTPPVLDSDGSFFLYSKLTVDKSRWQQGNVFSCSVMHEALHN
HYTQKSLSLSPGK**ITIFITLFLLSVCYSATVTFFKVKWIFSSVVDLKQTIIPDYRNMIGQGA**

>B*48:01-Final

**MSRSVALAVLALLSLSGLEA**IQRTPKIQVYSRHPAENGKSNFLNCYVSGFHPSDIEVDLLKNGERIEKVE
HSDLSFSKDWSFYLLYYTEFTPTEKDEYACRVNHVTLSQPKIVKWGKSYILL**GGGGSGGGGSGGGGS**GSH
SMRYFYTSVSRPGRGEPRFISVGYVDDTQFVRFDSDAASPREEPRAPWIEQEGPEYWDRETQISKTNTQT
YRESLRNLRGYYNQSEAGSHTLQSMYGCDVGPDGRLLRGHNQYAYDGKDYIALNEDLRSWTAADTAAQIS
QRKLEAARVAEQLRAYLEGECVEWLRRYLENGKDKLERADPPKTHVTHHPISDHEATLRCWALGFYPAEI
TLTWQRDGEDQTQDTELVETRPAGDRTFQKWTAVVVPSGEEQRYTCHVQHEGLPKPLTLRWEPKSCDKTH
TCPPCPAPELLGGPSVFLFPPKPKDTLMISRTPEVTCVVVDVSHEDPEVKFNWYVDGVEVHNAKTKPREE
QYNSTYRVVSVLTVLHQDWLNGKEYKCKVSNKGLPSSIEKTISKAKGQPREPQVYTLPPSRDELTKNQVS
LTCLVKGFYPSDIAVEWESNGQPENNYKTTPPVLDSDGSFFLYSKLTVDKSRWQQGNVFSCSVMHEALHN
HYTQKSLSLSPGK**ITIFITLFLLSVCYSATVTFFKVKWIFSSVVDLKQTIIPDYRNMIGQGA**

>B*49:01-Final

**MSRSVALAVLALLSLSGLEA**IQRTPKIQVYSRHPAENGKSNFLNCYVSGFHPSDIEVDLLKNGERIEKVE
HSDLSFSKDWSFYLLYYTEFTPTEKDEYACRVNHVTLSQPKIVKWGKSYILL**GGGGSGGGGSGGGGS**GSH
SMRYFHTAMSRPGRGEPRFITVGYVDDTLFVRFDSDATSPRKEPRAPWIEQEGPEYWDRETQISKTNTQT
YRENLRIALRYYNQSEAGSHTWQRMYGCDLGPDGRLLRGYNQLAYDGKDYIALNEDLSSWTAADTAAQIT
QRKWEAAREAEQLRAYLEGLCVEWLRRYLENGKETLQRADPPKTHVTHHPISDHEATLRCWALGFYPAEI
TLTWQRDGEDQTQDTELVETRPAGDRTFQKWAAVVVPSGEEQRYTCHVQHEGLPKPLTLRWEPKSCDKTH
TCPPCPAPELLGGPSVFLFPPKPKDTLMISRTPEVTCVVVDVSHEDPEVKFNWYVDGVEVHNAKTKPREE
QYNSTYRVVSVLTVLHQDWLNGKEYKCKVSNKGLPSSIEKTISKAKGQPREPQVYTLPPSRDELTKNQVS
LTCLVKGFYPSDIAVEWESNGQPENNYKTTPPVLDSDGSFFLYSKLTVDKSRWQQGNVFSCSVMHEALHN
HYTQKSLSLSPGK**ITIFITLFLLSVCYSATVTFFKVKWIFSSVVDLKQTIIPDYRNMIGQGA**

>B*50:01-Final

**MSRSVALAVLALLSLSGLEA**IQRTPKIQVYSRHPAENGKSNFLNCYVSGFHPSDIEVDLLKNGERIEKVE
HSDLSFSKDWSFYLLYYTEFTPTEKDEYACRVNHVTLSQPKIVKWGKSYILL**GGGGSGGGGSGGGGS**GSH
SMRYFHTAMSRPGRGEPRFITVGYVDDTLFVRFDSDATSPRKEPRAPWIEQEGPEYWDRETQISKTNTQT
YRESLRNLRGYYNQSEAGSHTWQRMYGCDLGPDGRLLRGYNQLAYDGKDYIALNEDLSSWTAADTAAQIT
QRKWEAAREAEQLRAYLEGLCVEWLRRYLENGKETLQRADPPKTHVTHHPISDHEATLRCWALGFYPAEI
TLTWQRDGEDQTQDTELVETRPAGDRTFQKWAAVVVPSGEEQRYTCHVQHEGLPKPLTLRWEPKSCDKTH
TCPPCPAPELLGGPSVFLFPPKPKDTLMISRTPEVTCVVVDVSHEDPEVKFNWYVDGVEVHNAKTKPREE
QYNSTYRVVSVLTVLHQDWLNGKEYKCKVSNKGLPSSIEKTISKAKGQPREPQVYTLPPSRDELTKNQVS
LTCLVKGFYPSDIAVEWESNGQPENNYKTTPPVLDSDGSFFLYSKLTVDKSRWQQGNVFSCSVMHEALHN
HYTQKSLSLSPGK**ITIFITLFLLSVCYSATVTFFKVKWIFSSVVDLKQTIIPDYRNMIGQGA**

>B*51:01-Final

**MSRSVALAVLALLSLSGLEA**IQRTPKIQVYSRHPAENGKSNFLNCYVSGFHPSDIEVDLLKNGERIEKVE
HSDLSFSKDWSFYLLYYTEFTPTEKDEYACRVNHVTLSQPKIVKWGKSYILL**GGGGSGGGGSGGGGS**GSH
SMRYFYTAMSRPGRGEPRFIAVGYVDDTQFVRFDSDAASPRTEPRAPWIEQEGPEYWDRNTQIFKTNTQT
YRENLRIALRYYNQSEAGSHTWQTMYGCDVGPDGRLLRGHNQYAYDGKDYIALNEDLSSWTAADTAAQIT
QRKWEAAREAEQLRAYLEGLCVEWLRRHLENGKETLQRADPPKTHVTHHPVSDHEATLRCWALGFYPAEI
TLTWQRDGEDQTQDTELVETRPAGDRTFQKWAAVVVPSGEEQRYTCHVQHEGLPKPLTLRWEPKSCDKTH
TCPPCPAPELLGGPSVFLFPPKPKDTLMISRTPEVTCVVVDVSHEDPEVKFNWYVDGVEVHNAKTKPREE
QYNSTYRVVSVLTVLHQDWLNGKEYKCKVSNKGLPSSIEKTISKAKGQPREPQVYTLPPSRDELTKNQVS
LTCLVKGFYPSDIAVEWESNGQPENNYKTTPPVLDSDGSFFLYSKLTVDKSRWQQGNVFSCSVMHEALHN
HYTQKSLSLSPGK**ITIFITLFLLSVCYSATVTFFKVKWIFSSVVDLKQTIIPDYRNMIGQGA**

>B*51:02-Final

**MSRSVALAVLALLSLSGLEA**IQRTPKIQVYSRHPAENGKSNFLNCYVSGFHPSDIEVDLLKNGERIEKVE
HSDLSFSKDWSFYLLYYTEFTPTEKDEYACRVNHVTLSQPKIVKWGKSYILL**GGGGSGGGGSGGGGS**GSH
SMRYFYTAMSRPGRGEPRFIAVGYVDDTQFVRFDSDAASPRTEPRAPWIEQEGPEYWDRNTQIFKTNTQT
YRENLRIALRYYNQSEAGSHTWQTMYGCDVGPDGRLLRGHNQYAYDGKDYIALNEDLSSWTAADTAAQIT
QRKWEAAREAEQLRAYLEGLCVEWLRRYLENGKETLQRADPPKTHVTHHPVSDHEATLRCWALGFYPAEI
TLTWQRDGEDQTQDTELVETRPAGDRTFQKWAAVVVPSGEEQRYTCHVQHEGLPKPLTLRWEPKSCDKTH
TCPPCPAPELLGGPSVFLFPPKPKDTLMISRTPEVTCVVVDVSHEDPEVKFNWYVDGVEVHNAKTKPREE
QYNSTYRVVSVLTVLHQDWLNGKEYKCKVSNKGLPSSIEKTISKAKGQPREPQVYTLPPSRDELTKNQVS
LTCLVKGFYPSDIAVEWESNGQPENNYKTTPPVLDSDGSFFLYSKLTVDKSRWQQGNVFSCSVMHEALHN
HYTQKSLSLSPGK**ITIFITLFLLSVCYSATVTFFKVKWIFSSVVDLKQTIIPDYRNMIGQGA**

>B*52:01-Final

**MSRSVALAVLALLSLSGLEA**IQRTPKIQVYSRHPAENGKSNFLNCYVSGFHPSDIEVDLLKNGERIEKVE
HSDLSFSKDWSFYLLYYTEFTPTEKDEYACRVNHVTLSQPKIVKWGKSYILL**GGGGSGGGGSGGGGS**GSH
SMRYFYTAMSRPGRGEPRFIAVGYVDDTQFVRFDSDAASPRTEPRAPWIEQEGPEYWDRETQISKTNTQT
YRENLRIALRYYNQSEAGSHTWQTMYGCDVGPDGRLLRGHNQYAYDGKDYIALNEDLSSWTAADTAAQIT
QRKWEAAREAEQLRAYLEGLCVEWLRRHLENGKETLQRADPPKTHVTHHPVSDHEATLRCWALGFYPAEI
TLTWQRDGEDQTQDTELVETRPAGDRTFQKWAAVVVPSGEEQRYTCHVQHEGLPKPLTLRWEPKSCDKTH
TCPPCPAPELLGGPSVFLFPPKPKDTLMISRTPEVTCVVVDVSHEDPEVKFNWYVDGVEVHNAKTKPREE
QYNSTYRVVSVLTVLHQDWLNGKEYKCKVSNKGLPSSIEKTISKAKGQPREPQVYTLPPSRDELTKNQVS
LTCLVKGFYPSDIAVEWESNGQPENNYKTTPPVLDSDGSFFLYSKLTVDKSRWQQGNVFSCSVMHEALHN
HYTQKSLSLSPGK**ITIFITLFLLSVCYSATVTFFKVKWIFSSVVDLKQTIIPDYRNMIGQGA**

>B*53:01-Final

**MSRSVALAVLALLSLSGLEA**IQRTPKIQVYSRHPAENGKSNFLNCYVSGFHPSDIEVDLLKNGERIEKVE
HSDLSFSKDWSFYLLYYTEFTPTEKDEYACRVNHVTLSQPKIVKWGKSYILL**GGGGSGGGGSGGGGS**GSH
SMRYFYTAMSRPGRGEPRFIAVGYVDDTQFVRFDSDAASPRTEPRAPWIEQEGPEYWDRNTQIFKTNTQT
YRENLRIALRYYNQSEAGSHIIQRMYGCDLGPDGRLLRGHDQSAYDGKDYIALNEDLSSWTAADTAAQIT
QRKWEAARVAEQLRAYLEGLCVEWLRRYLENGKETLQRADPPKTHVTHHPVSDHEATLRCWALGFYPAEI

TLTWQRDGEDQTQDTELVETRPAGDRTFQKWAAVVVPSGEEQRYTCHVQHEGLPKPLTLRWEPKSCDKTH
TCPPCPAPELLGGPSVFLFPPKPKDTLMISRTPEVTCVVVDVSHEDPEVKFNWYVDGVEVHNAKTKPREE
QYNSTYRVVSVLTVLHQDWLNGKEYKCKVSNKGLPSSIEKTISKAKGQPREPQVYTLPPSRDELTKNQVS
LTCLVKGFYPSDIAVEWESNGQPENNYKTTPPVLDSDGSFFLYSKLTVDKSRWQQGNVFSCSVMHEALHN
HYTQKSLSLSPGK**ITIFITLFLLSVCYSATVTFFKVKWIFSSVVDLKQTIIPDYRNMIGQGA**

>B*54:01-Final

**MSRSVALAVLALLSLSGLEA**IQRTPKIQVYSRHPAENGKSNFLNCYVSGFHPSDIEVDLLKNGERIEKVE
HSDLSFSKDWSFYLLYYTEFTPTEKDEYACRVNHVTLSQPKIVKWGKSYILL**GGGGSGGGGSGGGGS**GSH
SMRYFYTAMSRPGRGEPRFIAVGYVDDTQFVRFDSDAASPRGEPRAPWVEQEGPEYWDRNTQIYKAQAQT
DRESLRNLRGYYNQSEAGSHTWQTMYGCDLGPDGRLLRGHNQLAYDGKDYIALNEDLSSWTAADTAAQIT
QRKWEAARVAEQLRAYLEGTCVEWLRRYLENGKETLQRADPPKTHVTHHPISDHEATLRCWALGFYPAEI
TLTWQRDGEDQTQDTELVETRPAGDRTFQKWAAVVVPSGEEQRYTCHVQHEGLPKPLTLRWEPKSCDKTH
TCPPCPAPELLGGPSVFLFPPKPKDTLMISRTPEVTCVVVDVSHEDPEVKFNWYVDGVEVHNAKTKPREE
QYNSTYRVVSVLTVLHQDWLNGKEYKCKVSNKGLPSSIEKTISKAKGQPREPQVYTLPPSRDELTKNQVS
LTCLVKGFYPSDIAVEWESNGQPENNYKTTPPVLDSDGSFFLYSKLTVDKSRWQQGNVFSCSVMHEALHN
HYTQKSLSLSPGK**ITIFITLFLLSVCYSATVTFFKVKWIFSSVVDLKQTIIPDYRNMIGQGA**

>B*55:01-Final

**MSRSVALAVLALLSLSGLEA**IQRTPKIQVYSRHPAENGKSNFLNCYVSGFHPSDIEVDLLKNGERIEKVE
HSDLSFSKDWSFYLLYYTEFTPTEKDEYACRVNHVTLSQPKIVKWGKSYILL**GGGGSGGGGSGGGGS**GSH
SMRYFYTAMSRPGRGEPRFIAVGYVDDTQFVRFDSDAASPREEPRAPWIEQEGPEYWDRNTQIYKAQAQT
DRESLRNLRGYYNQSEAGSHTWQTMYGCDLGPDGRLLRGHNQLAYDGKDYIALNEDLSSWTAADTAAQIT
QRKWEAAREAEQLRAYLEGTCVEWLRRYLENGKETLQRADPPKTHVTHHPISDHEATLRCWALGFYPAEI
TLTWQRDGEDQTQDTELVETRPAGDRTFQKWAAVVVPSGEEQRYTCHVQHEGLPKPLTLRWEPKSCDKTH
TCPPCPAPELLGGPSVFLFPPKPKDTLMISRTPEVTCVVVDVSHEDPEVKFNWYVDGVEVHNAKTKPREE
QYNSTYRVVSVLTVLHQDWLNGKEYKCKVSNKGLPSSIEKTISKAKGQPREPQVYTLPPSRDELTKNQVS
LTCLVKGFYPSDIAVEWESNGQPENNYKTTPPVLDSDGSFFLYSKLTVDKSRWQQGNVFSCSVMHEALHN
HYTQKSLSLSPGK**ITIFITLFLLSVCYSATVTFFKVKWIFSSVVDLKQTIIPDYRNMIGQGA**

>B*56:01-Final

**MSRSVALAVLALLSLSGLEA**IQRTPKIQVYSRHPAENGKSNFLNCYVSGFHPSDIEVDLLKNGERIEKVE
HSDLSFSKDWSFYLLYYTEFTPTEKDEYACRVNHVTLSQPKIVKWGKSYILL**GGGGSGGGGSGGGGS**GSH
SMRYFYTAMSRPGRGEPRFIAVGYVDDTQFVRFDSDAASPREEPRAPWIEQEGPEYWDRNTQIYKAQAQT
DRESLRNLRGYYNQSEAGSHTWQTMYGCDLGPDGRLLRGHNQLAYDGKDYIALNEDLSSWTAADTAAQIT
QRKWEAARVAEQLRAYLEGLCVEWLRRYLENGKETLQRADPPKTHVTHHPISDHEATLRCWALGFYPAEI
TLTWQRDGEDQTQDTELVETRPAGDRTFQKWAAVVVPSGEEQRYTCHVQHEGLPKPLTLRWEPKSCDKTH
TCPPCPAPELLGGPSVFLFPPKPKDTLMISRTPEVTCVVVDVSHEDPEVKFNWYVDGVEVHNAKTKPREE
QYNSTYRVVSVLTVLHQDWLNGKEYKCKVSNKGLPSSIEKTISKAKGQPREPQVYTLPPSRDELTKNQVS
LTCLVKGFYPSDIAVEWESNGQPENNYKTTPPVLDSDGSFFLYSKLTVDKSRWQQGNVFSCSVMHEALHN
HYTQKSLSLSPGK**ITIFITLFLLSVCYSATVTFFKVKWIFSSVVDLKQTIIPDYRNMIGQGA**

>B*57:01-Final

**MSRSVALAVLALLSLSGLEA**IQRTPKIQVYSRHPAENGKSNFLNCYVSGFHPSDIEVDLLKNGERIEKVE
HSDLSFSKDWSFYLLYYTEFTPTEKDEYACRVNHVTLSQPKIVKWGKSYILL**GGGGSGGGGSGGGGS**GSH
SMRYFYTAMSRPGRGEPRFIAVGYVDDTQFVRFDSDAASPRMAPRAPWIEQEGPEYWDGETRNMKASAQT
YRENLRIALRYYNQSEAGSHIIQVMYGCDVGPDGRLLRGHDQSAYDGKDYIALNEDLSSWTAADTAAQIT
QRKWEAARVAEQLRAYLEGLCVEWLRRYLENGKETLQRADPPKTHVTHHPISDHEATLRCWALGFYPAEI
TLTWQRDGEDQTQDTELVETRPAGDRTFQKWAAVVVPSGEEQRYTCHVQHEGLPKPLTLRWEPKSCDKTH
TCPPCPAPELLGGPSVFLFPPKPKDTLMISRTPEVTCVVVDVSHEDPEVKFNWYVDGVEVHNAKTKPREE
QYNSTYRVVSVLTVLHQDWLNGKEYKCKVSNKGLPSSIEKTISKAKGQPREPQVYTLPPSRDELTKNQVS
LTCLVKGFYPSDIAVEWESNGQPENNYKTTPPVLDSDGSFFLYSKLTVDKSRWQQGNVFSCSVMHEALHN
HYTQKSLSLSPGK**ITIFITLFLLSVCYSATVTFFKVKWIFSSVVDLKQTIIPDYRNMIGQGA**

>B*57:03-Final

**MSRSVALAVLALLSLSGLEA**IQRTPKIQVYSRHPAENGKSNFLNCYVSGFHPSDIEVDLLKNGERIEKVE
HSDLSFSKDWSFYLLYYTEFTPTEKDEYACRVNHVTLSQPKIVKWGKSYILL**GGGGSGGGGSGGGGS**GSH
SMRYFYTAMSRPGRGEPRFIAVGYVDDTQFVRFDSDAASPRMAPRAPWIEQEGPEYWDGETRNMKASAQT
YRENLRIALRYYNQSEAGSHIIQVMYGCDVGPDGRLLRGHNQYAYDGKDYIALNEDLSSWTAADTAAQIT
QRKWEAARVAEQLRAYLEGLCVEWLRRYLENGKETLQRADPPKTHVTHHPISDHEATLRCWALGFYPAEI
TLTWQRDGEDQTQDTELVETRPAGDRTFQKWAAVVVPSGEEQRYTCHVQHEGLPKPLTLRWEPKSCDKTH
TCPPCPAPELLGGPSVFLFPPKPKDTLMISRTPEVTCVVVDVSHEDPEVKFNWYVDGVEVHNAKTKPREE
QYNSTYRVVSVLTVLHQDWLNGKEYKCKVSNKGLPSSIEKTISKAKGQPREPQVYTLPPSRDELTKNQVS
LTCLVKGFYPSDIAVEWESNGQPENNYKTTPPVLDSDGSFFLYSKLTVDKSRWQQGNVFSCSVMHEALHN
HYTQKSLSLSPGK**ITIFITLFLLSVCYSATVTFFKVKWIFSSVVDLKQTIIPDYRNMIGQGA**

>B*58:01-Final

**MSRSVALAVLALLSLSGLEA**IQRTPKIQVYSRHPAENGKSNFLNCYVSGFHPSDIEVDLLKNGERIEKVE
HSDLSFSKDWSFYLLYYTEFTPTEKDEYACRVNHVTLSQPKIVKWGKSYILL**GGGGSGGGGSGGGGS**GSH
SMRYFYTAMSRPGRGEPRFIAVGYVDDTQFVRFDSDAASPRTEPRAPWIEQEGPEYWDGETRNMKASAQT
YRENLRIALRYYNQSEAGSHIIQRMYGCDLGPDGRLLRGHDQSAYDGKDYIALNEDLSSWTAADTAAQIT
QRKWEAARVAEQLRAYLEGLCVEWLRRYLENGKETLQRADPPKTHVTHHPVSDHEATLRCWALGFYPAEI
TLTWQRDGEDQTQDTELVETRPAGDRTFQKWAAVVVPSGEEQRYTCHVQHEGLPKPLTLRWEPKSCDKTH
TCPPCPAPELLGGPSVFLFPPKPKDTLMISRTPEVTCVVVDVSHEDPEVKFNWYVDGVEVHNAKTKPREE
QYNSTYRVVSVLTVLHQDWLNGKEYKCKVSNKGLPSSIEKTISKAKGQPREPQVYTLPPSRDELTKNQVS
LTCLVKGFYPSDIAVEWESNGQPENNYKTTPPVLDSDGSFFLYSKLTVDKSRWQQGNVFSCSVMHEALHN
HYTQKSLSLSPGK**ITIFITLFLLSVCYSATVTFFKVKWIFSSVVDLKQTIIPDYRNMIGQGA**

>B*59:01-Final

**MSRSVALAVLALLSLSGLEA**IQRTPKIQVYSRHPAENGKSNFLNCYVSGFHPSDIEVDLLKNGERIEKVE
HSDLSFSKDWSFYLLYYTEFTPTEKDEYACRVNHVTLSQPKIVKWGKSYILL**GGGGSGGGGSGGGGS**GSH
SMRYFYTAMSRPGRGEPRFIAVGYVDDTQFVRFDSDAASPREEPRAPWIEQEGPEYWDRNTQIFKTNTQT
YRENLRIALRYYNQSEAGSHTWQTMYGCDLGPDGRLLRGHNQLAYDGKDYIALNEDLSSWTAADTAAQIT
QRKWEAARVAEQLRAYLEGTCVEWLRRYLENGKETLQRADPPKTHVTHHPISDHEATLRCWALGFYPAEI
TLTWQRDGEDQTQDTELVETRPAGDRTFQKWAAVVVPSGEEQRYTCHVQHEGLPKPLTLRWEPKSCDKTH

TCPPCPAPELLGGPSVFLFPPKPKDTLMISRTPEVTCVVVDVSHEDPEVKFNWYVDGVEVHNAKTKPREE
QYNSTYRVVSVLTVLHQDWLNGKEYKCKVSNKGLPSSIEKTISKAKGQPREPQVYTLPPSRDELTKNQVS
LTCLVKGFYPSDIAVEWESNGQPENNYKTTPPVLDSDGSFFLYSKLTVDKSRWQQGNVFSCSVMHEALHN
HYTQKSLSLSPGK**ITIFITLFLLSVCYSATVTFFKVKWIFSSVVDLKQTIIPDYRNMIGQGA**

>B*40:01-Final

**MSRSVALAVLALLSLSGLEA**IQRTPKIQVYSRHPAENGKSNFLNCYVSGFHPSDIEVDLLKNGERIEKVE
HSDLSFSKDWSFYLLYYTEFTPTEKDEYACRVNHVTLSQPKIVKWGKSYILL**GGGGSGGGGSGGGGS**GSH
SMRYFHTAMSRPGRGEPRFITVGYVDDTLFVRFDSDATSPRKEPRAPWIEQEGPEYWDRETQISKTNTQT
YRESLRNLRGYYNQSEAGSHTLQRMYGCDVGPDGRLLRGHNQYAYDGKDYIALNEDLRSWTAADTAAQIS
QRKLEAARVAEQLRAYLEGECVEWLRRYLENGKDKLERADPPKTHVTHHPISDHEATLRCWALGFYPAEI
TLTWQRDGEDQTQDTELVETRPAGDRTFQKWAAVVVPSGEEQRYTCHVQHEGLPKPLTLRWEPKSCDKTH
TCPPCPAPELLGGPSVFLFPPKPKDTLMISRTPEVTCVVVDVSHEDPEVKFNWYVDGVEVHNAKTKPREE
QYNSTYRVVSVLTVLHQDWLNGKEYKCKVSNKGLPSSIEKTISKAKGQPREPQVYTLPPSRDELTKNQVS
LTCLVKGFYPSDIAVEWESNGQPENNYKTTPPVLDSDGSFFLYSKLTVDKSRWQQGNVFSCSVMHEALHN
HYTQKSLSLSPGK**ITIFITLFLLSVCYSATVTFFKVKWIFSSVVDLKQTIIPDYRNMIGQGA**

>B*40:02-Final

**MSRSVALAVLALLSLSGLEA**IQRTPKIQVYSRHPAENGKSNFLNCYVSGFHPSDIEVDLLKNGERIEKVE
HSDLSFSKDWSFYLLYYTEFTPTEKDEYACRVNHVTLSQPKIVKWGKSYILL**GGGGSGGGGSGGGGS**GSH
SMRYFHTSVSRPGRGEPRFITVGYVDDTLFVRFDSDATSPRKEPRAPWIEQEGPEYWDRETQISKTNTQT
YRESLRNLRGYYNQSEAGSHTLQSMYGCDVGPDGRLLRGHNQYAYDGKDYIALNEDLRSWTAADTAAQIT
QRKWEAARVAEQLRAYLEGECVEWLRRYLENGKETLQRADPPKTHVTHHPISDHEATLRCWALGFYPAEI
TLTWQRDGEDQTQDTELVETRPAGDRTFQKWAAVVVPSGEEQRYTCHVQHEGLPKPLTLRWEPKSCDKTH
TCPPCPAPELLGGPSVFLFPPKPKDTLMISRTPEVTCVVVDVSHEDPEVKFNWYVDGVEVHNAKTKPREE
QYNSTYRVVSVLTVLHQDWLNGKEYKCKVSNKGLPSSIEKTISKAKGQPREPQVYTLPPSRDELTKNQVS
LTCLVKGFYPSDIAVEWESNGQPENNYKTTPPVLDSDGSFFLYSKLTVDKSRWQQGNVFSCSVMHEALHN
HYTQKSLSLSPGK**ITIFITLFLLSVCYSATVTFFKVKWIFSSVVDLKQTIIPDYRNMIGQGA**

>B*40:06-Final

**MSRSVALAVLALLSLSGLEA**IQRTPKIQVYSRHPAENGKSNFLNCYVSGFHPSDIEVDLLKNGERIEKVE
HSDLSFSKDWSFYLLYYTEFTPTEKDEYACRVNHVTLSQPKIVKWGKSYILL**GGGGSGGGGSGGGGS**GSH
SMRYFHTSVSRPGRGEPRFITVGYVDDTLFVRFDSDATSPRKEPRAPWIEQEGPEYWDRETQISKTNTQT
YRESLRNLRGYYNQSEAGSHTWQTMYGCDVGPDGRLLRGHNQYAYDGKDYIALNEDLRSWTAADTAAQIT
QRKWEAARVAEQLRAYLEGECVEWLRRYLENGKETLQRADPPKTHVTHHPISDHEATLRCWALGFYPAEI
TLTWQRDGEDQTQDTELVETRPAGDRTFQKWAAVVVPSGEEQRYTCHVQHEGLPKPLTLRWEPKSCDKTH
TCPPCPAPELLGGPSVFLFPPKPKDTLMISRTPEVTCVVVDVSHEDPEVKFNWYVDGVEVHNAKTKPREE
QYNSTYRVVSVLTVLHQDWLNGKEYKCKVSNKGLPSSIEKTISKAKGQPREPQVYTLPPSRDELTKNQVS
LTCLVKGFYPSDIAVEWESNGQPENNYKTTPPVLDSDGSFFLYSKLTVDKSRWQQGNVFSCSVMHEALHN
HYTQKSLSLSPGK**ITIFITLFLLSVCYSATVTFFKVKWIFSSVVDLKQTIIPDYRNMIGQGA**

>B*15:01-Final

**MSRSVALAVLALLSLSGLEA**IQRTPKIQVYSRHPAENGKSNFLNCYVSGFHPSDIEVDLLKNGERIEKVE
HSDLSFSKDWSFYLLYYTEFTPTEKDEYACRVNHVTLSQPKIVKWGKSYILL**GGGGSGGGGSGGGGS**GSH
SMRYFYTAMSRPGRGEPRFIAVGYVDDTQFVRFDSDAASPRMAPRAPWIEQEGPEYWDRETQISKTNTQT
YRESLRNLRGYYNQSEAGSHTLQRMYGCDVGPDGRLLRGHDQSAYDGKDYIALNEDLSSWTAADTAAQIT
QRKWEAAREAEQWRAYLEGLCVEWLRRYLENGKETLQRADPPKTHVTHHPISDHEATLRCWALGFYPAEI
TLTWQRDGEDQTQDTELVETRPAGDRTFQKWAAVVVPSGEEQRYTCHVQHEGLPKPLTLRWEPKSCDKTH
TCPPCPAPELLGGPSVFLFPPKPKDTLMISRTPEVTCVVVDVSHEDPEVKFNWYVDGVEVHNAKTKPREE
QYNSTYRVVSVLTVLHQDWLNGKEYKCKVSNKGLPSSIEKTISKAKGQPREPQVYTLPPSRDELTKNQVS
LTCLVKGFYPSDIAVEWESNGQPENNYKTTPPVLDSDGSFFLYSKLTVDKSRWQQGNVFSCSVMHEALHN
HYTQKSLSLSPGK**ITIFITLFLLSVCYSATVTFFKVKWIFSSVVDLKQTIIPDYRNMIGQGA**

>B*15:16-Final

**MSRSVALAVLALLSLSGLEA**IQRTPKIQVYSRHPAENGKSNFLNCYVSGFHPSDIEVDLLKNGERIEKVE
HSDLSFSKDWSFYLLYYTEFTPTEKDEYACRVNHVTLSQPKIVKWGKSYILL**GGGGSGGGGSGGGGS**GSH
FMRYFYTAMSRPGRGEPRFIAVGYVDDTQFVRFDSDAASPRMAPRAPWIEQEGPEYWDRETRNMKASAQT
YRENLRIALRYYNQSEAGSHTWQRMYGCDLGPDGRLLRGHDQSAYDGKDYIALNEDLSSWTAADTAAQIT
QRKWEAAREAEQLRAYLEGLCVEWLRRYLENGKETLQRADPPKTHVTHHPISDHEATLRCWALGFYPAEI
TLTWQRDGEDQTQDTELVETRPAGDRTFQKWAAVVVPSGEEQRYTCHVQHEGLPKPLTLRWEPKSCDKTH
TCPPCPAPELLGGPSVFLFPPKPKDTLMISRTPEVTCVVVDVSHEDPEVKFNWYVDGVEVHNAKTKPREE
QYNSTYRVVSVLTVLHQDWLNGKEYKCKVSNKGLPSSIEKTISKAKGQPREPQVYTLPPSRDELTKNQVS
LTCLVKGFYPSDIAVEWESNGQPENNYKTTPPVLDSDGSFFLYSKLTVDKSRWQQGNVFSCSVMHEALHN
HYTQKSLSLSPGK**ITIFITLFLLSVCYSATVTFFKVKWIFSSVVDLKQTIIPDYRNMIGQGA**

>B*14:01-Final

**MSRSVALAVLALLSLSGLEA**IQRTPKIQVYSRHPAENGKSNFLNCYVSGFHPSDIEVDLLKNGERIEKVE
HSDLSFSKDWSFYLLYYTEFTPTEKDEYACRVNHVTLSQPKIVKWGKSYILL**GGGGSGGGGSGGGGS**GSH
SMRYFYTSVSRPGRGEPRFISVGYVDDTQFVRFDSDAASPREEPRAPWIEQEGPEYWDRNTQICKTNTQT
DRESLRNLRGYYNQSEAGSHTLQWMYGCDVGPDGRLLRGYNQFAYDGKDYIALNEDLSSWTAADTAAQIT
QRKWEAAREAEQLRAYLEGTCVEWLRRHLENGKETLQRADPPKTHVTHHPISDHEATLRCWALGFYPAEI
TLTWQRDGEDQTQDTELVETRPAGDRTFQKWAAVVVPSGEEQRYTCHVQHEGLPKPLTLRWEPKSCDKTH
TCPPCPAPELLGGPSVFLFPPKPKDTLMISRTPEVTCVVVDVSHEDPEVKFNWYVDGVEVHNAKTKPREE
QYNSTYRVVSVLTVLHQDWLNGKEYKCKVSNKGLPSSIEKTISKAKGQPREPQVYTLPPSRDELTKNQVS
LTCLVKGFYPSDIAVEWESNGQPENNYKTTPPVLDSDGSFFLYSKLTVDKSRWQQGNVFSCSVMHEALHN
HYTQKSLSLSPGK**ITIFITLFLLSVCYSATVTFFKVKWIFSSVVDLKQTIIPDYRNMIGQGA**

>B*14:02-Final

**MSRSVALAVLALLSLSGLEA**IQRTPKIQVYSRHPAENGKSNFLNCYVSGFHPSDIEVDLLKNGERIEKVE
HSDLSFSKDWSFYLLYYTEFTPTEKDEYACRVNHVTLSQPKIVKWGKSYILL**GGGGSGGGGSGGGGS**GSH
SMRYFYTAVSRPGRGEPRFISVGYVDDTQFVRFDSDAASPREEPRAPWIEQEGPEYWDRNTQICKTNTQT
DRESLRNLRGYYNQSEAGSHTLQWMYGCDVGPDGRLLRGYNQFAYDGKDYIALNEDLSSWTAADTAAQIT
QRKWEAAREAEQLRAYLEGTCVEWLRRHLENGKETLQRADPPKTHVTHHPISDHEATLRCWALGFYPAEI
TLTWQRDGEDQTQDTELVETRPAGDRTFQKWAAVVVPSGEEQRYTCHVQHEGLPKPLTLRWEPKSCDKTH

TCPPCPAPELLGGPSVFLFPPKPKDTLMISRTPEVTCVVVDVSHEDPEVKFNWYVDGVEVHNAKTKPREE
QYNSTYRVVSVLTVLHQDWLNGKEYKCKVSNKGLPSSIEKTISKAKGQPREPQVYTLPPSRDELTKNQVS
LTCLVKGFYPSDIAVEWESNGQPENNYKTTPPVLDSDGSFFLYSKLTVDKSRWQQGNVFSCSVMHEALHN
HYTQKSLSLSPGK**ITIFITLFLLSVCYSATVTFFKVKWIFSSVVDLKQTIIPDYRNMIGQGA**

>B*67:01-Final

**MSRSVALAVLALLSLSGLEA**IQRTPKIQVYSRHPAENGKSNFLNCYVSGFHPSDIEVDLLKNGERIEKVE
HSDLSFSKDWSFYLLYYTEFTPTEKDEYACRVNHVTLSQPKIVKWGKSYILL**GGGGSGGGGSGGGGS**GSH
SMRYFYTSVSRPGRGEPRFISVGYVDDTQFVRFDSDAASPREEPRAPWIEQEGPEYWDRNTQIYKAQAQT
DRESLRNLRGYYNQSEAGSHTLQRMYGCDVGPDGRLLRGHNQFAYDGKDYIALNEDLSSWTAADTAAQIT
QRKWEAARVAEQLRTYLEGTCVEWLRRYLENGKETLQRADPPKTHVTHHPISDHEATLRCWALGFYPAEI
TLTWQRDGEDQTQDTELVETRPAGDRTFQKWAAVVVPSGEEQRYTCHVQHEGLPKPLTLRWEPKSCDKTH
TCPPCPAPELLGGPSVFLFPPKPKDTLMISRTPEVTCVVVDVSHEDPEVKFNWYVDGVEVHNAKTKPREE
QYNSTYRVVSVLTVLHQDWLNGKEYKCKVSNKGLPSSIEKTISKAKGQPREPQVYTLPPSRDELTKNQVS
LTCLVKGFYPSDIAVEWESNGQPENNYKTTPPVLDSDGSFFLYSKLTVDKSRWQQGNVFSCSVMHEALHN
HYTQKSLSLSPGK**ITIFITLFLLSVCYSATVTFFKVKWIFSSVVDLKQTIIPDYRNMIGQGA**

>B*15:10-Final

**MSRSVALAVLALLSLSGLEA**IQRTPKIQVYSRHPAENGKSNFLNCYVSGFHPSDIEVDLLKNGERIEKVE
HSDLSFSKDWSFYLLYYTEFTPTEKDEYACRVNHVTLSQPKIVKWGKSYILL**GGGGSGGGGSGGGGS**GSH
SMRYFYTAMSRPGRGEPRFISVGYVDDTQFVRFDSDAASPREEPRAPWIEQEGPEYWDRNTQICKTNTQT
YRESLRNLRGYYNQSEAGSHTLQRMYGCDVGPDGRLLRGHDQYAYDGKDYIALNEDLSSWTAADTAAQIT
QRKWEAAREAEQLRAYLEGLCVEWLRRYLENGKETLQRADPPKTHVTHHPISDHEATLRCWALGFYPAEI
TLTWQRDGEDQTQDTELVETRPAGDRTFQKWAAVVVPSGEEQRYTCHVQHEGLPKPLTLRWEPKSCDKTH
TCPPCPAPELLGGPSVFLFPPKPKDTLMISRTPEVTCVVVDVSHEDPEVKFNWYVDGVEVHNAKTKPREE
QYNSTYRVVSVLTVLHQDWLNGKEYKCKVSNKGLPSSIEKTISKAKGQPREPQVYTLPPSRDELTKNQVS
LTCLVKGFYPSDIAVEWESNGQPENNYKTTPPVLDSDGSFFLYSKLTVDKSRWQQGNVFSCSVMHEALHN
HYTQKSLSLSPGK**ITIFITLFLLSVCYSATVTFFKVKWIFSSVVDLKQTIIPDYRNMIGQGA**

>B*15:03-Final

**MSRSVALAVLALLSLSGLEA**IQRTPKIQVYSRHPAENGKSNFLNCYVSGFHPSDIEVDLLKNGERIEKVE
HSDLSFSKDWSFYLLYYTEFTPTEKDEYACRVNHVTLSQPKIVKWGKSYILL**GGGGSGGGGSGGGGS**GSH
SMRYFYTAMSRPGRGEPRFISVGYVDDTQFVRFDSDAASPREEPRAPWIEQEGPEYWDRETQISKTNTQT
YRESLRNLRGYYNQSEAGSHTLQRMYGCDVGPDGRLLRGHDQSAYDGKDYIALNEDLSSWTAADTAAQIT
QRKWEAAREAEQLRAYLEGLCVEWLRRYLENGKETLQRADPPKTHVTHHPISDHEATLRCWALGFYPAEI
TLTWQRDGEDQTQDTELVETRPAGDRTFQKWAAVVVPSGEEQRYTCHVQHEGLPKPLTLRWEPKSCDKTH
TCPPCPAPELLGGPSVFLFPPKPKDTLMISRTPEVTCVVVDVSHEDPEVKFNWYVDGVEVHNAKTKPREE
QYNSTYRVVSVLTVLHQDWLNGKEYKCKVSNKGLPSSIEKTISKAKGQPREPQVYTLPPSRDELTKNQVS
LTCLVKGFYPSDIAVEWESNGQPENNYKTTPPVLDSDGSFFLYSKLTVDKSRWQQGNVFSCSVMHEALHN
HYTQKSLSLSPGK**ITIFITLFLLSVCYSATVTFFKVKWIFSSVVDLKQTIIPDYRNMIGQGA**

>B*73:01-Final

**MSRSVALAVLALLSLSGLEA**IQRTPKIQVYSRHPAENGKSNFLNCYVSGFHPSDIEVDLLKNGERIEKVE
HSDLSFSKDWSFYLLYYTEFTPTEKDEYACRVNHVTLSQPKIVKWGKSYILL**GGGGSGGGGSGGGGS**GSH
SMRYFHTSVSRPGRGEPRFITVGYVDDTQFVRFDSDAASPREEPRAPWIEQEGPEYWDRNTQICKAKAQT
DRVGLRNLRGYYNQSEDGSHTWQTMYGCDMGPDGRLLRGYNQFAYDGKDYIALNEDLRSWTAADTAAQIT
QRKWEAARVAEQLRAYLEGECVEWLRRHLENGKETLQRADPPKTHVTHHPISDHEATLRCWALGFYPAEI
TLTWQRDGEDQTQDTELVETRPAGDGTFQKWAAVVVPSGQEQRYTCHVQHEGLQEPCTLRWEPKSCDKTH
TCPPCPAPELLGGPSVFLFPPKPKDTLMISRTPEVTCVVVDVSHEDPEVKFNWYVDGVEVHNAKTKPREE
QYNSTYRVVSVLTVLHQDWLNGKEYKCKVSNKGLPSSIEKTISKAKGQPREPQVYTLPPSRDELTKNQVS
LTCLVKGFYPSDIAVEWESNGQPENNYKTTPPVLDSDGSFFLYSKLTVDKSRWQQGNVFSCSVMHEALHN
HYTQKSLSLSPGK**ITIFITLFLLSVCYSATVTFFKVKWIFSSVVDLKQTIIPDYRNMIGQGA**

>B*15:02-Final

**MSRSVALAVLALLSLSGLEA**IQRTPKIQVYSRHPAENGKSNFLNCYVSGFHPSDIEVDLLKNGERIEKVE
HSDLSFSKDWSFYLLYYTEFTPTEKDEYACRVNHVTLSQPKIVKWGKSYILL**GGGGSGGGGSGGGGS**GSH
SMRYFYTAMSRPGRGEPRFIAVGYVDDTQFVRFDSDAASPRMAPRAPWIEQEGPEYWDRNTQISKTNTQT
YRESLRNLRGYYNQSEAGSHIIQRMYGCDVGPDGRLLRGYDQSAYDGKDYIALNEDLSSWTAADTAAQIT
QRKWEAAREAEQLRAYLEGLCVEWLRRYLENGKETLQRADPPKTHVTHHPISDHEATLRCWALGFYPAEI
TLTWQRDGEDQTQDTELVETRPAGDRTFQKWAAVVVPSGEEQRYTCHVQHEGLPKPLTLRWEPKSCDKTH
TCPPCPAPELLGGPSVFLFPPKPKDTLMISRTPEVTCVVVDVSHEDPEVKFNWYVDGVEVHNAKTKPREE
QYNSTYRVVSVLTVLHQDWLNGKEYKCKVSNKGLPSSIEKTISKAKGQPREPQVYTLPPSRDELTKNQVS
LTCLVKGFYPSDIAVEWESNGQPENNYKTTPPVLDSDGSFFLYSKLTVDKSRWQQGNVFSCSVMHEALHN
HYTQKSLSLSPGK**ITIFITLFLLSVCYSATVTFFKVKWIFSSVVDLKQTIIPDYRNMIGQGA**

>B*15:11-Final

**MSRSVALAVLALLSLSGLEA**IQRTPKIQVYSRHPAENGKSNFLNCYVSGFHPSDIEVDLLKNGERIEKVE
HSDLSFSKDWSFYLLYYTEFTPTEKDEYACRVNHVTLSQPKIVKWGKSYILL**GGGGSGGGGSGGGGS**GSH
SMRYFYTAMSRPGRGEPRFIAVGYVDDTQFVRFDSDAASPRMAPRAPWIEQEGPEYWDRNTQIYKTNTQT
YRESLRNLRGYYNQSEAGSHTLQRMYGCDVGPDGRLLRGHDQSAYDGKDYIALNEDLSSWTAADTAAQIT
QRKWEAAREAEQWRAYLEGLCVEWLRRYLENGKETLQRADPPKTHVTHHPISDHEATLRCWALGFYPAEI
TLTWQRDGEDQTQDTELVETRPAGDRTFQKWAAVVVPSGEEQRYTCHVQHEGLPKPLTLRWEPKSCDKTH
TCPPCPAPELLGGPSVFLFPPKPKDTLMISRTPEVTCVVVDVSHEDPEVKFNWYVDGVEVHNAKTKPREE
QYNSTYRVVSVLTVLHQDWLNGKEYKCKVSNKGLPSSIEKTISKAKGQPREPQVYTLPPSRDELTKNQVS
LTCLVKGFYPSDIAVEWESNGQPENNYKTTPPVLDSDGSFFLYSKLTVDKSRWQQGNVFSCSVMHEALHN
HYTQKSLSLSPGK**ITIFITLFLLSVCYSATVTFFKVKWIFSSVVDLKQTIIPDYRNMIGQGA**

>B*15:12-Final

**MSRSVALAVLALLSLSGLEA**IQRTPKIQVYSRHPAENGKSNFLNCYVSGFHPSDIEVDLLKNGERIEKVE
HSDLSFSKDWSFYLLYYTEFTPTEKDEYACRVNHVTLSQPKIVKWGKSYILL**GGGGSGGGGSGGGGS**GSH
SMRYFYTAMSRPGRGEPRFIAVGYVDDTQFVRFDSDAASPRMAPRAPWIEQEGPEYWDRETQISKTNTQT
YRESLRNLRGYYNQSEAGSHTLQRMYGCDVGPDGRLLRGHDQSAYDGKDYIALNEDLSSWTAADTAAQIT
QRKWEAAREAEQWRAYLEGLCVDGLRRYLENGKETLQRADPPKTHVTHHPISDHEATLRCWALGFYPAEI
TLTWQRDGEDQTQDTELVETRPAGDRTFQKWAAVVVPSGEEQRYTCHVQHEGLPKPLTLRWEPKSCDKTH

TCPPCPAPELLGGPSVFLFPPKPKDTLMISRTPEVTCVVVDVSHEDPEVKFNWYVDGVEVHNAKTKPREE
QYNSTYRVVSVLTVLHQDWLNGKEYKCKVSNKGLPSSIEKTISKAKGQPREPQVYTLPPSRDELTKNQVS
LTCLVKGFYPSDIAVEWESNGQPENNYKTTPPVLDSDGSFFLYSKLTVDKSRWQQGNVFSCSVMHEALHN
HYTQKSLSLSPGK**ITIFITLFLLSVCYSATVTFFKVKWIFSSVVDLKQTIIPDYRNMIGQGA**

>B*15:13-Final

**MSRSVALAVLALLSLSGLEA**IQRTPKIQVYSRHPAENGKSNFLNCYVSGFHPSDIEVDLLKNGERIEKVE
HSDLSFSKDWSFYLLYYTEFTPTEKDEYACRVNHVTLSQPKIVKWGKSYILL**GGGGSGGGGSGGGGS**GSH
SMRYFYTAMSRPGRGEPRFIAVGYVDDTQFVRFDSDAASPRMAPRAPWIEQEGPEYWDRNTQISKTNTQT
YRENLRIALRYYNQSEAGSHIIQRMYGCDVGPDGRLLRGYDQSAYDGKDYIALNEDLSSWTAADTAAQIT
QRKWEAAREAEQLRAYLEGLCVEWLRRYLENGKETLQRADPPKTHVTHHPISDHEATLRCWALGFYPAEI
TLTWQRDGEDQTQDTELVETRPAGDRTFQKWAAVVVPSGEEQRYTCHVQHEGLPKPLTLRWEPKSCDKTH
TCPPCPAPELLGGPSVFLFPPKPKDTLMISRTPEVTCVVVDVSHEDPEVKFNWYVDGVEVHNAKTKPREE
QYNSTYRVVSVLTVLHQDWLNGKEYKCKVSNKGLPSSIEKTISKAKGQPREPQVYTLPPSRDELTKNQVS
LTCLVKGFYPSDIAVEWESNGQPENNYKTTPPVLDSDGSFFLYSKLTVDKSRWQQGNVFSCSVMHEALHN
HYTQKSLSLSPGK**ITIFITLFLLSVCYSATVTFFKVKWIFSSVVDLKQTIIPDYRNMIGQGA**

>B*81:01-Final

**MSRSVALAVLALLSLSGLEA**IQRTPKIQVYSRHPAENGKSNFLNCYVSGFHPSDIEVDLLKNGERIEKVE
HSDLSFSKDWSFYLLYYTEFTPTEKDEYACRVNHVTLSQPKIVKWGKSYILL**GGGGSGGGGSGGGGS**GSH
SMRYFYTSVSRPGRGEPRFISVGYVDDTQFVRFDSDAASPREEPRAPWIEQEGPEYWDRNTQIYKAQAQT
DRESLRNLRGYYNQSEAGSHTLQSMYGCDVGPDGRLLRGHNQYAYDGKDYIALNEDLRSWTAADTAAQIS
QRKLEAARVAEQLRAYLEGECVEWLRRYLENGKDKLERADPPKTHVTHHPISDHEATLRCWALGFYPAEI
TLTWQRDGEDQTQDTELVETRPAGDRTFQKWTAVVVPSGEEQRYTCHVQHEGLPKPLTLRWEPKSCDKTH
TCPPCPAPELLGGPSVFLFPPKPKDTLMISRTPEVTCVVVDVSHEDPEVKFNWYVDGVEVHNAKTKPREE
QYNSTYRVVSVLTVLHQDWLNGKEYKCKVSNKGLPSSIEKTISKAKGQPREPQVYTLPPSRDELTKNQVS
LTCLVKGFYPSDIAVEWESNGQPENNYKTTPPVLDSDGSFFLYSKLTVDKSRWQQGNVFSCSVMHEALHN
HYTQKSLSLSPGK**ITIFITLFLLSVCYSATVTFFKVKWIFSSVVDLKQTIIPDYRNMIGQGA**

>B*82:01-Final

**MSRSVALAVLALLSLSGLEA**IQRTPKIQVYSRHPAENGKSNFLNCYVSGFHPSDIEVDLLKNGERIEKVE
HSDLSFSKDWSFYLLYYTEFTPTEKDEYACRVNHVTLSQPKIVKWGKSYILL**GGGGSGGGGSGGGGS**GSH
SMRYFYTAMSRPGRGEPRFISVGYVDDTQFVRFDSDAASPREEPRAPWIEQEGPEYWDRNTQIYKAQAQT
DRESLRNLRGYYNQSEAGSHTLQRMFGCDLGPDGRLLRGHNQLAYDGKDYIALNEDLSSWTAADTAAQIT
QRKWEAARVAEQDRAYLEDLCVESLRRYLENGKETLQRADPPKTHVTHHPISDHEATLRCWALGFYPAEI
TLTWQRDGEDQTQDTELVETRPAGDRTFQKWAAVVVPSGEEQRYTCHVQHEGLPKPLTLRWEPKSCDKTH
TCPPCPAPELLGGPSVFLFPPKPKDTLMISRTPEVTCVVVDVSHEDPEVKFNWYVDGVEVHNAKTKPREE
QYNSTYRVVSVLTVLHQDWLNGKEYKCKVSNKGLPSSIEKTISKAKGQPREPQVYTLPPSRDELTKNQVS
LTCLVKGFYPSDIAVEWESNGQPENNYKTTPPVLDSDGSFFLYSKLTVDKSRWQQGNVFSCSVMHEALHN
HYTQKSLSLSPGK**ITIFITLFLLSVCYSATVTFFKVKWIFSSVVDLKQTIIPDYRNMIGQGA**

HLA C

>C*01:02-Final

**MSRSVALAVLALLSLSGLEA**IQRTPKIQVYSRHPAENGKSNFLNCYVSGFHPSDIEVDLLKNGERIEKVE
HSDLSFSKDWSFYLLYYTEFTPTEKDEYACRVNHVTLSQPKIVKWGKSYILL**GGGGSGGGGSGGGGS**CSH
SMKYFFTSVSRPGRGEPRFISVGYVDDTQFVRFDSDAASPRGEPRAPWVEQEGPEYWDRETQKYKRQAQT
DRVSLRNLRGYYNQSEAGSHTLQWMCGCDLGPDGRLLRGYDQYAYDGKDYIALNEDLRSWTAADTAAQIT
QRKWEAAREAEQRRAYLEGTCVEWLRRYLENGKETLQRAEHPKTHVTHHPVSDHEATLRCWALGFYPAEI
TLTWQWDGEDQTQDTELVETRPAGDGTFQKWAAVMVPSGEEQRYTCHVQHEGLPEPLTLRWEPKSCDKTH
TCPPCPAPELLGGPSVFLFPPKPKDTLMISRTPEVTCVVVDVSHEDPEVKFNWYVDGVEVHNAKTKPREE
QYNSTYRVVSVLTVLHQDWLNGKEYKCKVSNKGLPSSIEKTISKAKGQPREPQVYTLPPSRDELTKNQVS
LTCLVKGFYPSDIAVEWESNGQPENNYKTTPPVLDSDGSFFLYSKLTVDKSRWQQGNVFSCSVMHEALHN
HYTQKSLSLSPGK**ITIFITLFLLSVCYSATVTFFKVKWIFSSVVDLKQTIIPDYRNMIGQGA**

>C*02:02-Final

**MSRSVALAVLALLSLSGLEA**IQRTPKIQVYSRHPAENGKSNFLNCYVSGFHPSDIEVDLLKNGERIEKVE
HSDLSFSKDWSFYLLYYTEFTPTEKDEYACRVNHVTLSQPKIVKWGKSYILL**GGGGSGGGGSGGGGS**CSH
SMRYFYTAVSRPSRGEPHFIAVGYVDDTQFVRFDSDAASPRGEPRAPWVEQEGPEYWDRETQKYKRQAQT
DRVNLRKLRGYYNQSEAGSHTLQRMYGCDLGPDGRLLRGYDQSAYDGKDYIALNEDLRSWTAADTAAQIT
QRKWEAAREAEQWRAYLEGECVEWLRRYLENGKETLQRAEHPKTHVTHHPVSDHEATLRCWALGFYPTEI
TLTWQRDGEDQTQDTELVETRPAGDGTFQKWAAVVVPSGEEQRYTCHVQHEGLPEPLTLRWEPKSCDKTH
TCPPCPAPELLGGPSVFLFPPKPKDTLMISRTPEVTCVVVDVSHEDPEVKFNWYVDGVEVHNAKTKPREE
QYNSTYRVVSVLTVLHQDWLNGKEYKCKVSNKGLPSSIEKTISKAKGQPREPQVYTLPPSRDELTKNQVS
LTCLVKGFYPSDIAVEWESNGQPENNYKTTPPVLDSDGSFFLYSKLTVDKSRWQQGNVFSCSVMHEALHN
HYTQKSLSLSPGK**ITIFITLFLLSVCYSATVTFFKVKWIFSSVVDLKQTIIPDYRNMIGQGA**

>C*04:01-Final

**MSRSVALAVLALLSLSGLEA**IQRTPKIQVYSRHPAENGKSNFLNCYVSGFHPSDIEVDLLKNGERIEKVE
HSDLSFSKDWSFYLLYYTEFTPTEKDEYACRVNHVTLSQPKIVKWGKSYILL**GGGGSGGGGSGGGGS**GSH
SMRYFSTSVSWPGRGEPRFIAVGYVDDTQFVRFDSDAASPRGEPREPWVEQEGPEYWDRETQKYKRQAQA
DRVNLRKLRGYYNQSEDGSHTLQRMFGCDLGPDGRLLRGYNQFAYDGKDYIALNEDLRSWTAADTAAQIT
QRKWEAAREAEQRRAYLEGTCVEWLRRYLENGKETLQRAEHPKTHVTHHPVSDHEATLRCWALGFYPAEI
TLTWQWDGEDQTQDTELVETRPAGDGTFQKWAAVVVPSGEEQRYTCHVQHEGLPEPLTLRWEPKSCDKTH
TCPPCPAPELLGGPSVFLFPPKPKDTLMISRTPEVTCVVVDVSHEDPEVKFNWYVDGVEVHNAKTKPREE
QYNSTYRVVSVLTVLHQDWLNGKEYKCKVSNKGLPSSIEKTISKAKGQPREPQVYTLPPSRDELTKNQVS
LTCLVKGFYPSDIAVEWESNGQPENNYKTTPPVLDSDGSFFLYSKLTVDKSRWQQGNVFSCSVMHEALHN
HYTQKSLSLSPGK**ITIFITLFLLSVCYSATVTFFKVKWIFSSVVDLKQTIIPDYRNMIGQGA**

>C*05:01-Final

**MSRSVALAVLALLSLSGLEA**IQRTPKIQVYSRHPAENGKSNFLNCYVSGFHPSDIEVDLLKNGERIEKVE
HSDLSFSKDWSFYLLYYTEFTPTEKDEYACRVNHVTLSQPKIVKWGKSYILL**GGGGSGGGGSGGGGS**CSH
SMRYFYTAVSRPGRGEPRFIAVGYVDDTQFVQFDSDAASPRGEPRAPWVEQEGPEYWDRETQKYKRQAQT

DRVNLRKLRGYYNQSEAGSHTLQRMYGCDLGPDGRLLRGYNQFAYDGKDYIALNEDLRSWTAADKAAQIT
QRKWEAAREAEQRRAYLEGTCVEWLRRYLENGKKTLQRAEHPKTHVTHHPVSDHEATLRCWALGFYPAEI
TLTWQRDGEDQTQDTELVETRPAGDGTFQKWAAVVVPSGEEQRYTCHVQHEGLPEPLTLRWEPKSCDKTH
TCPPCPAPELLGGPSVFLFPPKPKDTLMISRTPEVTCVVVDVSHEDPEVKFNWYVDGVEVHNAKTKPREE
QYNSTYRVVSVLTVLHQDWLNGKEYKCKVSNKGLPSSIEKTISKAKGQPREPQVYTLPPSRDELTKNQVS
LTCLVKGFYPSDIAVEWESNGQPENNYKTTPPVLDSDGSFFLYSKLTVDKSRWQQGNVFSCSVMHEALHN
HYTQKSLSLSPGK**ITIFITLFLLSVCYSATVTFFKVKWIFSSVVDLKQTIIPDYRNMIGQGA**

>C*06:02-Final

**MSRSVALAVLALLSLSGLEA**IQRTPKIQVYSRHPAENGKSNFLNCYVSGFHPSDIEVDLLKNGERIEKVE
HSDLSFSKDWSFYLLYYTEFTPTEKDEYACRVNHVTLSQPKIVKWGKSYILL**GGGGSGGGGSGGGGS**CSH
SMRYFDTAVSRPGRGEPRFISVGYVDDTQFVRFDSDAASPRGEPRAPWVEQEGPEYWDRETQKYKRQAQA
DRVNLRKLRGYYNQSEDGSHTLQWMYGCDLGPDGRLLRGYDQSAYDGKDYIALNEDLRSWTAADTAAQIT
QRKWEAAREAEQWRAYLEGTCVEWLRRYLENGKETLQRAEHPKTHVTHHPVSDHEATLRCWALGFYPAEI
TLTWQRDGEDQTQDTELVETRPAGDGTFQKWAAVVVPSGEEQRYTCHVQHEGLPEPLTLRWEPKSCDKTH
TCPPCPAPELLGGPSVFLFPPKPKDTLMISRTPEVTCVVVDVSHEDPEVKFNWYVDGVEVHNAKTKPREE
QYNSTYRVVSVLTVLHQDWLNGKEYKCKVSNKGLPSSIEKTISKAKGQPREPQVYTLPPSRDELTKNQVS
LTCLVKGFYPSDIAVEWESNGQPENNYKTTPPVLDSDGSFFLYSKLTVDKSRWQQGNVFSCSVMHEALHN
HYTQKSLSLSPGK**ITIFITLFLLSVCYSATVTFFKVKWIFSSVVDLKQTIIPDYRNMIGQGA**

>C*07:02-Final

**MSRSVALAVLALLSLSGLEA**IQRTPKIQVYSRHPAENGKSNFLNCYVSGFHPSDIEVDLLKNGERIEKVE
HSDLSFSKDWSFYLLYYTEFTPTEKDEYACRVNHVTLSQPKIVKWGKSYILL**GGGGSGGGGSGGGGS**CSH
SMRYFDTAVSRPGRGEPRFISVGYVDDTQFVRFDSDAASPRGEPRAPWVEQEGPEYWDRETQKYKRQAQA
DRVSLRNLRGYYNQSEDGSHTLQRMSGCDLGPDGRLLRGYDQSAYDGKDYIALNEDLRSWTAADTAAQIT
QRKLEAARAAEQLRAYLEGTCVEWLRRYLENGKETLQRAEPPKTHVTHHPLSDHEATLRCWALGFYPAEI
TLTWQRDGEDQTQDTELVETRPAGDGTFQKWAAVVVPSGQEQRYTCHMQHEGLQEPLTLSWEPKSCDKTH
TCPPCPAPELLGGPSVFLFPPKPKDTLMISRTPEVTCVVVDVSHEDPEVKFNWYVDGVEVHNAKTKPREE
QYNSTYRVVSVLTVLHQDWLNGKEYKCKVSNKGLPSSIEKTISKAKGQPREPQVYTLPPSRDELTKNQVS
LTCLVKGFYPSDIAVEWESNGQPENNYKTTPPVLDSDGSFFLYSKLTVDKSRWQQGNVFSCSVMHEALHN
HYTQKSLSLSPGK**ITIFITLFLLSVCYSATVTFFKVKWIFSSVVDLKQTIIPDYRNMIGQGA**

>C*08:01-Final

**MSRSVALAVLALLSLSGLEA**IQRTPKIQVYSRHPAENGKSNFLNCYVSGFHPSDIEVDLLKNGERIEKVE
HSDLSFSKDWSFYLLYYTEFTPTEKDEYACRVNHVTLSQPKIVKWGKSYILL**GGGGSGGGGSGGGGS**CSH
SMRYFYTAVSRPGRGEPRFIAVGYVDDTQFVQFDSDAASPRGEPRAPWVEQEGPEYWDRETQKYKRQAQT
DRVSLRNLRGYYNQSEAGSHTLQRMYGCDLGPDGRLLRGYNQFAYDGKDYIALNEDLRSWTAADTAAQIT
QRKWEAARTAEQLRAYLEGTCVEWLRRYLENGKKTLQRAEHPKTHVTHHPVSDHEATLRCWALGFYPAEI
TLTWQRDGEDQTQDTELVETRPAGDGTFQKWAAVVVPSGEEQRYTCHVQHEGLPEPLTLRWEPKSCDKTH
TCPPCPAPELLGGPSVFLFPPKPKDTLMISRTPEVTCVVVDVSHEDPEVKFNWYVDGVEVHNAKTKPREE
QYNSTYRVVSVLTVLHQDWLNGKEYKCKVSNKGLPSSIEKTISKAKGQPREPQVYTLPPSRDELTKNQVS

LTCLVKGFYPSDIAVEWESNGQPENNYKTTPPVLDSDGSFFLYSKLTVDKSRWQQGNVFSCSVMHEALHN
HYTQKSLSLSPGK**ITIFITLFLLSVCYSATVTFFKVKWIFSSVVDLKQTIIPDYRNMIGQGA**

>C*03:03-Final

**MSRSVALAVLALLSLSGLEA**IQRTPKIQVYSRHPAENGKSNFLNCYVSGFHPSDIEVDLLKNGERIEKVE
HSDLSFSKDWSFYLLYYTEFTPTEKDEYACRVNHVTLSQPKIVKWGKSYILL**GGGGSGGGGSGGGGS**GSH
SMRYFYTAVSRPGRGEPHFIAVGYVDDTQFVRFDSDAASPRGEPRAPWVEQEGPEYWDRETQKYKRQAQT
DRVSLRNLRGYYNQSEARSHIIQRMYGCDVGPDGRLLRGYDQYAYDGKDYIALNEDLRSWTAADTAAQIT
QRKWEAAREAEQLRAYLEGLCVEWLRRYLKNGKETLQRAEHPKTHVTHHPVSDHEATLRCWALGFYPAEI
TLTWQWDGEDQTQDTELVETRPAGDGTFQKWAAVVVPSGEEQRYTCHVQHEGLPEPLTLRWEPKSCDKTH
TCPPCPAPELLGGPSVFLFPPKPKDTLMISRTPEVTCVVVDVSHEDPEVKFNWYVDGVEVHNAKTKPREE
QYNSTYRVVSVLTVLHQDWLNGKEYKCKVSNKGLPSSIEKTISKAKGQPREPQVYTLPPSRDELTKNQVS
LTCLVKGFYPSDIAVEWESNGQPENNYKTTPPVLDSDGSFFLYSKLTVDKSRWQQGNVFSCSVMHEALHN
HYTQKSLSLSPGK**ITIFITLFLLSVCYSATVTFFKVKWIFSSVVDLKQTIIPDYRNMIGQGA**

>C*03:02-Final

**MSRSVALAVLALLSLSGLEA**IQRTPKIQVYSRHPAENGKSNFLNCYVSGFHPSDIEVDLLKNGERIEKVE
HSDLSFSKDWSFYLLYYTEFTPTEKDEYACRVNHVTLSQPKIVKWGKSYILL**GGGGSGGGGSGGGGS**GSH
SMRYFYTAVSRPGRGEPHFIAVGYVDDTQFVRFDSDAASPRGEPRAPWVEQEGPEYWDRETQKYKRQAQT
DRVSLRNLRGYYNQSEAGSHILQRMYGCDVGPDGRLLRGYDQSAYDGKDYIALNEDLRSWTAADTAAQIT
QRKWEAAREAEQLRAYLEGLCVEWLRRYLKNGKETLQRAEHPKTHVTHHPVSDHEATLRCWALGFYPAEI
TLTWQWDGEDQTQDTELVETRPAGDGTFQKWAAVVVPSGEEQRYTCHVQHEGLPEPLTLRWEPKSCDKTH
TCPPCPAPELLGGPSVFLFPPKPKDTLMISRTPEVTCVVVDVSHEDPEVKFNWYVDGVEVHNAKTKPREE
QYNSTYRVVSVLTVLHQDWLNGKEYKCKVSNKGLPSSIEKTISKAKGQPREPQVYTLPPSRDELTKNQVS
LTCLVKGFYPSDIAVEWESNGQPENNYKTTPPVLDSDGSFFLYSKLTVDKSRWQQGNVFSCSVMHEALHN
HYTQKSLSLSPGK**ITIFITLFLLSVCYSATVTFFKVKWIFSSVVDLKQTIIPDYRNMIGQGA**

>C*03:04-Final

**MSRSVALAVLALLSLSGLEA**IQRTPKIQVYSRHPAENGKSNFLNCYVSGFHPSDIEVDLLKNGERIEKVE
HSDLSFSKDWSFYLLYYTEFTPTEKDEYACRVNHVTLSQPKIVKWGKSYILL**GGGGSGGGGSGGGGS**GSH
SMRYFYTAVSRPGRGEPHFIAVGYVDDTQFVRFDSDAASPRGEPRAPWVEQEGPEYWDRETQKYKRQAQT
DRVSLRNLRGYYNQSEAGSHIIQRMYGCDVGPDGRLLRGYDQYAYDGKDYIALNEDLRSWTAADTAAQIT
QRKWEAAREAEQLRAYLEGLCVEWLRRYLKNGKETLQRAEHPKTHVTHHPVSDHEATLRCWALGFYPAEI
TLTWQWDGEDQTQDTELVETRPAGDGTFQKWAAVVVPSGEEQRYTCHVQHEGLPEPLTLRWEPKSCDKTH
TCPPCPAPELLGGPSVFLFPPKPKDTLMISRTPEVTCVVVDVSHEDPEVKFNWYVDGVEVHNAKTKPREE
QYNSTYRVVSVLTVLHQDWLNGKEYKCKVSNKGLPSSIEKTISKAKGQPREPQVYTLPPSRDELTKNQVS
LTCLVKGFYPSDIAVEWESNGQPENNYKTTPPVLDSDGSFFLYSKLTVDKSRWQQGNVFSCSVMHEALHN
HYTQKSLSLSPGK**ITIFITLFLLSVCYSATVTFFKVKWIFSSVVDLKQTIIPDYRNMIGQGA**

>C*12:03-Final

**MSRSVALAVLALLSLSGLEA**IQRTPKIQVYSRHPAENGKSNFLNCYVSGFHPSDIEVDLLKNGERIEKVE
HSDLSFSKDWSFYLLYYTEFTPTEKDEYACRVNHVTLSQPKIVKWGKSYILL**GGGGSGGGGSGGGGS**CSH
SMRYFYTAVSRPGRGEPRFIAVGYVDDTQFVRFDSDAASPRGEPRAPWVEQEGPEYWDRETQKYKRQAQA
DRVSLRNLRGYYNQSEAGSHTLQWMYGCDLGPDGRLLRGYDQSAYDGKDYIALNEDLRSWTAADTAAQIT
QRKWEAAREAEQWRAYLEGTCVEWLRRYLENGKETLQRAEHPKTHVTHHPVSDHEATLRCWALGFYPAEI
TLTWQRDGEDQTQDTELVETRPAGDGTFQKWAAVVVPSGEEQRYTCHVQHEGLPEPLTLRWEPKSCDKTH
TCPPCPAPELLGGPSVFLFPPKPKDTLMISRTPEVTCVVVDVSHEDPEVKFNWYVDGVEVHNAKTKPREE
QYNSTYRVVSVLTVLHQDWLNGKEYKCKVSNKGLPSSIEKTISKAKGQPREPQVYTLPPSRDELTKNQVS
LTCLVKGFYPSDIAVEWESNGQPENNYKTTPPVLDSDGSFFLYSKLTVDKSRWQQGNVFSCSVMHEALHN
HYTQKSLSLSPGK**ITIFITLFLLSVCYSATVTFFKVKWIFSSVVDLKQTIIPDYRNMIGQGA**

>C*14:02-Final

**MSRSVALAVLALLSLSGLEA**IQRTPKIQVYSRHPAENGKSNFLNCYVSGFHPSDIEVDLLKNGERIEKVE
HSDLSFSKDWSFYLLYYTEFTPTEKDEYACRVNHVTLSQPKIVKWGKSYILL**GGGGSGGGGSGGGGS**CSH
SMRYFSTSVSRPGRGEPRFIAVGYVDDTQFVRFDSDAASPRGEPRAPWVEQEGPEYWDRETQKYKRQAQT
DRVSLRNLRGYYNQSEAGSHTLQWMFGCDLGPDGRLLRGYDQSAYDGKDYIALNEDLRSWTAADTAAQIT
QRKWEAAREAEQRRAYLEGTCVEWLRRYLENGKETLQRAEHPKTHVTHHPVSDHEATLRCWALGFYPAEI
TLTWQWDGEDQTQDTELVETRPAGDGTFQKWAAVVVPSGEEQRYTCHVQHEGLPEPLTLRWEPKSCDKTH
TCPPCPAPELLGGPSVFLFPPKPKDTLMISRTPEVTCVVVDVSHEDPEVKFNWYVDGVEVHNAKTKPREE
QYNSTYRVVSVLTVLHQDWLNGKEYKCKVSNKGLPSSIEKTISKAKGQPREPQVYTLPPSRDELTKNQVS
LTCLVKGFYPSDIAVEWESNGQPENNYKTTPPVLDSDGSFFLYSKLTVDKSRWQQGNVFSCSVMHEALHN
HYTQKSLSLSPGK**ITIFITLFLLSVCYSATVTFFKVKWIFSSVVDLKQTIIPDYRNMIGQGA**

>C*15:02-Final

**MSRSVALAVLALLSLSGLEA**IQRTPKIQVYSRHPAENGKSNFLNCYVSGFHPSDIEVDLLKNGERIEKVE
HSDLSFSKDWSFYLLYYTEFTPTEKDEYACRVNHVTLSQPKIVKWGKSYILL**GGGGSGGGGSGGGGS**CSH
SMRYFYTAVSRPGRGEPHFIAVGYVDDTQFVRFDSDAASPRGEPRAPWVEQEGPEYWDRETQNYKRQAQT
DRVNLRKLRGYYNQSEAGSHIIQRMYGCDLGPDGRLLRGHDQLAYDGKDYIALNEDLRSWTAADTAAQIT
QRKWEAAREAEQLRAYLEGTCVEWLRRYLENGKETLQRAEHPKTHVTHHPVSDHEATLRCWALGFYPAEI
TLTWQRDGEDQTQDTELVETRPAGDGTFQKWAAVVVPSGEEQRYTCHVQHEGLPEPLTLRWEPKSCDKTH
TCPPCPAPELLGGPSVFLFPPKPKDTLMISRTPEVTCVVVDVSHEDPEVKFNWYVDGVEVHNAKTKPREE
QYNSTYRVVSVLTVLHQDWLNGKEYKCKVSNKGLPSSIEKTISKAKGQPREPQVYTLPPSRDELTKNQVS
LTCLVKGFYPSDIAVEWESNGQPENNYKTTPPVLDSDGSFFLYSKLTVDKSRWQQGNVFSCSVMHEALHN
HYTQKSLSLSPGK**ITIFITLFLLSVCYSATVTFFKVKWIFSSVVDLKQTIIPDYRNMIGQGA**

>C*16:01-Final

**MSRSVALAVLALLSLSGLEA**IQRTPKIQVYSRHPAENGKSNFLNCYVSGFHPSDIEVDLLKNGERIEKVE
HSDLSFSKDWSFYLLYYTEFTPTEKDEYACRVNHVTLSQPKIVKWGKSYILL**GGGGSGGGGSGGGGS**CSH
SMRYFYTAVSRPGRGEPRFIAVGYVDDTQFVRFDSDAASPRGEPRAPWVEQEGPEYWDRETQKYKRQAQT
DRVSLRNLRGYYNQSEAGSHTLQWMYGCDLGPDGRLLRGYDQSAYDGKDYIALNEDLRSWTAADTAAQIT
QRKWEAARAAEQQRAYLEGTCVEWLRRYLENGKETLQRAEHPKTHVTHHLVSDHEATLRCWALGFYPAEI
TLTWQRDGEDQTQDTELVETRPAGDGTFQKWAAVVVPSGEEQRYTCHVQHEGLPEPLTLRWEPKSCDKTH

TCPPCPAPELLGGPSVFLFPPKPKDTLMISRTPEVTCVVVDVSHEDPEVKFNWYVDGVEVHNAKTKPREE
QYNSTYRVVSVLTVLHQDWLNGKEYKCKVSNKGLPSSIEKTISKAKGQPREPQVYTLPPSRDELTKNQVS
LTCLVKGFYPSDIAVEWESNGQPENNYKTTPPVLDSDGSFFLYSKLTVDKSRWQQGNVFSCSVMHEALHN
HYTQKSLSLSPGK**ITIFITLFLLSVCYSATVTFFKVKWIFSSVVDLKQTIIPDYRNMIGQGA**

>C*17:01-Final

**MSRSVALAVLALLSLSGLEA**IQRTPKIQVYSRHPAENGKSNFLNCYVSGFHPSDIEVDLLKNGERIEKVE
HSDLSFSKDWSFYLLYYTEFTPTEKDEYACRVNHVTLSQPKIVKWGKSYILL**GGGGSGGGGSGGGGS**GSH
SMRYFYTAVSRPGRGEPRFIAVGYVDDTQFVRFDSDAASPRGEPRAPWVEQEGPEYWDRETQKYKRQAQA
DRVNLRKLRGYYNQSEAGSHTIQRMYGCDLGPDGRLLRGYNQFAYDGKDYIALNEDLRSWTAADTAAQIS
QRKLEAAREAEQLRAYLEGECVEWLRGYLENGKETLQRAERPKTHVTHHPVSDHEATLRCWALGFYPAEI
TLTWQRDGEDQTQDTELVETRPAGDGTFQKWAAVVVPSGQEQRYTCHVQHEGLQEPCTLRWEPKSCDKTH
TCPPCPAPELLGGPSVFLFPPKPKDTLMISRTPEVTCVVVDVSHEDPEVKFNWYVDGVEVHNAKTKPREE
QYNSTYRVVSVLTVLHQDWLNGKEYKCKVSNKGLPSSIEKTISKAKGQPREPQVYTLPPSRDELTKNQVS
LTCLVKGFYPSDIAVEWESNGQPENNYKTTPPVLDSDGSFFLYSKLTVDKSRWQQGNVFSCSVMHEALHN
HYTQKSLSLSPGK**ITIFITLFLLSVCYSATVTFFKVKWIFSSVVDLKQTIIPDYRNMIGQGA**

>C*18:02-Final

**MSRSVALAVLALLSLSGLEA**IQRTPKIQVYSRHPAENGKSNFLNCYVSGFHPSDIEVDLLKNGERIEKVE
HSDLSFSKDWSFYLLYYTEFTPTEKDEYACRVNHVTLSQPKIVKWGKSYILL**GGGGSGGGGSGGGGS**CSH
SMRYFDTAVSRPGRGEPRFISVGYVDDTQFVRFDSDAASPRGEPRAPWVEQEGPEYWDRETQKYKRQAQA
DRVNLRKLRGYYNQSEDGSHTLQRMFGCDLGPDGRLLRGYNQFAYDGKDYIALNEDLRSWTAADTAAQIT
QRKWEAAREAEQRRAYLEGTCVEWLRRYLENGKETLQRAEHPKTHVTHHPVSDHEATLRCWALGFYPAEI
TLTWQWDGEDQTQDTELVETRPAGDGTFQKWAAVVVPSGEEQRYTCHVQHEGLPEPLTLRWEPKSCDKTH
TCPPCPAPELLGGPSVFLFPPKPKDTLMISRTPEVTCVVVDVSHEDPEVKFNWYVDGVEVHNAKTKPREE
QYNSTYRVVSVLTVLHQDWLNGKEYKCKVSNKGLPSSIEKTISKAKGQPREPQVYTLPPSRDELTKNQVS
LTCLVKGFYPSDIAVEWESNGQPENNYKTTPPVLDSDGSFFLYSKLTVDKSRWQQGNVFSCSVMHEALHN
HYTQKSLSLSPGK**ITIFITLFLLSVCYSATVTFFKVKWIFSSVVDLKQTIIPDYRNMIGQGA**

HLA Class II –

DRB – alpha chain is always DRA*01:01P. Use signal sequence of this chain so select exon 1, 2 and 3 of this chain i.e. total 203 aa.

**MAISGVPVLGFFIIAVLMSAQESWAIK**EEHVIIQAEFYLNPDQSGEFMFDFDGDEIFHVDMAKKETVWRL
EEFGRFASFEAQGALANIAVDKANLEIMTKRSNYTPITNVPPEVTVLTNSPVELREPNVLICFIDKFTPP
VVNVTWLRNGKPVTTGVSETVFLPREDHLFRKFHYLPFLPSTEDVYDCRVEHWGLDEPLLKHW

For DRB1 – first select 217 aa (exon1,2 and 3 then delete 33 aa for exon 1)

>DRB1*01:01(DR1)

**MAISGVPVLGFFIIAVLMSAQESWAIK**EEHVIIQAEFYLNPDQSGEFMFDFDGDEIFHVDMAKKETVWRL
EEFGRFASFEAQGALANIAVDKANLEIMTKRSNYTPITNVPPEVTVLTNSPVELREPNVLICFIDKFTPP
VVNVTWLRNGKPVTTGVSETVFLPREDHLFRKFHYLPFLPSTEDVYDCRVEHWGLDEPLLKHW**GGGGSGG**
**GGSGGGGS**PRFLWQLKFECHFFNGTERVRLLERCIYNQEESVRFDSDVGEYRAVTELGRPDAEYWNSQKD
LLEQRRAAVDTYCRHNYGVGESFTVQRRVEPKVTVYPSKTQPLQHHNLLVCSVSGFYPGSIEVRWFRNGQ
EEKAGVVSTGLIQNGDWTFQTLVMLETVPRSGEVYTCQVEHPSVTSPLTVEWEPKSCDKTHTCPPCPAPE
LLGGPSVFLFPPKPKDTLMISRTPEVTCVVVDVSHEDPEVKFNWYVDGVEVHNAKTKPREEQYNSTYRVV
SVLTVLHQDWLNGKEYKCKVSNKGLPSSIEKTISKAKGQPREPQVYTLPPSRDELTKNQVSLTCLVKGFY
PSDIAVEWESNGQPENNYKTTPPVLDSDGSFFLYSKLTVDKSRWQQGNVFSCSVMHEALHNHYTQKSLSL
SPGK**ITIFITLFLLSVCYSATVTFFKVKWIFSSVVDLKQTIIPDYRNMIGQGA**

>DRB1*01:02(DR1)

**MAISGVPVLGFFIIAVLMSAQESWAIK**EEHVIIQAEFYLNPDQSGEFMFDFDGDEIFHVDMAKKETVWRL
EEFGRFASFEAQGALANIAVDKANLEIMTKRSNYTPITNVPPEVTVLTNSPVELREPNVLICFIDKFTPP
VVNVTWLRNGKPVTTGVSETVFLPREDHLFRKFHYLPFLPSTEDVYDCRVEHWGLDEPLLKHW**GGGGSGG**
**GGSGGGGS**PRFLWQLKFECHFFNGTERVRLLERCIYNQEESVRFDSDVGEYRAVTELGRPDAEYWNSQKD
LLEQRRAAVDTYCRHNYGAVESFTVQRRVEPKVTVYPSKTQPLQHHNLLVCSVSGFYPGSIEVRWFRNGQ
EEKAGVVSTGLIQNGDWTFQTLVMLETVPRSGEVYTCQVEHPSVTSPLTVEWEPKSCDKTHTCPPCPAPE
LLGGPSVFLFPPKPKDTLMISRTPEVTCVVVDVSHEDPEVKFNWYVDGVEVHNAKTKPREEQYNSTYRVV
SVLTVLHQDWLNGKEYKCKVSNKGLPSSIEKTISKAKGQPREPQVYTLPPSRDELTKNQVSLTCLVKGFY
PSDIAVEWESNGQPENNYKTTPPVLDSDGSFFLYSKLTVDKSRWQQGNVFSCSVMHEALHNHYTQKSLSL
SPGK**ITIFITLFLLSVCYSATVTFFKVKWIFSSVVDLKQTIIPDYRNMIGQGA**

>DRB1*04:01(DR4)

**MAISGVPVLGFFIIAVLMSAQESWAIK**EEHVIIQAEFYLNPDQSGEFMFDFDGDEIFHVDMAKKETVWRL
EEFGRFASFEAQGALANIAVDKANLEIMTKRSNYTPITNVPPEVTVLTNSPVELREPNVLICFIDKFTPP
VVNVTWLRNGKPVTTGVSETVFLPREDHLFRKFHYLPFLPSTEDVYDCRVEHWGLDEPLLKHW**GGGGSGG**
**GGSGGGGS**PRFLEQVKHECHFFNGTERVRFLDRYFYHQEEYVRFDSDVGEYRAVTELGRPDAEYWNSQKD
LLEQKRAAVDTYCRHNYGVGESFTVQRRVYPEVTVYPAKTQPLQHHNLLVCSVNGFYPGSIEVRWFRNGQ
EEKTGVVSTGLIQNGDWTFQTLVMLETVPRSGEVYTCQVEHPSLTSPLTVEWEPKSCDKTHTCPPCPAPE
LLGGPSVFLFPPKPKDTLMISRTPEVTCVVVDVSHEDPEVKFNWYVDGVEVHNAKTKPREEQYNSTYRVV
SVLTVLHQDWLNGKEYKCKVSNKGLPSSIEKTISKAKGQPREPQVYTLPPSRDELTKNQVSLTCLVKGFY
PSDIAVEWESNGQPENNYKTTPPVLDSDGSFFLYSKLTVDKSRWQQGNVFSCSVMHEALHNHYTQKSLSL
SPGK**ITIFITLFLLSVCYSATVTFFKVKWIFSSVVDLKQTIIPDYRNMIGQGA**

>DRB1*04:02(DR4)

**MAISGVPVLGFFIIAVLMSAQESWAIK**EEHVIIQAEFYLNPDQSGEFMFDFDGDEIFHVDMAKKETVWRL
EEFGRFASFEAQGALANIAVDKANLEIMTKRSNYTPITNVPPEVTVLTNSPVELREPNVLICFIDKFTPP
VVNVTWLRNGKPVTTGVSETVFLPREDHLFRKFHYLPFLPSTEDVYDCRVEHWGLDEPLLKHW**GGGGSGG**

**GGSGGGGS**PRFLEQVKHECHFFNGTERVRFLDRYFYHQEEYVRFDSDVGEYRAVTELGRPDAEYWNSQKD
ILEDERAAVDTYCRHNYGVVESFTVQRRVYPEVTVYPAKTQPLQHHNLLVCSVNGFYPGSIEVRWFRNGQ
EEKTGVVSTGLIQNGDWTFQTLVMLETVPRSGEVYTCQVEHPSLTSPLTVEW
EPKSCDKTHTCPPCPAPELLGGPSVFLFPPKPKDTLMISRTPEVTCVVVDVSHEDPEVKFNWYVDGVEVH
NAKTKPREEQYNSTYRVVSVLTVLHQDWLNGKEYKCKVSNKGLPSSIEKTISKAKGQPREPQVYTLPPSR
DELTKNQVSLTCLVKGFYPSDIAVEWESNGQPENNYKTTPPVLDSDGSFFLYSKLTVDKSRWQQGNVFSC
SVMHEALHNHYTQKSLSLSPGK**ITIFITLFLLSVCYSATVTFFKVKWIFSSVVDLKQTIIPDYRNMIGQG
A**

>DRB1*04:03(DR4)

**MAISGVPVLGFFIIAVLMSAQESWAIK**EEHVIIQAEFYLNPDQSGEFMFDFDGDEIFHVDMAKKETVWRL
EEFGRFASFEAQGALANIAVDKANLEIMTKRSNYTPITNVPPEVTVLTNSPVELREPNVLICFIDKFTPP
VVNVTWLRNGKPVTTGVSETVFLPREDHLFRKFHYLPFLPSTEDVYDCRVEHWGLDEPLLKHW**GGGGSGG
GGSGGGGS**PRFLEQVKHECHFFNGTERVRFLDRYFYHQEEYVRFDSDVGEYRAVTELGRPDAEYWNSQKD
LLEQRRAEVDTYCRHNYGVVESFTVQRRVYPEVTVYPAKTQPLQHHNLLVCSVNGFYPGSIEVRWFRNGQ
EEKTGVVSTGLIQNGDWTFQTLVMLETVPRSGEVYTCQVEHPSLTSPLTVEW
EPKSCDKTHTCPPCPAPELLGGPSVFLFPPKPKDTLMISRTPEVTCVVVDVSHEDPEVKFNWYVDGVEVH
NAKTKPREEQYNSTYRVVSVLTVLHQDWLNGKEYKCKVSNKGLPSSIEKTISKAKGQPREPQVYTLPPSR
DELTKNQVSLTCLVKGFYPSDIAVEWESNGQPENNYKTTPPVLDSDGSFFLYSKLTVDKSRWQQGNVFSC
SVMHEALHNHYTQKSLSLSPGK**ITIFITLFLLSVCYSATVTFFKVKWIFSSVVDLKQTIIPDYRNMIGQG
A**

>DRB1*04:04(DR4)

**MAISGVPVLGFFIIAVLMSAQESWAIK**EEHVIIQAEFYLNPDQSGEFMFDFDGDEIFHVDMAKKETVWRL
EEFGRFASFEAQGALANIAVDKANLEIMTKRSNYTPITNVPPEVTVLTNSPVELREPNVLICFIDKFTPP
VVNVTWLRNGKPVTTGVSETVFLPREDHLFRKFHYLPFLPSTEDVYDCRVEHWGLDEPLLKHW**GGGGSGG
GGSGGGGS**PRFLEQVKHECHFFNGTERVRFLDRYFYHQEEYVRFDSDVGEYRAVTELGRPDAEYWNSQKD
LLEQRRAAVDTYCRHNYGVVESFTVQRRVYPEVTVYPAKTQPLQHHNLLVCSVNGFYPGSIEVRWFRNGQ
EEKTGVVSTGLIQNGDWTFQTLVMLETVPRSGEVYTCQVEHPSLTSPLTVEW
EPKSCDKTHTCPPCPAPELLGGPSVFLFPPKPKDTLMISRTPEVTCVVVDVSHEDPEVKFNWYVDGVEVH
NAKTKPREEQYNSTYRVVSVLTVLHQDWLNGKEYKCKVSNKGLPSSIEKTISKAKGQPREPQVYTLPPSR
DELTKNQVSLTCLVKGFYPSDIAVEWESNGQPENNYKTTPPVLDSDGSFFLYSKLTVDKSRWQQGNVFSC
SVMHEALHNHYTQKSLSLSPGK**ITIFITLFLLSVCYSATVTFFKVKWIFSSVVDLKQTIIPDYRNMIGQG
A**

>DRB1*04:05(DR4)

**MAISGVPVLGFFIIAVLMSAQESWAIK**EEHVIIQAEFYLNPDQSGEFMFDFDGDEIFHVDMAKKETVWRL
EEFGRFASFEAQGALANIAVDKANLEIMTKRSNYTPITNVPPEVTVLTNSPVELREPNVLICFIDKFTPP
VVNVTWLRNGKPVTTGVSETVFLPREDHLFRKFHYLPFLPSTEDVYDCRVEHWGLDEPLLKHW**GGGGSGG
GGSGGGGS**PRFLEQVKHECHFFNGTERVRFLDRYFYHQEEYVRFDSDVGEYRAVTELGRPSAEYWNSQKD
LLEQRRAAVDTYCRHNYGVGESFTVQRRVYPEVTVYPAKTQPLQHHNLLVCSVNGFYPGSIEVRWFRNGQ
EEKTGVVSTGLIQNGDWTFQTLVMLETVPRSGEVYTCQVEHPSLTSPLTVEW

EPKSCDKTHTCPPCPAPELLGGPSVFLFPPKPKDTLMISRTPEVTCVVVDVSHEDPEVKFNWYVDGVEVH
NAKTKPREEQYNSTYRVVSVLTVLHQDWLNGKEYKCKVSNKGLPSSIEKTISKAKGQPREPQVYTLPPSR
DELTKNQVSLTCLVKGFYPSDIAVEWESNGQPENNYKTTPPVLDSDGSFFLYSKLTVDKSRWQQGNVFSC
SVMHEALHNHYTQKSLSLSPGK**ITIFITLFLLSVCYSATVTFFKVKWIFSSVVDLKQTIIPDYRNMIGQG
A**

>DRB1*07:01(DR7)

**MAISGVPVLGFFIIAVLMSAQESWAIK**EEHVIIQAEFYLNPDQSGEFMFDFDGDEIFHVDMAKKETVWRL
EEFGRFASFEAQGALANIAVDKANLEIMTKRSNYTPITNVPPEVTVLTNSPVELREPNVLICFIDKFTPP
VVNVTWLRNGKPVTTGVSETVFLPREDHLFRKFHYLPFLPSTEDVYDCRVEHWGLDEPLLKHW**GGGGSGG
GGSGGGGS**PRFLWQGKYKCHFFNGTERVQFLERLFYNQEEFVRFDSDVGEYRAVTELGRPVAESWNSQKD
ILEDRRGQVDTVCRHNYGVGESFTVQRRVHPEVTVYPAKTQPLQHHNLLVCSVSGFYPGSIEVRWFRNGQ
EEKAGVVSTGLIQNGDWTFQTLVMLETVPRSGEVYTCQVEHPSVMSPLTVEW
EPKSCDKTHTCPPCPAPELLGGPSVFLFPPKPKDTLMISRTPEVTCVVVDVSHEDPEVKFNWYVDGVEVH
NAKTKPREEQYNSTYRVVSVLTVLHQDWLNGKEYKCKVSNKGLPSSIEKTISKAKGQPREPQVYTLPPSR
DELTKNQVSLTCLVKGFYPSDIAVEWESNGQPENNYKTTPPVLDSDGSFFLYSKLTVDKSRWQQGNVFSC
SVMHEALHNHYTQKSLSLSPGK**ITIFITLFLLSVCYSATVTFFKVKWIFSSVVDLKQTIIPDYRNMIGQG
A**

>DRB1*08:01(DR8)

**MAISGVPVLGFFIIAVLMSAQESWAIK**EEHVIIQAEFYLNPDQSGEFMFDFDGDEIFHVDMAKKETVWRL
EEFGRFASFEAQGALANIAVDKANLEIMTKRSNYTPITNVPPEVTVLTNSPVELREPNVLICFIDKFTPP
VVNVTWLRNGKPVTTGVSETVFLPREDHLFRKFHYLPFLPSTEDVYDCRVEHWGLDEPLLKHW**GGGGSGG
GGSGGGGS**PRFLEYSTGECYFFNGTERVRFLDRYFYNQEEYVRFDSDVGEYRAVTELGRPSAEYWNSQKD
FLEDRRALVDTYCRHNYGVGESFTVQRRVHPKVTVYPSKTQPLQHHNLLVCSVSGFYPGSIEVRWFRNGQ
EEKTGVVSTGLIHNGDWTFQTLVMLETVPRSGEVYTCQVEHPSVTSPLTVEW
EPKSCDKTHTCPPCPAPELLGGPSVFLFPPKPKDTLMISRTPEVTCVVVDVSHEDPEVKFNWYVDGVEVH
NAKTKPREEQYNSTYRVVSVLTVLHQDWLNGKEYKCKVSNKGLPSSIEKTISKAKGQPREPQVYTLPPSR
DELTKNQVSLTCLVKGFYPSDIAVEWESNGQPENNYKTTPPVLDSDGSFFLYSKLTVDKSRWQQGNVFSC
SVMHEALHNHYTQKSLSLSPGK**ITIFITLFLLSVCYSATVTFFKVKWIFSSVVDLKQTIIPDYRNMIGQG
A**

>DRB1*09:01(DR9)

**MAISGVPVLGFFIIAVLMSAQESWAIK**EEHVIIQAEFYLNPDQSGEFMFDFDGDEIFHVDMAKKETVWRL
EEFGRFASFEAQGALANIAVDKANLEIMTKRSNYTPITNVPPEVTVLTNSPVELREPNVLICFIDKFTPP
VVNVTWLRNGKPVTTGVSETVFLPREDHLFRKFHYLPFLPSTEDVYDCRVEHWGLDEPLLKHW**GGGGSGG
GGSGGGGS**PRFLKQDKFECHFFNGTERVRYLHRGIYNQEENVRFDSDVGEYRAVTELGRPVAESWNSQKD
FLERRRAEVDTVCRHNYGVGESFTVQRRVHPEVTVYPAKTQPLQHHNLLVCSVSGFYPGSIEVRWFRNGQ
EEKAGVVSTGLIQNGDWTFQTLVMLETVPRSGEVYTCQVEHPSVMSPLTVEW
EPKSCDKTHTCPPCPAPELLGGPSVFLFPPKPKDTLMISRTPEVTCVVVDVSHEDPEVKFNWYVDGVEVH
NAKTKPREEQYNSTYRVVSVLTVLHQDWLNGKEYKCKVSNKGLPSSIEKTISKAKGQPREPQVYTLPPSR
DELTKNQVSLTCLVKGFYPSDIAVEWESNGQPENNYKTTPPVLDSDGSFFLYSKLTVDKSRWQQGNVFSC

SVMHEALHNHYTQKSLSLSPGK**ITIFITLFLLSVCYSATVTFFKVKWIFSSVVDLKQTIIPDYRNMIGQG A**

>DRB1*10:01(DR10)

**MAISGVPVLGFFIIAVLMSAQESWAIK**EEHVIIQAEFYLNPDQSGEFMFDFDGDEIFHVDMAKKETVWRL EEFGRFASFEAQGALANIAVDKANLEIMTKRSNYTPITNVPPEVTVLTNSPVELREPNVLICFIDKFTPP VVNVTWLRNGKPVTTGVSETVFLPREDHLFRKFHYLPFLPSTEDVYDCRVEHWGLDEPLLKHW**GGGGSGG GGSGGGGS**PRFLEEVKFECHFFNGTERVRLLERRVHNQEEYARYDSDVGEYRAVTELGRPDAEYWNSQKD LLERRRAAVDTYCRHNYGVGESFTVQRRVQPKVTVYPSKTQPLQHHNLLVCSVNGFYPGSIEVRWFRNGQ EEKTGVVSTGLIQNGDWTFQTLVMLETVPQSGEVYTCQVEHPSVMSPLTVEW EPKSCDKTHTCPPCPAPELLGGPSVFLFPPKPKDTLMISRTPEVTCVVVDVSHEDPEVKFNWYVDGVEVH NAKTKPREEQYNSTYRVVSVLTVLHQDWLNGKEYKCKVSNKGLPSSIEKTISKAKGQPREPQVYTLPPSR DELTKNQVSLTCLVKGFYPSDIAVEWESNGQPENNYKTTPPVLDSDGSFFLYSKLTVDKSRWQQGNVFSC SVMHEALHNHYTQKSLSLSPGK**ITIFITLFLLSVCYSATVTFFKVKWIFSSVVDLKQTIIPDYRNMIGQG A**

>DRB1*11:01(DR11)

**MAISGVPVLGFFIIAVLMSAQESWAIK**EEHVIIQAEFYLNPDQSGEFMFDFDGDEIFHVDMAKKETVWRL EEFGRFASFEAQGALANIAVDKANLEIMTKRSNYTPITNVPPEVTVLTNSPVELREPNVLICFIDKFTPP VVNVTWLRNGKPVTTGVSETVFLPREDHLFRKFHYLPFLPSTEDVYDCRVEHWGLDEPLLKHW**GGGGSGG GGSGGGGS**PRFLEYSTSECHFFNGTERVRFLDRYFYNQEEYVRFDSDVGEFRAVTELGRPDEEYWNSQKD FLEDRRAAVDTYCRHNYGVGESFTVQRRVHPKVTVYPSKTQPLQHHNLLVCSVSGFYPGSIEVRWFRNGQ EEKTGVVSTGLIHNGDWTFQTLVMLETVPRSGEVYTCQVEHPSVTSPLTVEW EPKSCDKTHTCPPCPAPELLGGPSVFLFPPKPKDTLMISRTPEVTCVVVDVSHEDPEVKFNWYVDGVEVH NAKTKPREEQYNSTYRVVSVLTVLHQDWLNGKEYKCKVSNKGLPSSIEKTISKAKGQPREPQVYTLPPSR DELTKNQVSLTCLVKGFYPSDIAVEWESNGQPENNYKTTPPVLDSDGSFFLYSKLTVDKSRWQQGNVFSC SVMHEALHNHYTQKSLSLSPGK**ITIFITLFLLSVCYSATVTFFKVKWIFSSVVDLKQTIIPDYRNMIGQG A**

>DRB1*11:04(DR11)

**MAISGVPVLGFFIIAVLMSAQESWAIK**EEHVIIQAEFYLNPDQSGEFMFDFDGDEIFHVDMAKKETVWRL EEFGRFASFEAQGALANIAVDKANLEIMTKRSNYTPITNVPPEVTVLTNSPVELREPNVLICFIDKFTPP VVNVTWLRNGKPVTTGVSETVFLPREDHLFRKFHYLPFLPSTEDVYDCRVEHWGLDEPLLKHW**GGGGSGG GGSGGGGS**PRFLEYSTSECHFFNGTERVRFLDRYFYNQEEYVRFDSDVGEFRAVTELGRPDEEYWNSQKD FLEDRRAAVDTYCRHNYGVVESFTVQRRVHPKVTVYPSKTQPLQHHNLLVCSVSGFYPGSIEVRWFRNGQ EEKTGVVSTGLIHNGDWTFQTLVMLETVPRSGEVYTCQVEHPSVTSPLTVEW EPKSCDKTHTCPPCPAPELLGGPSVFLFPPKPKDTLMISRTPEVTCVVVDVSHEDPEVKFNWYVDGVEVH NAKTKPREEQYNSTYRVVSVLTVLHQDWLNGKEYKCKVSNKGLPSSIEKTISKAKGQPREPQVYTLPPSR DELTKNQVSLTCLVKGFYPSDIAVEWESNGQPENNYKTTPPVLDSDGSFFLYSKLTVDKSRWQQGNVFSC SVMHEALHNHYTQKSLSLSPGK**ITIFITLFLLSVCYSATVTFFKVKWIFSSVVDLKQTIIPDYRNMIGQG A**

>DRB1*12:01(DR12)

**MAISGVPVLGFFIIAVLMSAQESWAIK**EEHVIIQAEFYLNPDQSGEFMFDFDGDEIFHVDMAKKETVWRL
EEFGRFASFEAQGALANIAVDKANLEIMTKRSNYTPITNVPPEVTVLTNSPVELREPNVLICFIDKFTPP
VVNVTWLRNGKPVTTGVSETVFLPREDHLFRKFHYLPFLPSTEDVYDCRVEHWGLDEPLLKHW**GGGGSGG**
**GGSGGGGS**PRFLEYSTGECYFFNGTERVRLLERHFHNQEELLRFDSDVGEFRAVTELGRPVAESWNSQKD
ILEDRRAAVDTYCRHNYGAVESFTVQRRVHPKVTVYPSKTQPLQHHNLLVCSVSGFYPGSIEVRWFRNGQ
EEKTGVVSTGLIHNGDWTFQTLVMLETVPRSGEVYTCQVEHPSVTSPLTVEW
EPKSCDKTHTCPPCPAPELLGGPSVFLFPPKPKDTLMISRTPEVTCVVVDVSHEDPEVKFNWYVDGVEVH
NAKTKPREEQYNSTYRVVSVLTVLHQDWLNGKEYKCKVSNKGLPSSIEKTISKAKGQPREPQVYTLPPSR
DELTKNQVSLTCLVKGFYPSDIAVEWESNGQPENNYKTTPPVLDSDGSFFLYSKLTVDKSRWQQGNVFSC
SVMHEALHNHYTQKSLSLSPGK**ITIFITLFLLSVCYSATVTFFKVKWIFSSVVDLKQTIIPDYRNMIGQG**
**A**

>DRB1*12:02(DR12)

**MAISGVPVLGFFIIAVLMSAQESWAIK**EEHVIIQAEFYLNPDQSGEFMFDFDGDEIFHVDMAKKETVWRL
EEFGRFASFEAQGALANIAVDKANLEIMTKRSNYTPITNVPPEVTVLTNSPVELREPNVLICFIDKFTPP
VVNVTWLRNGKPVTTGVSETVFLPREDHLFRKFHYLPFLPSTEDVYDCRVEHWGLDEPLLKHW**GGGGSGG**
**GGSGGGGS**PRFLEYSTGECYFFNGTERVRLLERHFHNQEELLRFDSDVGEFRAVTELGRPVAESWNSQKD
FLEDRRAAVDTYCRHNYGAVESFTVQRRVHPKVTVYPSKTQPLQHHNLLVCSVSGFYPGSIEVRWFRNGQ
EEKTGVVSTGLIHNGDWTFQTLVMLETVPRSGEVYTCQVEHPSVTSPLTVEW
EPKSCDKTHTCPPCPAPELLGGPSVFLFPPKPKDTLMISRTPEVTCVVVDVSHEDPEVKFNWYVDGVEVH
NAKTKPREEQYNSTYRVVSVLTVLHQDWLNGKEYKCKVSNKGLPSSIEKTISKAKGQPREPQVYTLPPSR
DELTKNQVSLTCLVKGFYPSDIAVEWESNGQPENNYKTTPPVLDSDGSFFLYSKLTVDKSRWQQGNVFSC
SVMHEALHNHYTQKSLSLSPGK**ITIFITLFLLSVCYSATVTFFKVKWIFSSVVDLKQTIIPDYRNMIGQG**
**A**

>DRB1*13:01(DR13)

**MAISGVPVLGFFIIAVLMSAQESWAIK**EEHVIIQAEFYLNPDQSGEFMFDFDGDEIFHVDMAKKETVWRL
EEFGRFASFEAQGALANIAVDKANLEIMTKRSNYTPITNVPPEVTVLTNSPVELREPNVLICFIDKFTPP
VVNVTWLRNGKPVTTGVSETVFLPREDHLFRKFHYLPFLPSTEDVYDCRVEHWGLDEPLLKHW**GGGGSGG**
**GGSGGGGS**PRFLEYSTSECHFFNGTERVRFLDRYFHNQEENVRFDSDVGEFRAVTELGRPDAEYWNSQKD
ILEDERAAVDTYCRHNYGVVESFTVQRRVHPKVTVYPSKTQPLQHHNLLVCSVSGFYPGSIEVRWFRNGQ
EEKTGVVSTGLIHNGDWTFQTLVMLETVPRSGEVYTCQVEHPSVTSPLTVEW
EPKSCDKTHTCPPCPAPELLGGPSVFLFPPKPKDTLMISRTPEVTCVVVDVSHEDPEVKFNWYVDGVEVH
NAKTKPREEQYNSTYRVVSVLTVLHQDWLNGKEYKCKVSNKGLPSSIEKTISKAKGQPREPQVYTLPPSR
DELTKNQVSLTCLVKGFYPSDIAVEWESNGQPENNYKTTPPVLDSDGSFFLYSKLTVDKSRWQQGNVFSC
SVMHEALHNHYTQKSLSLSPGK**ITIFITLFLLSVCYSATVTFFKVKWIFSSVVDLKQTIIPDYRNMIGQG**
**A**

>DRB1*13:03(DR13)

**MAISGVPVLGFFIIAVLMSAQESWAIK**EEHVIIQAEFYLNPDQSGEFMFDFDGDEIFHVDMAKKETVWRL
EEFGRFASFEAQGALANIAVDKANLEIMTKRSNYTPITNVPPEVTVLTNSPVELREPNVLICFIDKFTPP

VVNVTWLRNGKPVTTGVSETVFLPREDHLFRKFHYLPFLPSTEDVYDCRVEHWGLDEPLLKHW**GGGGSGG**
**GGSGGGGS**PRFLEYSTSECHFFNGTERVRFLDRYFYNQEEYVRFDSDVGEYRAVTELGRPSAEYWNSQKD
ILEDKRAAVDTYCRHNYGVGESFTVQRRVHPKVTVYPSKTQPLQHHNLLVCSVSGFYPGSIEVRWFRNGQ
EEKTGVVSTGLIHNGDWTFQTLVMLETVPRSGEVYTCQVEHPSVTSPLTVEW
EPKSCDKTHTCPPCPAPELLGGPSVFLFPPKPKDTLMISRTPEVTCVVVDVSHEDPEVKFNWYVDGVEVH
NAKTKPREEQYNSTYRVVSVLTVLHQDWLNGKEYKCKVSNKGLPSSIEKTISKAKGQPREPQVYTLPPSR
DELTKNQVSLTCLVKGFYPSDIAVEWESNGQPENNYKTTPPVLDSDGSFFLYSKLTVDKSRWQQGNVFSC
SVMHEALHNHYTQKSLSLSPGK**ITIFITLFLLSVCYSATVTFFKVKWIFSSVVDLKQTIIPDYRNMIGQG**
**A**

>DRB1*14:01(DR14)

**MAISGVPVLGFFIIAVLMSAQESWAIK**EEHVIIQAEFYLNPDQSGEFMFDFDGDEIFHVDMAKKETVWRL
EEFGRFASFEAQGALANIAVDKANLEIMTKRSNYTPITNVPPEVTVLTNSPVELREPNVLICFIDKFTPP
VVNVTWLRNGKPVTTGVSETVFLPREDHLFRKFHYLPFLPSTEDVYDCRVEHWGLDEPLLKHW**GGGGSGG**
**GGSGGGGS**PRFLEYSTSECHFFNGTERVRFLDRYFHNQEEFVRFDSDVGEYRAVTELGRPAAEHWNSQKD
LLERRRAEVDTYCRHNYGVVESFTVQRRVHPKVTVYPSKTQPLQHYNLLVCSVSGFYPGSIEVRWFRNGQ
EEKTGVVSTGLIHNGDWTFQTLVMLETVPRSGEVYTCQVEHPSVTSPLTVEW
EPKSCDKTHTCPPCPAPELLGGPSVFLFPPKPKDTLMISRTPEVTCVVVDVSHEDPEVKFNWYVDGVEVH
NAKTKPREEQYNSTYRVVSVLTVLHQDWLNGKEYKCKVSNKGLPSSIEKTISKAKGQPREPQVYTLPPSR
DELTKNQVSLTCLVKGFYPSDIAVEWESNGQPENNYKTTPPVLDSDGSFFLYSKLTVDKSRWQQGNVFSC
SVMHEALHNHYTQKSLSLSPGK**ITIFITLFLLSVCYSATVTFFKVKWIFSSVVDLKQTIIPDYRNMIGQG**
**A**

>DRB1*14:02(DR14)

**MAISGVPVLGFFIIAVLMSAQESWAIK**EEHVIIQAEFYLNPDQSGEFMFDFDGDEIFHVDMAKKETVWRL
EEFGRFASFEAQGALANIAVDKANLEIMTKRSNYTPITNVPPEVTVLTNSPVELREPNVLICFIDKFTPP
VVNVTWLRNGKPVTTGVSETVFLPREDHLFRKFHYLPFLPSTEDVYDCRVEHWGLDEPLLKHW**GGGGSGG**
**GGSGGGGS**PRFLEYSTSECHFFNGTERVRFLERYFHNQEENVRFDSDVGEYRAVTELGRPDAEYWNSQKD
LLEQRRAAVDTYCRHNYGVGESFTVQRRVHPKVTVYPSKTQPLQHHNLLVCSVSGFYPGSIEVRWFRNGQ
EEKTGVVSTGLIHNGDWTFQTLVMLETVPRSGEVYTCQVEHPSVTSPLTVEW
EPKSCDKTHTCPPCPAPELLGGPSVFLFPPKPKDTLMISRTPEVTCVVVDVSHEDPEVKFNWYVDGVEVH
NAKTKPREEQYNSTYRVVSVLTVLHQDWLNGKEYKCKVSNKGLPSSIEKTISKAKGQPREPQVYTLPPSR
DELTKNQVSLTCLVKGFYPSDIAVEWESNGQPENNYKTTPPVLDSDGSFFLYSKLTVDKSRWQQGNVFSC
SVMHEALHNHYTQKSLSLSPGK**ITIFITLFLLSVCYSATVTFFKVKWIFSSVVDLKQTIIPDYRNMIGQG**
**A**

>DRB1*14:54(DR14)

**MAISGVPVLGFFIIAVLMSAQESWAIK**EEHVIIQAEFYLNPDQSGEFMFDFDGDEIFHVDMAKKETVWRL
EEFGRFASFEAQGALANIAVDKANLEIMTKRSNYTPITNVPPEVTVLTNSPVELREPNVLICFIDKFTPP
VVNVTWLRNGKPVTTGVSETVFLPREDHLFRKFHYLPFLPSTEDVYDCRVEHWGLDEPLLKHW**GGGGSGG**
**GGSGGGGS**PRFLEYSTSECHFFNGTERVRFLDRYFHNQEEFVRFDSDVGEYRAVTELGRPAAEHWNSQKD
LLERRRAEVDTYCRHNYGVVESFTVQRRVHPKVTVYPSKTQPLQHHNLLVCSVSGFYPGSIEVRWFRNGQ

EEKTGVVSTGLIHNGDWTFQTLVMLETVPRSGEVYTCQVEHPSVTSPLTVEW
EPKSCDKTHTCPPCPAPELLGGPSVFLFPPKPKDTLMISRTPEVTCVVVDVSHEDPEVKFNWYVDGVEVH
NAKTKPREEQYNSTYRVVSVLTVLHQDWLNGKEYKCKVSNKGLPSSIEKTISKAKGQPREPQVYTLPPSR
DELTKNQVSLTCLVKGFYPSDIAVEWESNGQPENNYKTTPPVLDSDGSFFLYSKLTVDKSRWQQGNVFSC
SVMHEALHNHYTQKSLSLSPGK**ITIFITLFLLSVCYSATVTFFKVKWIFSSVVDLKQTIIPDYRNMIGQG**
**A**

>DRB1*15:01(DR15)

**MAISGVPVLGFFIIAVLMSAQESWAIK**EEHVIIQAEFYLNPDQSGEFMFDFDGDEIFHVDMAKKETVWRL
EEFGRFASFEAQGALANIAVDKANLEIMTKRSNYTPITNVPPEVTVLTNSPVELREPNVLICFIDKFTPP
VVNVTWLRNGKPVTTGVSETVFLPREDHLFRKFHYLPFLPSTEDVYDCRVEHWGLDEPLLKHW**GGGGSGG**
**GGSGGGGS**PRFLWQPKRECHFFNGTERVRFLDRYFYNQEESVRFDSDVGEFRAVTELGRPDAEYWNSQKD
ILEQARAAVDTYCRHNYGVVESFTVQRRVQPKVTVYPSKTQPLQHHNLLVCSVSGFYPGSIEVRWFLNGQ
EEKAGMVSTGLIQNGDWTFQTLVMLETVPRSGEVYTCQVEHPSVTSPLTVEW
EPKSCDKTHTCPPCPAPELLGGPSVFLFPPKPKDTLMISRTPEVTCVVVDVSHEDPEVKFNWYVDGVEVH
NAKTKPREEQYNSTYRVVSVLTVLHQDWLNGKEYKCKVSNKGLPSSIEKTISKAKGQPREPQVYTLPPSR
DELTKNQVSLTCLVKGFYPSDIAVEWESNGQPENNYKTTPPVLDSDGSFFLYSKLTVDKSRWQQGNVFSC
SVMHEALHNHYTQKSLSLSPGK**ITIFITLFLLSVCYSATVTFFKVKWIFSSVVDLKQTIIPDYRNMIGQG**
**A**

>DRB1*15:02(DR15)

**MAISGVPVLGFFIIAVLMSAQESWAIK**EEHVIIQAEFYLNPDQSGEFMFDFDGDEIFHVDMAKKETVWRL
EEFGRFASFEAQGALANIAVDKANLEIMTKRSNYTPITNVPPEVTVLTNSPVELREPNVLICFIDKFTPP
VVNVTWLRNGKPVTTGVSETVFLPREDHLFRKFHYLPFLPSTEDVYDCRVEHWGLDEPLLKHW**GGGGSGG**
**GGSGGGGS**PRFLWQPKRECHFFNGTERVRFLDRYFYNQEESVRFDSDVGEFRAVTELGRPDAEYWNSQKD
ILEQARAAVDTYCRHNYGVGESFTVQRRVQPKVTVYPSKTQPLQHHNLLVCSVSGFYPGSIEVRWFLNGQ
EEKAGMVSTGLIQNGDWTFQTLVMLETVPRSGEVYTCQVEHPSVTSPLTVEW
EPKSCDKTHTCPPCPAPELLGGPSVFLFPPKPKDTLMISRTPEVTCVVVDVSHEDPEVKFNWYVDGVEVH
NAKTKPREEQYNSTYRVVSVLTVLHQDWLNGKEYKCKVSNKGLPSSIEKTISKAKGQPREPQVYTLPPSR
DELTKNQVSLTCLVKGFYPSDIAVEWESNGQPENNYKTTPPVLDSDGSFFLYSKLTVDKSRWQQGNVFSC
SVMHEALHNHYTQKSLSLSPGK**ITIFITLFLLSVCYSATVTFFKVKWIFSSVVDLKQTIIPDYRNMIGQG**
**A**

>DRB1*15:03(DR15)

**MAISGVPVLGFFIIAVLMSAQESWAIK**EEHVIIQAEFYLNPDQSGEFMFDFDGDEIFHVDMAKKETVWRL
EEFGRFASFEAQGALANIAVDKANLEIMTKRSNYTPITNVPPEVTVLTNSPVELREPNVLICFIDKFTPP
VVNVTWLRNGKPVTTGVSETVFLPREDHLFRKFHYLPFLPSTEDVYDCRVEHWGLDEPLLKHW**GGGGSGG**
**GGSGGGGS**PRFLWQPKRECHFFNGTERVRFLDRHFYNQEESVRFDSDVGEFRAVTELGRPDAEYWNSQKD
ILEQARAAVDTYCRHNYGVVESFTVQRRVQPKVTVYPSKTQPLQHHNLLVCSVSGFYPGSIEVRWFLNGQ
EEKAGMVSTGLIQNGDWTFQTLVMLETVPRSGEVYTCQVEHPSVTSPLTVEW
EPKSCDKTHTCPPCPAPELLGGPSVFLFPPKPKDTLMISRTPEVTCVVVDVSHEDPEVKFNWYVDGVEVH
NAKTKPREEQYNSTYRVVSVLTVLHQDWLNGKEYKCKVSNKGLPSSIEKTISKAKGQPREPQVYTLPPSR

DELTKNQVSLTCLVKGFYPSDIAVEWESNGQPENNYKTTPPVLDSDGSFFLYSKLTVDKSRWQQGNVFSC
SVMHEALHNHYTQKSLSLSPGK**ITIFITLFLLSVCYSATVTFFKVKWIFSSVVDLKQTIIPDYRNMIGQG
A**

>DRB1*16:01(DR16)

**MAISGVPVLGFFIIAVLMSAQESWAIK**EEHVIIQAEFYLNPDQSGEFMFDFDGDEIFHVDMAKKETVWRL
EEFGRFASFEAQGALANIAVDKANLEIMTKRSNYTPITNVPPEVTVLTNSPVELREPNVLICFIDKFTPP
VVNVTWLRNGKPVTTGVSETVFLPREDHLFRKFHYLPFLPSTEDVYDCRVEHWGLDEPLLKHW**GGGGSGG
GGSGGGGS**PRFLWQPKRECHFFNGTERVRFLDRYFYNQEESVRFDSDVGEYRAVTELGRPDAEYWNSQKD
FLEDRRAAVDTYCRHNYGVGESFTVQRRVQPKVTVYPSKTQPLQHHNLLVCSVSGFYPGSIEVRWFLNGQ
EEKAGMVSTGLIQNGDWTFQTLVMLETVPRSGEVYTCQVEHPSVTSPLTVEW
EPKSCDKTHTCPPCPAPELLGGPSVFLFPPKPKDTLMISRTPEVTCVVVDVSHEDPEVKFNWYVDGVEVH
NAKTKPREEQYNSTYRVVSVLTVLHQDWLNGKEYKCKVSNKGLPSSIEKTISKAKGQPREPQVYTLPPSR
DELTKNQVSLTCLVKGFYPSDIAVEWESNGQPENNYKTTPPVLDSDGSFFLYSKLTVDKSRWQQGNVFSC
SVMHEALHNHYTQKSLSLSPGK**ITIFITLFLLSVCYSATVTFFKVKWIFSSVVDLKQTIIPDYRNMIGQG
A**

>DRB1*16:02(DR16)

**MAISGVPVLGFFIIAVLMSAQESWAIK**EEHVIIQAEFYLNPDQSGEFMFDFDGDEIFHVDMAKKETVWRL
EEFGRFASFEAQGALANIAVDKANLEIMTKRSNYTPITNVPPEVTVLTNSPVELREPNVLICFIDKFTPP
VVNVTWLRNGKPVTTGVSETVFLPREDHLFRKFHYLPFLPSTEDVYDCRVEHWGLDEPLLKHW**GGGGSGG
GGSGGGGS**PRFLWQPKRECHFFNGTERVRFLDRYFYNQEESVRFDSDVGEYRAVTELGRPDAEYWNSQKD
LLEDRRAAVDTYCRHNYGVGESFTVQRRVQPKVTVYPSKTQPLQHHNLLVCSVSGFYPGSIEVRWFLNGQ
EEKAGMVSTGLIQNGDWTFQTLVMLETVPRSGEVYTCQVEHPSVTSPLTVEW
EPKSCDKTHTCPPCPAPELLGGPSVFLFPPKPKDTLMISRTPEVTCVVVDVSHEDPEVKFNWYVDGVEVH
NAKTKPREEQYNSTYRVVSVLTVLHQDWLNGKEYKCKVSNKGLPSSIEKTISKAKGQPREPQVYTLPPSR
DELTKNQVSLTCLVKGFYPSDIAVEWESNGQPENNYKTTPPVLDSDGSFFLYSKLTVDKSRWQQGNVFSC
SVMHEALHNHYTQKSLSLSPGK**ITIFITLFLLSVCYSATVTFFKVKWIFSSVVDLKQTIIPDYRNMIGQG
A**

>DRB1*03:01(DR17)

**MAISGVPVLGFFIIAVLMSAQESWAIK**EEHVIIQAEFYLNPDQSGEFMFDFDGDEIFHVDMAKKETVWRL
EEFGRFASFEAQGALANIAVDKANLEIMTKRSNYTPITNVPPEVTVLTNSPVELREPNVLICFIDKFTPP
VVNVTWLRNGKPVTTGVSETVFLPREDHLFRKFHYLPFLPSTEDVYDCRVEHWGLDEPLLKHW**GGGGSGG
GGSGGGGS**PRFLEYSTSECHFFNGTERVRYLDRYFHNQEENVRFDSDVGEFRAVTELGRPDAEYWNSQKD
LLEQKRGRVDNYCRHNYGVVESFTVQRRVHPKVTVYPSKTQPLQHHNLLVCSVSGFYPGSIEVRWFRNGQ
EEKTGVVSTGLIHNGDWTFQTLVMLETVPRSGEVYTCQVEHPSVTSPLTVEW
EPKSCDKTHTCPPCPAPELLGGPSVFLFPPKPKDTLMISRTPEVTCVVVDVSHEDPEVKFNWYVDGVEVH
NAKTKPREEQYNSTYRVVSVLTVLHQDWLNGKEYKCKVSNKGLPSSIEKTISKAKGQPREPQVYTLPPSR
DELTKNQVSLTCLVKGFYPSDIAVEWESNGQPENNYKTTPPVLDSDGSFFLYSKLTVDKSRWQQGNVFSC
SVMHEALHNHYTQKSLSLSPGK**ITIFITLFLLSVCYSATVTFFKVKWIFSSVVDLKQTIIPDYRNMIGQG
A**

>DRB1*03:02(DR18)

**MAISGVPVLGFFIIAVLMSAQESWAIK**EEHVIIQAEFYLNPDQSGEFMFDFDGDEIFHVDMAKKETVWRL
EEFGRFASFEAQGALANIAVDKANLEIMTKRSNYTPITNVPPEVTVLTNSPVELREPNVLICFIDKFTPP
VVNVTWLRNGKPVTTGVSETVFLPREDHLFRKFHYLPFLPSTEDVYDCRVEHWGLDEPLLKHW**GGGGSGG**
**GGSGGGGS**PRFLEYSTSECHFFNGTERVRFLERYFHNQEENVRFDSDVGEYRAVTELGRPDAEYWNSQKD
LLEQKRGRVDNYCRHNYGVGESFTVQRRVHPKVTVYPSKTQPLQHHNLLVCSVSGFYPGSIEVRWFRNGQ
EEKTGVVSTGLIHNGDWTFQTLVMLETVPRSGEVYTCQVEHPSVTSPLTVEW
EPKSCDKTHTCPPCPAPELLGGPSVFLFPPKPKDTLMISRTPEVTCVVVDVSHEDPEVKFNWYVDGVEVH
NAKTKPREEQYNSTYRVVSVLTVLHQDWLNGKEYKCKVSNKGLPSSIEKTISKAKGQPREPQVYTLPPSR
DELTKNQVSLTCLVKGFYPSDIAVEWESNGQPENNYKTTPPVLDSDGSFFLYSKLTVDKSRWQQGNVFSC
SVMHEALHNHYTQKSLSLSPGK**ITIFITLFLLSVCYSATVTFFKVKWIFSSVVDLKQTIIPDYRNMIGQG**
**A**

>DRB5*01:01(DR51)

**MAISGVPVLGFFIIAVLMSAQESWAIK**EEHVIIQAEFYLNPDQSGEFMFDFDGDEIFHVDMAKKETVWRL
EEFGRFASFEAQGALANIAVDKANLEIMTKRSNYTPITNVPPEVTVLTNSPVELREPNVLICFIDKFTPP
VVNVTWLRNGKPVTTGVSETVFLPREDHLFRKFHYLPFLPSTEDVYDCRVEHWGLDEPLLKHW**GGGGSGG**
**GGSGGGGS**PRFLQQDKYECHFFNGTERVRFLHRDIYNQEEDLRFDSDVGEYRAVTELGRPDAEYWNSQKD
FLEDRRAAVDTYCRHNYGVGESFTVQRRVEPKVTVYPARTQTLQHHNLLVCSVNGFYPGSIEVRWFRNSQ
EEKAGVVSTGLIQNGDWTFQTLVMLETVPRSGEVYTCQVEHPSVTSPLTVEW
EPKSCDKTHTCPPCPAPELLGGPSVFLFPPKPKDTLMISRTPEVTCVVVDVSHEDPEVKFNWYVDGVEVH
NAKTKPREEQYNSTYRVVSVLTVLHQDWLNGKEYKCKVSNKGLPSSIEKTISKAKGQPREPQVYTLPPSR
DELTKNQVSLTCLVKGFYPSDIAVEWESNGQPENNYKTTPPVLDSDGSFFLYSKLTVDKSRWQQGNVFSC
SVMHEALHNHYTQKSLSLSPGK**ITIFITLFLLSVCYSATVTFFKVKWIFSSVVDLKQTIIPDYRNMIGQG**
**A**

>DRB5*02:02(DR51)

**MAISGVPVLGFFIIAVLMSAQESWAIK**EEHVIIQAEFYLNPDQSGEFMFDFDGDEIFHVDMAKKETVWRL
EEFGRFASFEAQGALANIAVDKANLEIMTKRSNYTPITNVPPEVTVLTNSPVELREPNVLICFIDKFTPP
VVNVTWLRNGKPVTTGVSETVFLPREDHLFRKFHYLPFLPSTEDVYDCRVEHWGLDEPLLKHW**GGGGSGG**
**GGSGGGGS**PCFLQQDKYECHFFNGTERVRFLHRGIYNQEENVRFDSDVGEYRAVTELGRPDAEYWNSQKD
ILEQARAAVDTYCRHNYGAVESFTVQRRVEPKVTVYPARTQTLQHHNLLVCSVNGFYPGSIEVRWFRNGQ
EEKAGVVSTGLIQNGDWTFQILVMLETVPRSGEVYTCQVEHPSVTSPLTVEW
EPKSCDKTHTCPPCPAPELLGGPSVFLFPPKPKDTLMISRTPEVTCVVVDVSHEDPEVKFNWYVDGVEVH
NAKTKPREEQYNSTYRVVSVLTVLHQDWLNGKEYKCKVSNKGLPSSIEKTISKAKGQPREPQVYTLPPSR
DELTKNQVSLTCLVKGFYPSDIAVEWESNGQPENNYKTTPPVLDSDGSFFLYSKLTVDKSRWQQGNVFSC
SVMHEALHNHYTQKSLSLSPGK**ITIFITLFLLSVCYSATVTFFKVKWIFSSVVDLKQTIIPDYRNMIGQG**
**A**

>DRB3*01:01(DR52)

**MAISGVPVLGFFIIAVLMSAQESWAIK**EEHVIIQAEFYLNPDQSGEFMFDFDGDEIFHVDMAKKETVWRL
EEFGRFASFEAQGALANIAVDKANLEIMTKRSNYTPITNVPPEVTVLTNSPVELREPNVLICFIDKFTPP
VVNVTWLRNGKPVTTGVSETVFLPREDHLFRKFHYLPFLPSTEDVYDCRVEHWGLDEPLLKHW**GGGGSGG
GGSGGGGS**PRFLELRKSECHFFNGTERVRYLDRYFHNQEEFLRFDSDVGEYRAVTELGRPVAESWNSQKD
LLEQKRGRVDNYCRHNYGVGESFTVQRRVHPQVTVYPAKTQPLQHHNLLVCSVSGFYPGSIEVRWFRNGQ
EEKAGVVSTGLIQNGDWTFQTLVMLETVPRSGEVYTCQVEHPSVTSALTVEW
EPKSCDKTHTCPPCPAPELLGGPSVFLFPPKPKDTLMISRTPEVTCVVVDVSHEDPEVKFNWYVDGVEVH
NAKTKPREEQYNSTYRVVSVLTVLHQDWLNGKEYKCKVSNKGLPSSIEKTISKAKGQPREPQVYTLPPSR
DELTKNQVSLTCLVKGFYPSDIAVEWESNGQPENNYKTTPPVLDSDGSFFLYSKLTVDKSRWQQGNVFSC
SVMHEALHNHYTQKSLSLSPGK**ITIFITLFLLSVCYSATVTFFKVKWIFSSVVDLKQTIIPDYRNMIGQG
A**

>DRB3*02:02(DR52)

**MAISGVPVLGFFIIAVLMSAQESWAIK**EEHVIIQAEFYLNPDQSGEFMFDFDGDEIFHVDMAKKETVWRL
EEFGRFASFEAQGALANIAVDKANLEIMTKRSNYTPITNVPPEVTVLTNSPVELREPNVLICFIDKFTPP
VVNVTWLRNGKPVTTGVSETVFLPREDHLFRKFHYLPFLPSTEDVYDCRVEHWGLDEPLLKHW**GGGGSGG
GGSGGGGS**PRFLELLKSECHFFNGTERVRFLERHFHNQEEYARFDSDVGEYRAVRELGRPDAEYWNSQKD
LLEQKRGQVDNYCRHNYGVGESFTVQRRVHPQVTVYPAKTQPLQHHNLLVCSVSGFYPGSIEVRWFRNGQ
EEKAGVVSTGLIQNGDWTFQTLVMLETVPRSGEVYTCQVEHPSVTSPLTVEW
EPKSCDKTHTCPPCPAPELLGGPSVFLFPPKPKDTLMISRTPEVTCVVVDVSHEDPEVKFNWYVDGVEVH
NAKTKPREEQYNSTYRVVSVLTVLHQDWLNGKEYKCKVSNKGLPSSIEKTISKAKGQPREPQVYTLPPSR
DELTKNQVSLTCLVKGFYPSDIAVEWESNGQPENNYKTTPPVLDSDGSFFLYSKLTVDKSRWQQGNVFSC
SVMHEALHNHYTQKSLSLSPGK**ITIFITLFLLSVCYSATVTFFKVKWIFSSVVDLKQTIIPDYRNMIGQG
A**

>DRB3*03:01(DR52)

**MAISGVPVLGFFIIAVLMSAQESWAIK**EEHVIIQAEFYLNPDQSGEFMFDFDGDEIFHVDMAKKETVWRL
EEFGRFASFEAQGALANIAVDKANLEIMTKRSNYTPITNVPPEVTVLTNSPVELREPNVLICFIDKFTPP
VVNVTWLRNGKPVTTGVSETVFLPREDHLFRKFHYLPFLPSTEDVYDCRVEHWGLDEPLLKHW**GGGGSGG
GGSGGGGS**PRFLELLKSECHFFNGTERVRFLERYFHNQEEFVRFDSDVGEYRAVTELGRPVAESWNSQKD
LLEQKRGQVDNYCRHNYGVVESFTVQRRVHPQVTVYPAKTQPLQHHNLLVCSVSGFYPGSIEVRWFRNGQ
EEKTGVVSTGLIHNGDWTFQTLVMLETVPRSGEVYTCQVEHPSVTSPLTVEW
EPKSCDKTHTCPPCPAPELLGGPSVFLFPPKPKDTLMISRTPEVTCVVVDVSHEDPEVKFNWYVDGVEVH
NAKTKPREEQYNSTYRVVSVLTVLHQDWLNGKEYKCKVSNKGLPSSIEKTISKAKGQPREPQVYTLPPSR
DELTKNQVSLTCLVKGFYPSDIAVEWESNGQPENNYKTTPPVLDSDGSFFLYSKLTVDKSRWQQGNVFSC
SVMHEALHNHYTQKSLSLSPGK**ITIFITLFLLSVCYSATVTFFKVKWIFSSVVDLKQTIIPDYRNMIGQG
A**

>DRB4*01:01(DR53)

**MAISGVPVLGFFIIAVLMSAQESWAIK**EEHVIIQAEFYLNPDQSGEFMFDFDGDEIFHVDMAKKETVWRL
EEFGRFASFEAQGALANIAVDKANLEIMTKRSNYTPITNVPPEVTVLTNSPVELREPNVLICFIDKFTPP
VVNVTWLRNGKPVTTGVSETVFLPREDHLFRKFHYLPFLPSTEDVYDCRVEHWGLDEPLLKHW**GGGGSGG**

**GGSGGGGS**PRFLEQAKCECHFLNGTERVWNLIRYIYNQEEYARYNSDLGEYQAVTELGRPDAEYWNSQKD
LLERRRAEVDTYCRYNYGVVESFTVQRRVQPKVTVYPSKTQPLQHHNLLVCSVNGFYPGSIEVRWFRNSQ
EEKAGVVSTGLIQNGDWTFQTLVMLETVPRSGEVYTCQVEHPSMMSPLTVQW
EPKSCDKTHTCPPCPAPELLGGPSVFLFPPKPKDTLMISRTPEVTCVVVDVSHEDPEVKFNWYVDGVEVH
NAKTKPREEQYNSTYRVVSVLTVLHQDWLNGKEYKCKVSNKGLPSSIEKTISKAKGQPREPQVYTLPPSR
DELTKNQVSLTCLVKGFYPSDIAVEWESNGQPENNYKTTPPVLDSDGSFFLYSKLTVDKSRWQQGNVFSC
SVMHEALHNHYTQKSLSLSPGK**ITIFITLFLLSVCYSATVTFFKVKWIFSSVVDLKQTIIPDYRNMIGQG
A**

>DRB4*01:03(DR53)

**MAISGVPVLGFFIIAVLMSAQESWAIK**EEHVIIQAEFYLNPDQSGEFMFDFDGDEIFHVDMAKKETVWRL
EEFGRFASFEAQGALANIAVDKANLEIMTKRSNYTPITNVPPEVTVLTNSPVELREPNVLICFIDKFTPP
VVNVTWLRNGKPVTTGVSETVFLPREDHLFRKFHYLPFLPSTEDVYDCRVEHWGLDEPLLKHW**GGGGSGG
GGSGGGGS**PRFLEQAKCECHFLNGTERVWNLIRYIYNQEEYARYNSDLGEYQAVTELGRPDAEYWNSQKD
LLERRRAEVDTYCRYNYGVVESFTVQRRVQPKVTVYPSKTQPLQHHNLLVCSVNGFYPGSIEVRWFRNGQ
EEKAGVVSTGLIQNGDWTFQTLVMLETVPRSGEVYTCQVEHPSMMSPLTVQW
EPKSCDKTHTCPPCPAPELLGGPSVFLFPPKPKDTLMISRTPEVTCVVVDVSHEDPEVKFNWYVDGVEVH
NAKTKPREEQYNSTYRVVSVLTVLHQDWLNGKEYKCKVSNKGLPSSIEKTISKAKGQPREPQVYTLPPSR
DELTKNQVSLTCLVKGFYPSDIAVEWESNGQPENNYKTTPPVLDSDGSFFLYSKLTVDKSRWQQGNVFSC
SVMHEALHNHYTQKSLSLSPGK**ITIFITLFLLSVCYSATVTFFKVKWIFSSVVDLKQTIIPDYRNMIGQG
A**

>DRB1*01:03(DR103)

**MAISGVPVLGFFIIAVLMSAQESWAIK**EEHVIIQAEFYLNPDQSGEFMFDFDGDEIFHVDMAKKETVWRL
EEFGRFASFEAQGALANIAVDKANLEIMTKRSNYTPITNVPPEVTVLTNSPVELREPNVLICFIDKFTPP
VVNVTWLRNGKPVTTGVSETVFLPREDHLFRKFHYLPFLPSTEDVYDCRVEHWGLDEPLLKHW**GGGGSGG
GGSGGGGS**PRFLWQLKFECHFFNGTERVRLLERCIYNQEESVRFDSDVGEYRAVTELGRPDAEYWNSQKD
ILEDERAAVDTYCRHNYGVGESFTVQRRVEPKVTVYPSKTQPLQHHNLLVCSVSGFYPGSIEVRWFRNGQ
EEKAGVVSTGLIQNGDWTFQTLVMLETVPRSGEVYTCQVEHPSVTSPLTVEW
EPKSCDKTHTCPPCPAPELLGGPSVFLFPPKPKDTLMISRTPEVTCVVVDVSHEDPEVKFNWYVDGVEVH
NAKTKPREEQYNSTYRVVSVLTVLHQDWLNGKEYKCKVSNKGLPSSIEKTISKAKGQPREPQVYTLPPSR
DELTKNQVSLTCLVKGFYPSDIAVEWESNGQPENNYKTTPPVLDSDGSFFLYSKLTVDKSRWQQGNVFSC
SVMHEALHNHYTQKSLSLSPGK**ITIFITLFLLSVCYSATVTFFKVKWIFSSVVDLKQTIIPDYRNMIGQG
A**

HLA DQ

>DQA1*02:01-DQB1*02:01(DQ2)

**MILNKALMLGALALTTVMSPCGGEDIV**ADHVASYGVNLYQSYGPSGQFTHEFDGDEEFYVDLERKETVWK
LPLFHRLRFDPQFALTNIAVLKHNLNILIKRSNSTAATNEVPEVTVFSKSPVTLGQPNTLICLVDNIFPP
VVNITWLSNGHSVTEGVSETSFLSKSDHSFFKISYLTFLPSADEIYDCKVEHWGLDEPLLKHW**GGGGSGG
GGSGGGGS**EDFVYQFKGMCYFTNGTERVRLVSRSIYNREEIVRFDSDVGEFRAVTLLGLPAAEYWNSQKD
ILERKRAAVDRVCRHNYQLELRTTLQRRVEPTVTISPSRTEALNHHNLLVCSVTDFYPAQIKVRWFRNDQ

EETAGVVSTPLIRNGDWTFQILVMLEMTPQRGDVYTCHVEHPSLQSPITVEWEPKSCDKTHTCPPCPAPE
LLGGPSVFLFPPKPKDTLMISRTPEVTCVVVDVSHEDPEVKFNWYVDGVEVHNAKTKPREEQYNSTYRVV
SVLTVLHQDWLNGKEYKCKVSNKGLPSSIEKTISKAKGQPREPQVYTLPPSRDELTKNQVSLTCLVKGFY
PSDIAVEWESNGQPENNYKTTPPVLDSDGSFFLYSKLTVDKSRWQQGNVFSCSVMHEALHNHYTQKSLSL
SPGK**ITIFITLFLLSVCYSATVTFFKVKWIFSSVVDLKQTIIPDYRNMIGQGA**

>DQA1*03:01-DQB1*02:01(DQ2)

**MILNKALMLGALALTTVMSPCGGEDIV**ADHVASYGVNLYQSYGPSGQYSHEFDGDEEFYVDLERKETVWQ
LPLFRRFRRFDPQFALTNIAVLKHNLNIVIKRSNSTAATNEVPEVTVFSKSPVTLGQPNTLICLVDNIFP
PVVNITWLSNGHSVTEGVSETSFLSKSDHSFFKISYLTFLPSADEIYDCKVEHWGLDEPLLKHW**GGGGSG**
**GGGSGGGGS**EDFVYQFKGMCYFTNGTERVRLVSRSIYNREEIVRFDSDVGEFRAVTLLGLPAAEYWNSQK
DILERKRAAVDRVCRHNYQLELRTTLQRRVEPTVTISPSRTEALNHHNLLVCSVTDFYPAQIKVRWFRND
QEETAGVVSTPLIRNGDWTFQILVMLEMTPQRGDVYTCHVEHPSLQSPITVEWEPKSCDKTHTCPPCPAP
ELLGGPSVFLFPPKPKDTLMISRTPEVTCVVVDVSHEDPEVKFNWYVDGVEVHNAKTKPREEQYNSTYRV
VSVLTVLHQDWLNGKEYKCKVSNKGLPSSIEKTISKAKGQPREPQVYTLPPSRDELTKNQVSLTCLVKGF
YPSDIAVEWESNGQPENNYKTTPPVLDSDGSFFLYSKLTVDKSRWQQGNVFSCSVMHEALHNHYTQKSLS
LSPGK**ITIFITLFLLSVCYSATVTFFKVKWIFSSVVDLKQTIIPDYRNMIGQGA**

>DQA1*04:01-DQB1*02:01(DQ2)

**MILNKALLLGALALTTVMSPCGGEDIV**ADHVASYGVNLYQSYGPSGQYTHEFDGDEQFYVDLGRKETVWC
LPVLRQFRFDPQFALTNIAVTKHNLNILIKRSNSTAATNEVPEVTVFSKSPVTLGQPNTLICLVDNIFPP
VVNITWLSNGHSVTEGVSETSFLSKSDHSFFKISYLTFLPSADEIYDCKVEHWGLDEPLLKHW**GGGGSGG**
**GGSGGGGS**EDFVYQFKGMCYFTNGTERVRLVSRSIYNREEIVRFDSDVGEFRAVTLLGLPAAEYWNSQKD
ILERKRAAVDRVCRHNYQLELRTTLQRRVEPTVTISPSRTEALNHHNLLVCSVTDFYPAQIKVRWFRNDQ
EETAGVVSTPLIRNGDWTFQILVMLEMTPQRGDVYTCHVEHPSLQSPITVEWEPKSCDKTHTCPPCPAPE
LLGGPSVFLFPPKPKDTLMISRTPEVTCVVVDVSHEDPEVKFNWYVDGVEVHNAKTKPREEQYNSTYRVV
SVLTVLHQDWLNGKEYKCKVSNKGLPSSIEKTISKAKGQPREPQVYTLPPSRDELTKNQVSLTCLVKGFY
PSDIAVEWESNGQPENNYKTTPPVLDSDGSFFLYSKLTVDKSRWQQGNVFSCSVMHEALHNHYTQKSLSL
SPGK**ITIFITLFLLSVCYSATVTFFKVKWIFSSVVDLKQTIIPDYRNMIGQGA**

>DQA1*05:01-DQB1*02:01(DQ2)

**MILNKALMLGALALTTVMSPCGGEDIV**ADHVASYGVNLYQSYGPSGQYTHEFDGDEQFYVDLGRKETVWC
LPVLRQFRFDPQFALTNIAVLKHNLNSLIKRSNSTAATNEVPEVTVFSKSPVTLGQPNILICLVDNIFPP
VVNITWLSNGHSVTEGVSETSFLSKSDHSFFKISYLTLLPSAEESYDCKVEHWGLDKPLLKHW**GGGGSGG**
**GGSGGGGS**EDFVYQFKGMCYFTNGTERVRLVSRSIYNREEIVRFDSDVGEFRAVTLLGLPAAEYWNSQKD
ILERKRAAVDRVCRHNYQLELRTTLQRRVEPTVTISPSRTEALNHHNLLVCSVTDFYPAQIKVRWFRNDQ
EETAGVVSTPLIRNGDWTFQILVMLEMTPQRGDVYTCHVEHPSLQSPITVEWEPKSCDKTHTCPPCPAPE
LLGGPSVFLFPPKPKDTLMISRTPEVTCVVVDVSHEDPEVKFNWYVDGVEVHNAKTKPREEQYNSTYRVV
SVLTVLHQDWLNGKEYKCKVSNKGLPSSIEKTISKAKGQPREPQVYTLPPSRDELTKNQVSLTCLVKGFY
PSDIAVEWESNGQPENNYKTTPPVLDSDGSFFLYSKLTVDKSRWQQGNVFSCSVMHEALHNHYTQKSLSL
SPGK**ITIFITLFLLSVCYSATVTFFKVKWIFSSVVDLKQTIIPDYRNMIGQGA**

>DQA1*02:01-DQB1*02:02(DQ2)

**MILNKALMLGALALTTVMSPCGGEDIV**ADHVASYGVNLYQSYGPSGQFTHEFDGDEEFYVDLERKETVWK
LPLFHRLRFDPQFALTNIAVLKHNLNILIKRSNSTAATNEVPEVTVFSKSPVTLGQPNTLICLVDNIFPP
VVNITWLSNGHSVTEGVSETSFLSKSDHSFFKISYLTFLPSADEIYDCKVEHWGLDEPLLKHW**GGGGSGG**
**GGSGGGGS**EDFVYQFKGMCYFTNGTERVRLVSRSIYNREEIVRFDSDVGEFRAVTLLGLPAAEYWNSQKD
ILERKRAAVDRVCRHNYQLELRTTLQRRVEPTVTISPSRTEALNHHNLLVCSVTDFYPAQIKVRWFRNGQ
EETAGVVSTPLIRNGDWTFQILVMLEMTPQRGDVYTCHVEHPSLQSPITVEWEPKSCDKTHTCPPCPAPE
LLGGPSVFLFPPKPKDTLMISRTPEVTCVVVDVSHEDPEVKFNWYVDGVEVHNAKTKPREEQYNSTYRVV
SVLTVLHQDWLNGKEYKCKVSNKGLPSSIEKTISKAKGQPREPQVYTLPPSRDELTKNQVSLTCLVKGFY
PSDIAVEWESNGQPENNYKTTPPVLDSDGSFFLYSKLTVDKSRWQQGNVFSCSVMHEALHNHYTQKSLSL
SPGK**ITIFITLFLLSVCYSATVTFFKVKWIFSSVVDLKQTIIPDYRNMIGQGA**

>DQA1*02:01-DQB1*04:01(DQ4)

**MILNKALMLGALALTTVMSPCGGEDIV**ADHVASYGVNLYQSYGPSGQFTHEFDGDEEFYVDLERKETVWK
LPLFHRLRFDPQFALTNIAVLKHNLNILIKRSNSTAATNEVPEVTVFSKSPVTLGQPNTLICLVDNIFPP
VVNITWLSNGHSVTEGVSETSFLSKSDHSFFKISYLTFLPSADEIYDCKVEHWGLDEPLLKHW**GGGGSGG**
**GGSGGGGS**EDFVFQFKGMCYFTNGTELVRGVTRYIYNREEYARFDSDVGVYRAVTPLGRLDAEYWNSQKD
ILEEDRASVDTVCRHNYQLELRTTLQRRVEPTVTISPSRTEALNHHNLLVCSVTDFYPAQIKVRWFRNDQ
EETTGVVSTPLIRNGDWTFQILVMLEMTPQRGDVYTCHVEHPSLQNPIIVEW
EPKSCDKTHTCPPCPAPELLGGPSVFLFPPKPKDTLMISRTPEVTCVVVDVSHEDPEVKFNWYVDGVEVH
NAKTKPREEQYNSTYRVVSVLTVLHQDWLNGKEYKCKVSNKGLPSSIEKTISKAKGQPREPQVYTLPPSR
DELTKNQVSLTCLVKGFYPSDIAVEWESNGQPENNYKTTPPVLDSDGSFFLYSKLTVDKSRWQQGNVFSC
SVMHEALHNHYTQKSLSLSPGK**ITIFITLFLLSVCYSATVTFFKVKWIFSSVVDLKQTIIPDYRNMIGQG**
**A**

>DQA1*03:03-DQB1*04:01(DQ4)

**MILNKALMLGALALTTVMSPCGGEDIV**ADHVASYGVNLYQSYGPSGQYSHEFDGDEEFYVDLERKETVWQ
LPLFRRFRRFDPQFALTNIAVLKHNLNIVIKRSNSTAATNEVPEVTVFSKSPVTLGQPNTLICLVDNIFP
PVVNITWLSNGHSVTEGVSETSFLSKSDHSFFKISYLTFLPSDDEIYDCKVEHWGLDEPLLKHW**GGGGSG**
**GGGSGGGGS**EDFVFQFKGMCYFTNGTELVRGVTRYIYNREEYARFDSDVGVYRAVTPLGRLDAEYWNSQK
DILEEDRASVDTVCRHNYQLELRTTLQRRVEPTVTISPSRTEALNHHNLLVCSVTDFYPAQIKVRWFRND
QEETTGVVSTPLIRNGDWTFQILVMLEMTPQRGDVYTCHVEHPSLQNPIIVEWEPKSCDKTHTCPPCPAP
ELLGGPSVFLFPPKPKDTLMISRTPEVTCVVVDVSHEDPEVKFNWYVDGVEVHNAKTKPREEQYNSTYRV
VSVLTVLHQDWLNGKEYKCKVSNKGLPSSIEKTISKAKGQPREPQVYTLPPSRDELTKNQVSLTCLVKGF
YPSDIAVEWESNGQPENNYKTTPPVLDSDGSFFLYSKLTVDKSRWQQGNVFSCSVMHEALHNHYTQKSLS
LSPGK**ITIFITLFLLSVCYSATVTFFKVKWIFSSVVDLKQTIIPDYRNMIGQGA**

>DQA1*02:01-DQB1*04:02(DQ4)

**MILNKALMLGALALTTVMSPCGGEDIV**ADHVASYGVNLYQSYGPSGQFTHEFDGDEEFYVDLERKETVWK
LPLFHRLRFDPQFALTNIAVLKHNLNILIKRSNSTAATNEVPEVTVFSKSPVTLGQPNTLICLVDNIFPP
VVNITWLSNGHSVTEGVSETSFLSKSDHSFFKISYLTFLPSADEIYDCKVEHWGLDEPLLKHW**GGGGSGG**
**GGSGGGGS**EDFVFQFKGMCYFTNGTERVRGVTRYIYNREEYARFDSDVGVYRAVTPLGRLDAEYWNSQKD

ILEEDRASVDTVCRHNYQLELRTTLQRRVEPTVTISPSRTEALNHHNLLVCSVTDFYPAQIKVRWFRNDQ
EETTGVVSTPLIRNGDWTFQILVMLEMTPQRGDVYTCHVEHPSLQNPIIVEW
EPKSCDKTHTCPPCPAPELLGGPSVFLFPPKPKDTLMISRTPEVTCVVVDVSHEDPEVKFNWYVDGVEVH
NAKTKPREEQYNSTYRVVSVLTVLHQDWLNGKEYKCKVSNKGLPSSIEKTISKAKGQPREPQVYTLPPSR
DELTKNQVSLTCLVKGFYPSDIAVEWESNGQPENNYKTTPPVLDSDGSFFLYSKLTVDKSRWQQGNVFSC
SVMHEALHNHYTQKSLSLSPGK**ITIFITLFLLSVCYSATVTFFKVKWIFSSVVDLKQTIIPDYRNMIGQG
A**

>DQA1*04:01-DQB1*04:02(DQ4)

**MILNKALLLGALALTTVMSPCGGEDIV**ADHVASYGVNLYQSYGPSGQYTHEFDGDEQFYVDLGRKETVWC
LPVLRQFRFDPQFALTNIAVTKHNLNILIKRSNSTAATNEVPEVTVFSKSPVTLGQPNTLICLVDNIFPP
VVNITWLSNGHSVTEGVSETSFLSKSDHSFFKISYLTFLPSADEIYDCKVEHWGLDEPLLKHW**GGGGSGG
GGSGGGGS**EDFVFQFKGMCYFTNGTERVRGVTRYIYNREEYARFDSDVGVYRAVTPLGRLDAEYWNSQKD
ILEEDRASVDTVCRHNYQLELRTTLQRRVEPTVTISPSRTEALNHHNLLVCSVTDFYPAQIKVRWFRNDQ
EETTGVVSTPLIRNGDWTFQILVMLEMTPQRGDVYTCHVEHPSLQNPIIVEW
EPKSCDKTHTCPPCPAPELLGGPSVFLFPPKPKDTLMISRTPEVTCVVVDVSHEDPEVKFNWYVDGVEVH
NAKTKPREEQYNSTYRVVSVLTVLHQDWLNGKEYKCKVSNKGLPSSIEKTISKAKGQPREPQVYTLPPSR
DELTKNQVSLTCLVKGFYPSDIAVEWESNGQPENNYKTTPPVLDSDGSFFLYSKLTVDKSRWQQGNVFSC
SVMHEALHNHYTQKSLSLSPGK**ITIFITLFLLSVCYSATVTFFKVKWIFSSVVDLKQTIIPDYRNMIGQG
A**

>DQA1*01:01-DQB1*05:01(DQ5)

**MILNKALLLGALALTTVMSPCGGEDIV**ADHVASCGVNLYQFYGPSGQYTHEFDGDEEFYVDLERKETAWR
WPEFSKFGGFDPQGALRNMAVAKHNLNIMIKRYNSTAATNEVPEVTVFSKSPVTLGQPNTLICLVDNIFP
PVVNITWLSNGQSVTEGVSETSFLSKSDHSFFKISYLTFLPSADEIYDCKVEHWGLDQPLLKHW**GGGGSG
GGGSGGGGS**EDFVYQFKGLCYFTNGTERVRGVTRHIYNREEYVRFDSDVGVYRAVTPQGRPVAEYWNSQK
EVLEGARASVDRVCRHNYEVAYRGILQRRVEPTVTISPSRTEALNHHNLLICSVTDFYPSQIKVRWFRND
QEETAGVVSTPLIRNGDWTFQILVMLEMTPQRGDVYTCHVEHPSLQSPITVEWEPKSCDKTHTCPPCPAP
ELLGGPSVFLFPPKPKDTLMISRTPEVTCVVVDVSHEDPEVKFNWYVDGVEVHNAKTKPREEQYNSTYRV
VSVLTVLHQDWLNGKEYKCKVSNKGLPSSIEKTISKAKGQPREPQVYTLPPSRDELTKNQVSLTCLVKGF
YPSDIAVEWESNGQPENNYKTTPPVLDSDGSFFLYSKLTVDKSRWQQGNVFSCSVMHEALHNHYTQKSLS
LSPGK**ITIFITLFLLSVCYSATVTFFKVKWIFSSVVDLKQTIIPDYRNMIGQGA**

>DQA1*01:02-DQB1*05:01(DQ5)

**MILNKALLLGALALTTVMSPCGGEDIV**ADHVASCGVNLYQFYGPSGQYTHEFDGDEQFYVDLERKETAWR
WPEFSKFGGFDPQGALRNMAVAKHNLNIMIKRYNSTAATNEVPEVTVFSKSPVTLGQPNTLICLVDNIFP
PVVNITWLSNGQSVTEGVSETSFLSKSDHSFFKISYLTFLPSADEIYDCKVEHWGLDQPLLKHW**GGGGSG
GGGSGGGGS**EDFVYQFKGLCYFTNGTERVRGVTRHIYNREEYVRFDSDVGVYRAVTPQGRPVAEYWNSQK
EVLEGARASVDRVCRHNYEVAYRGILQRRVEPTVTISPSRTEALNHHNLLICSVTDFYPSQIKVRWFRND
QEETAGVVSTPLIRNGDWTFQILVMLEMTPQRGDVYTCHVEHPSLQSPITVEWEPKSCDKTHTCPPCPAP
ELLGGPSVFLFPPKPKDTLMISRTPEVTCVVVDVSHEDPEVKFNWYVDGVEVHNAKTKPREEQYNSTYRV
VSVLTVLHQDWLNGKEYKCKVSNKGLPSSIEKTISKAKGQPREPQVYTLPPSRDELTKNQVSLTCLVKGF

YPSDIAVEWESNGQPENNYKTTPPVLDSDGSFFLYSKLTVDKSRWQQGNVFSCSVMHEALHNHYTQKSLS
LSPGK**ITIFITLFLLSVCYSATVTFFKVKWIFSSVVDLKQTIIPDYRNMIGQGA**

>DQA1*01:03-DQB1*06:01(DQ6)

**MILNKALLLGALALTTVMSPCGGEDIV**ADHVASCGVNLYQFYGPSGQFTHEFDGDEQFYVDLEKKETAWR
WPEFSKFGGFDPQGALRNMAVAKHNLNIMIKRYNSTAATNEVPEVTVFSKSPVTLGQPNTLICLVDNIFP
PVVNITWLSNGHAVTEGVSETSFLSKSDHSFFKISYLTFLPSADEIYDCKVEHWGLDQPLLKHW**GGGGSG**
**GGGSGGGGS**EDFVLQFKAMCYFTNGTERVRYVTRYIYNREEDVRFDSDVGVYRAVTPQGRPDAEYWNSQK
DILERTRAELDTVCRHNYEVAFRGILQRRVEPTVTISPSRTEALNHHNLLVCSVTDFYPGQIKVRWFRND
QEETAGVVSTPLIRNGDWTFQILVMLEMTPQHGDVYTCHVEHPSLQSPITVEWEPKSCDKTHTCPPCPAP
ELLGGPSVFLFPPKPKDTLMISRTPEVTCVVVDVSHEDPEVKFNWYVDGVEVHNAKTKPREEQYNSTYRV
VSVLTVLHQDWLNGKEYKCKVSNKGLPSSIEKTISKAKGQPREPQVYTLPPSRDELTKNQVSLTCLVKGF
YPSDIAVEWESNGQPENNYKTTPPVLDSDGSFFLYSKLTVDKSRWQQGNVFSCSVMHEALHNHYTQKSLS
LSPGK**ITIFITLFLLSVCYSATVTFFKVKWIFSSVVDLKQTIIPDYRNMIGQGA**

>DQA1*01:01-DQB1*06:02(DQ6)

**MILNKALLLGALALTTVMSPCGGEDIV**ADHVASCGVNLYQFYGPSGQYTHEFDGDEEFYVDLERKETAWR
WPEFSKFGGFDPQGALRNMAVAKHNLNIMIKRYNSTAATNEVPEVTVFSKSPVTLGQPNTLICLVDNIFP
PVVNITWLSNGQSVTEGVSETSFLSKSDHSFFKISYLTFLPSADEIYDCKVEHWGLDQPLLKHW**GGGGSG**
**GGGSGGGGS**EDFVFQFKGMCYFTNGTERVRLVTRYIYNREEYARFDSDVGVYRAVTPQGRPDAEYWNSQK
EVLEGTRAELDTVCRHNYEVAFRGILQRRVEPTVTISPSRTEALNHHNLLVCSVTDFYPGQIKVRWFRND
QEETAGVVSTPLIRNGDWTFQILVMLEMTPQRGDVYTCHVEHPSLQSPITVEWEPKSCDKTHTCPPCPAP
ELLGGPSVFLFPPKPKDTLMISRTPEVTCVVVDVSHEDPEVKFNWYVDGVEVHNAKTKPREEQYNSTYRV
VSVLTVLHQDWLNGKEYKCKVSNKGLPSSIEKTISKAKGQPREPQVYTLPPSRDELTKNQVSLTCLVKGF
YPSDIAVEWESNGQPENNYKTTPPVLDSDGSFFLYSKLTVDKSRWQQGNVFSCSVMHEALHNHYTQKSLS
LSPGK**ITIFITLFLLSVCYSATVTFFKVKWIFSSVVDLKQTIIPDYRNMIGQGA**

>DQA1*01:02-DQB1*06:02(DQ6)

**MILNKALLLGALALTTVMSPCGGEDIV**ADHVASCGVNLYQFYGPSGQYTHEFDGDEQFYVDLERKETAWR
WPEFSKFGGFDPQGALRNMAVAKHNLNIMIKRYNSTAATNEVPEVTVFSKSPVTLGQPNTLICLVDNIFP
PVVNITWLSNGQSVTEGVSETSFLSKSDHSFFKISYLTFLPSADEIYDCKVEHWGLDQPLLKHW**GGGGSG**
**GGGSGGGGS**EDFVFQFKGMCYFTNGTERVRLVTRYIYNREEYARFDSDVGVYRAVTPQGRPDAEYWNSQK
EVLEGTRAELDTVCRHNYEVAFRGILQRRVEPTVTISPSRTEALNHHNLLVCSVTDFYPGQIKVRWFRND
QEETAGVVSTPLIRNGDWTFQILVMLEMTPQRGDVYTCHVEHPSLQSPITVEWEPKSCDKTHTCPPCPAP
ELLGGPSVFLFPPKPKDTLMISRTPEVTCVVVDVSHEDPEVKFNWYVDGVEVHNAKTKPREEQYNSTYRV
VSVLTVLHQDWLNGKEYKCKVSNKGLPSSIEKTISKAKGQPREPQVYTLPPSRDELTKNQVSLTCLVKGF
YPSDIAVEWESNGQPENNYKTTPPVLDSDGSFFLYSKLTVDKSRWQQGNVFSCSVMHEALHNHYTQKSLS
LSPGK**ITIFITLFLLSVCYSATVTFFKVKWIFSSVVDLKQTIIPDYRNMIGQGA**

>DQA1*01:03-DQB1*06:03(DQ6)

**MILNKALLLGALALTTVMSPCGGEDIV**ADHVASCGVNLYQFYGPSGQFTHEFDGDEQFYVDLEKKETAWR
WPEFSKFGGFDPQGALRNMAVAKHNLNIMIKRYNSTAATNEVPEVTVFSKSPVTLGQPNTLICLVDNIFP

PVVNITWLSNGHAVTEGVSETSFLSKSDHSFFKISYLTFLPSADEIYDCKVEHWGLDQPLLKHW**GGGGSG GGGSGGGGS**EDFVYQFKGMCYFTNGTERVRLVTRHIYNREEYARFDSDVGVYRAVTPQGRPDAEYWNSQK EVLEGTRAELDTVCRHNYEVAFRGILQRRVEPTVTISPSRTEALNHHNLLVCSVTDFYPGQIKVRWFRND QEETAGVVSTPLIRNGDWTFQILVMLEMTPQRGDVYTCHVEHPSLQSPITVEWEPKSCDKTHTCPPCPAP ELLGGPSVFLFPPKPKDTLMISRTPEVTCVVVDVSHEDPEVKFNWYVDGVEVHNAKTKPREEQYNSTYRV VSVLTVLHQDWLNGKEYKCKVSNKGLPSSIEKTISKAKGQPREPQVYTLPPSRDELTKNQVSLTCLVKGF YPSDIAVEWESNGQPENNYKTTPPVLDSDGSFFLYSKLTVDKSRWQQGNVFSCSVMHEALHNHYTQKSLS LSPGK**ITIFITLFLLSVCYSATVTFFKVKWIFSSVVDLKQTIIPDYRNMIGQGA**

>DQA1*01:02-DQB1*06:04(DQ6)

**MILNKALLLGALALTTVMSPCGGEDIV**ADHVASCGVNLYQFYGPSGQYTHEFDGDEQFYVDLERKETAWR WPEFSKFGGFDPQGALRNMAVAKHNLNIMIKRYNSTAATNEVPEVTVFSKSPVTLGQPNTLICLVDNIFP PVVNITWLSNGQSVTEGVSETSFLSKSDHSFFKISYLTFLPSADEIYDCKVEHWGLDQPLLKHW**GGGGSG GGGSGGGGS**EDFVYQFKGMCYFTNGTERVRLVTRHIYNREEYARFDSDVGVYRAVTPQGRPVAEYWNSQK EVLERTRAELDTVCRHNYEVGYRGILQRRVEPTVTISPSRTEALNHHNLLVCSVTDFYPGQIKVQWFRND QEETAGVVSTPLIRNGDWTFQILVMLEMTPQRGDVYTCHVEHPSLQSPITVEWEPKSCDKTHTCPPCPAP ELLGGPSVFLFPPKPKDTLMISRTPEVTCVVVDVSHEDPEVKFNWYVDGVEVHNAKTKPREEQYNSTYRV VSVLTVLHQDWLNGKEYKCKVSNKGLPSSIEKTISKAKGQPREPQVYTLPPSRDELTKNQVSLTCLVKGF YPSDIAVEWESNGQPENNYKTTPPVLDSDGSFFLYSKLTVDKSRWQQGNVFSCSVMHEALHNHYTQKSLS LSPGK**ITIFITLFLLSVCYSATVTFFKVKWIFSSVVDLKQTIIPDYRNMIGQGA**

>DQA1*01:02-DQB1*06:09(DQ6)

**MILNKALLLGALALTTVMSPCGGEDIV**ADHVASCGVNLYQFYGPSGQYTHEFDGDEQFYVDLERKETAWR WPEFSKFGGFDPQGALRNMAVAKHNLNIMIKRYNSTAATNEVPEVTVFSKSPVTLGQPNTLICLVDNIFP PVVNITWLSNGQSVTEGVSETSFLSKSDHSFFKISYLTFLPSADEIYDCKVEHWGLDQPLLKHW**GGGGSG GGGSGGGGS**EDFVYQFKGMCYFTNGTERVRLVTRYIYNREEYARFDSDVGVYRAVTPQGRPVAEYWNSQK EVLERTRAELDTVCRHNYEVGYRGILQRRVEPTVTISPSRTEALNHHNLLVCSVTDFYPGQIKVQWFRND QEETAGVVSTPLIRNGDWTFQILVMLEMTPQRGDVYTCHVEHPSLQSPITVEWEPKSCDKTHTCPPCPAP ELLGGPSVFLFPPKPKDTLMISRTPEVTCVVVDVSHEDPEVKFNWYVDGVEVHNAKTKPREEQYNSTYRV VSVLTVLHQDWLNGKEYKCKVSNKGLPSSIEKTISKAKGQPREPQVYTLPPSRDELTKNQVSLTCLVKGF YPSDIAVEWESNGQPENNYKTTPPVLDSDGSFFLYSKLTVDKSRWQQGNVFSCSVMHEALHNHYTQKSLS LSPGK**ITIFITLFLLSVCYSATVTFFKVKWIFSSVVDLKQTIIPDYRNMIGQGA**

>DQA1*02:01-DQB1*03:01(DQ7)

**MILNKALMLGALALTTVMSPCGGEDIV**ADHVASYGVNLYQSYGPSGQFTHEFDGDEEFYVDLERKETVWK LPLFHRLRFDPQFALTNIAVLKHNLNILIKRSNSTAATNEVPEVTVFSKSPVTLGQPNTLICLVDNIFPP VVNITWLSNGHSVTEGVSETSFLSKSDHSFFKISYLTFLPSADEIYDCKVEHWGLDEPLLKHW**GGGGSGG GGSGGGGS**EDFVYQFKAMCYFTNGTERVRYVTRYIYNREEYARFDSDVEVYRAVTPLGPPDAEYWNSQKE VLERTRAELDTVCRHNYQLELRTTLQRRVEPTVTISPSRTEALNHHNLLVCSVTDFYPAQIKVRWFRNDQ EETTGVVSTPLIRNGDWTFQILVMLEMTPQHGDVYTCHVEHPSLQNPITVEW EPKSCDKTHTCPPCPAPELLGGPSVFLFPPKPKDTLMISRTPEVTCVVVDVSHEDPEVKFNWYVDGVEVH NAKTKPREEQYNSTYRVVSVLTVLHQDWLNGKEYKCKVSNKGLPSSIEKTISKAKGQPREPQVYTLPPSR DELTKNQVSLTCLVKGFYPSDIAVEWESNGQPENNYKTTPPVLDSDGSFFLYSKLTVDKSRWQQGNVFSC SVMHEALHNHYTQKSLSLSPGK**ITIFITLFLLSVCYSATVTFFKVKWIFSSVVDLKQTIIPDYRNMIGQG A**

>DQA1*03:01-DQB1*03:01(DQ7)

**MILNKALMLGALALTTVMSPCGGEDIV**ADHVASYGVNLYQSYGPSGQYSHEFDGDEEFYVDLERKETVWQ
LPLFRRFRRFDPQFALTNIAVLKHNLNIVIKRSNSTAATNEVPEVTVFSKSPVTLGQPNTLICLVDNIFP
PVVNITWLSNGHSVTEGVSETSFLSKSDHSFFKISYLTFLPSADEIYDCKVEHWGLDEPLLKHW**GGGGSG**
**GGGSGGGGS**EDFVYQFKAMCYFTNGTERVRYVTRYIYNREEYARFDSDVEVYRAVTPLGPPDAEYWNSQK
EVLERTRAELDTVCRHNYQLELRTTLQRRVEPTVTISPSRTEALNHHNLLVCSVTDFYPAQIKVRWFRND
QEETTGVVSTPLIRNGDWTFQILVMLEMTPQHGDVYTCHVEHPSLQNPITVEWEPKSCDKTHTCPPCPAP
ELLGGPSVFLFPPKPKDTLMISRTPEVTCVVVDVSHEDPEVKFNWYVDGVEVHNAKTKPREEQYNSTYRV
VSVLTVLHQDWLNGKEYKCKVSNKGLPSSIEKTISKAKGQPREPQVYTLPPSRDELTKNQVSLTCLVKGF
YPSDIAVEWESNGQPENNYKTTPPVLDSDGSFFLYSKLTVDKSRWQQGNVFSCSVMHEALHNHYTQKSLS
LSPGK**ITIFITLFLLSVCYSATVTFFKVKWIFSSVVDLKQTIIPDYRNMIGQGA**

>DQA1*05:03-DQB1*03:01(DQ7)

**MILNKALMLGALALTTVMSPCGGEDIV**ADHVASYGVNLYQSYGPSGQYTHEFDGDEQFYVDLGRKETVWC
LPVLRQFRFDPQFALTNIAVLKHNLNSLIKRSNSTAATNEVPEVTVFSKSPVTLGQPNILICLVDNIFPP
VVNITWLSNGHSVTEGVSETSFLSKSDHSFFKISYLTLLPSSEESYDCKVEHWGLDKPLLKHW**GGGGSGG**
**GGSGGGGS**EDFVYQFKAMCYFTNGTERVRYVTRYIYNREEYARFDSDVEVYRAVTPLGPPDAEYWNSQKE
VLERTRAELDTVCRHNYQLELRTTLQRRVEPTVTISPSRTEALNHHNLLVCSVTDFYPAQIKVRWFRNDQ
EETTGVVSTPLIRNGDWTFQILVMLEMTPQHGDVYTCHVEHPSLQNPITVEWEPKSCDKTHTCPPCPAPE
LLGGPSVFLFPPKPKDTLMISRTPEVTCVVVDVSHEDPEVKFNWYVDGVEVHNAKTKPREEQYNSTYRVV
SVLTVLHQDWLNGKEYKCKVSNKGLPSSIEKTISKAKGQPREPQVYTLPPSRDELTKNQVSLTCLVKGFY
PSDIAVEWESNGQPENNYKTTPPVLDSDGSFFLYSKLTVDKSRWQQGNVFSCSVMHEALHNHYTQKSLSL
SPGK**ITIFITLFLLSVCYSATVTFFKVKWIFSSVVDLKQTIIPDYRNMIGQGA**

>DQA1*06:01-DQB1*03:01(DQ7)

**MILNKALLLGALALTTVMSPCGGEDIV**ADHVASYGVNLYQSYGPSGQFTHEFDGDEQFYVDLGRKETVWC
LPVLRQFRFDPQFALTNIAVTKHNLNILIKRSNSTAATNEVPEVTVFSKSPVTLGQPNTLICLVDNIFPP
VVNITWLSNGHSVTEGVSETSFLSKSDHSFFKISYLTFLPSADEIYDCKVEHWGLDEPLLKHW**GGGGSGG**
**GGSGGGGS**EDFVYQFKAMCYFTNGTERVRYVTRYIYNREEYARFDSDVEVYRAVTPLGPPDAEYWNSQKE
VLERTRAELDTVCRHNYQLELRTTLQRRVEPTVTISPSRTEALNHHNLLVCSVTDFYPAQIKVRWFRNDQ
EETTGVVSTPLIRNGDWTFQILVMLEMTPQHGDVYTCHVEHPSLQNPITVEWEPKSCDKTHTCPPCPAPE
LLGGPSVFLFPPKPKDTLMISRTPEVTCVVVDVSHEDPEVKFNWYVDGVEVHNAKTKPREEQYNSTYRVV
SVLTVLHQDWLNGKEYKCKVSNKGLPSSIEKTISKAKGQPREPQVYTLPPSRDELTKNQVSLTCLVKGFY
PSDIAVEWESNGQPENNYKTTPPVLDSDGSFFLYSKLTVDKSRWQQGNVFSCSVMHEALHNHYTQKSLSL
SPGK**ITIFITLFLLSVCYSATVTFFKVKWIFSSVVDLKQTIIPDYRNMIGQGA**

>DQA1*05:05-DQB1*03:19(DQ7)

**MILNKALMLGTLALTTVMSPCGGEDIV**ADHVASYGVNLYQSYGPSGQYTHEFDGDEQFYVDLGRKETVWC
LPVLRQFRFDPQFALTNIAVLKHNLNSLIKRSNSTAATNEVPEVTVFSKSPVTLGQPNILICLVDNIFPP
VVNITWLSNGHSVTEGVSETSFLSKSDHSFFKISYLTLLPSAEESYDCKVEHWGLDKPLLKHW**GGGGSGG**
**GGSGGGGS**EDFVYQFKAMCYFTNGTERVRYVTRYIYNREEYARFDSDVEVYRAVTPLGPPDAEYWNSQKE
VLERTRAELDTVCRHNYQLELRTTLQRRVEPTVTISPSRTEALNHHNLLVCSVTDFYPAQIKVRWFRNDQ

EETTGVVSTPLIRNGDWTFQILVMLEMTPQHGDVYTCHVEHPSLQNPIIVEWEPKSCDKTHTCPPCPAPE
LLGGPSVFLFPPKPKDTLMISRTPEVTCVVVDVSHEDPEVKFNWYVDGVEVHNAKTKPREEQYNSTYRVV
SVLTVLHQDWLNGKEYKCKVSNKGLPSSIEKTISKAKGQPREPQVYTLPPSRDELTKNQVSLTCLVKGFY
PSDIAVEWESNGQPENNYKTTPPVLDSDGSFFLYSKLTVDKSRWQQGNVFSCSVMHEALHNHYTQKSLSL
SPGK**ITIFITLFLLSVCYSATVTFFKVKWIFSSVVDLKQTIIPDYRNMIGQGA**

>DQA1*02:01-DQB1*03:02(DQ8)

**MILNKALMLGALALTTVMSPCGGEDIV**ADHVASYGVNLYQSYGPSGQFTHEFDGDEEFYVDLERKETVWK
LPLFHRLRFDPQFALTNIAVLKHNLNILIKRSNSTAATNEVPEVTVFSKSPVTLGQPNTLICLVDNIFPP
VVNITWLSNGHSVTEGVSETSFLSKSDHSFFKISYLTFLPSADEIYDCKVEHWGLDEPLLKHW**GGGGSGG
GGSGGGGS**EDFVYQFKGMCYFTNGTERVRLVTRYIYNREEYARFDSDVGVYRAVTPLGPPAAEYWNSQKE
VLERTRAELDTVCRHNYQLELRTTLQRRVEPTVTISPSRTEALNHHNLLVCSVTDFYPAQIKVRWFRNDQ
EETTGVVSTPLIRNGDWTFQILVMLEMTPQRGDVYTCHVEHPSLQNPIIVEWEPKSCDKTHTCPPCPAPE
LLGGPSVFLFPPKPKDTLMISRTPEVTCVVVDVSHEDPEVKFNWYVDGVEVHNAKTKPREEQYNSTYRVV
SVLTVLHQDWLNGKEYKCKVSNKGLPSSIEKTISKAKGQPREPQVYTLPPSRDELTKNQVSLTCLVKGFY
PSDIAVEWESNGQPENNYKTTPPVLDSDGSFFLYSKLTVDKSRWQQGNVFSCSVMHEALHNHYTQKSLSL
SPGK**ITIFITLFLLSVCYSATVTFFKVKWIFSSVVDLKQTIIPDYRNMIGQGA**

>DQA1*03:01-DQB1*03:02(DQ8)

**MILNKALMLGALALTTVMSPCGGEDIV**ADHVASYGVNLYQSYGPSGQYSHEFDGDEEFYVDLERKETVWQ
LPLFRRFRRFDPQFALTNIAVLKHNLNIVIKRSNSTAATNEVPEVTVFSKSPVTLGQPNTLICLVDNIFP
PVVNITWLSNGHSVTEGVSETSFLSKSDHSFFKISYLTFLPSADEIYDCKVEHWGLDEPLLKHW**GGGGSG
GGGSGGGGS**EDFVYQFKGMCYFTNGTERVRLVTRYIYNREEYARFDSDVGVYRAVTPLGPPAAEYWNSQK
EVLERTRAELDTVCRHNYQLELRTTLQRRVEPTVTISPSRTEALNHHNLLVCSVTDFYPAQIKVRWFRND
QEETTGVVSTPLIRNGDWTFQILVMLEMTPQRGDVYTCHVEHPSLQNPIIVEWEPKSCDKTHTCPPCPAP
ELLGGPSVFLFPPKPKDTLMISRTPEVTCVVVDVSHEDPEVKFNWYVDGVEVHNAKTKPREEQYNSTYRV
VSVLTVLHQDWLNGKEYKCKVSNKGLPSSIEKTISKAKGQPREPQVYTLPPSRDELTKNQVSLTCLVKGF
YPSDIAVEWESNGQPENNYKTTPPVLDSDGSFFLYSKLTVDKSRWQQGNVFSCSVMHEALHNHYTQKSLS
LSPGK**ITIFITLFLLSVCYSATVTFFKVKWIFSSVVDLKQTIIPDYRNMIGQGA**

>DQA1*03:03-DQB1*03:02(DQ8)

**MILNKALMLGALALTTVMSPCGGEDIV**ADHVASYGVNLYQSYGPSGQYSHEFDGDEEFYVDLERKETVWQ
LPLFRRFRRFDPQFALTNIAVLKHNLNIVIKRSNSTAATNEVPEVTVFSKSPVTLGQPNTLICLVDNIFP
PVVNITWLSNGHSVTEGVSETSFLSKSDHSFFKISYLTFLPSDDEIYDCKVEHWGLDEPLLKHW**GGGGSG
GGGSGGGGS**EDFVYQFKGMCYFTNGTERVRLVTRYIYNREEYARFDSDVGVYRAVTPLGPPAAEYWNSQK
EVLERTRAELDTVCRHNYQLELRTTLQRRVEPTVTISPSRTEALNHHNLLVCSVTDFYPAQIKVRWFRND
QEETTGVVSTPLIRNGDWTFQILVMLEMTPQRGDVYTCHVEHPSLQNPIIVEWEPKSCDKTHTCPPCPAP
ELLGGPSVFLFPPKPKDTLMISRTPEVTCVVVDVSHEDPEVKFNWYVDGVEVHNAKTKPREEQYNSTYRV
VSVLTVLHQDWLNGKEYKCKVSNKGLPSSIEKTISKAKGQPREPQVYTLPPSRDELTKNQVSLTCLVKGF
YPSDIAVEWESNGQPENNYKTTPPVLDSDGSFFLYSKLTVDKSRWQQGNVFSCSVMHEALHNHYTQKSLS
LSPGK**ITIFITLFLLSVCYSATVTFFKVKWIFSSVVDLKQTIIPDYRNMIGQGA**

>DQA1*02:01-DQB1*03:03(DQ9)

**MILNKALMLGALALTTVMSPCGGEDIV**ADHVASYGVNLYQSYGPSGQFTHEFDGDEEFYVDLERKETVWK
LPLFHRLRFDPQFALTNIAVLKHNLNILIKRSNSTAATNEVPEVTVFSKSPVTLGQPNTLICLVDNIFPP
VVNITWLSNGHSVTEGVSETSFLSKSDHSFFKISYLTFLPSADEIYDCKVEHWGLDEPLLKHW**GGGGSGG**
**GGSGGGGS**EDFVYQFKGMCYFTNGTERVRLVTRYIYNREEYARFDSDVGVYRAVTPLGPPDAEYWNSQKE
VLERTRAELDTVCRHNYQLELRTTLQRRVEPTVTISPSRTEALNHHNLLVCSVTDFYPAQIKVRWFRNDQ
EETTGVVSTPLIRNGDWTFQILVMLEMTPQRGDVYTCHVEHPSLQNPIIVEWEPKSCDKTHTCPPCPAPE
LLGGPSVFLFPPKPKDTLMISRTPEVTCVVVDVSHEDPEVKFNWYVDGVEVHNAKTKPREEQYNSTYRVV
SVLTVLHQDWLNGKEYKCKVSNKGLPSSIEKTISKAKGQPREPQVYTLPPSRDELTKNQVSLTCLVKGFY
PSDIAVEWESNGQPENNYKTTPPVLDSDGSFFLYSKLTVDKSRWQQGNVFSCSVMHEALHNHYTQKSLSL
SPGK**ITIFITLFLLSVCYSATVTFFKVKWIFSSVVDLKQTIIPDYRNMIGQGA**

>DQA1*03:01-DQB1*03:03(DQ9)

**MILNKALMLGALALTTVMSPCGGEDIV**ADHVASYGVNLYQSYGPSGQYSHEFDGDEEFYVDLERKETVWQ
LPLFRRFRRFDPQFALTNIAVLKHNLNIVIKRSNSTAATNEVPEVTVFSKSPVTLGQPNTLICLVDNIFP
PVVNITWLSNGHSVTEGVSETSFLSKSDHSFFKISYLTFLPSADEIYDCKVEHWGLDEPLLKHW**GGGGSG**
**GGGSGGGGS**EDFVYQFKGMCYFTNGTERVRLVTRYIYNREEYARFDSDVGVYRAVTPLGPPDAEYWNSQK
EVLERTRAELDTVCRHNYQLELRTTLQRRVEPTVTISPSRTEALNHHNLLVCSVTDFYPAQIKVRWFRND
QEETTGVVSTPLIRNGDWTFQILVMLEMTPQRGDVYTCHVEHPSLQNPIIVEWEPKSCDKTHTCPPCPAP
ELLGGPSVFLFPPKPKDTLMISRTPEVTCVVVDVSHEDPEVKFNWYVDGVEVHNAKTKPREEQYNSTYRV
VSVLTVLHQDWLNGKEYKCKVSNKGLPSSIEKTISKAKGQPREPQVYTLPPSRDELTKNQVSLTCLVKGF
YPSDIAVEWESNGQPENNYKTTPPVLDSDGSFFLYSKLTVDKSRWQQGNVFSCSVMHEALHNHYTQKSLS
LSPGK**ITIFITLFLLSVCYSATVTFFKVKWIFSSVVDLKQTIIPDYRNMIGQGA**

>DQA1*03:02-DQB1*03:03(DQ9)

**MILNKALMLGALALTTVTSPCGGEDIV**ADHVASYGVNLYQSYGPSGQYSHEFDGDEEFYVDLERKETVWQ
LPLFRRFRRFDPQFALTNIAVLKHNLNIVIKRSNSTAATNEVPEVTVFSKSPVTLGQPNTLICLVDNIFP
PVVNITWLSNGHSVTEGVSETSFLSKSDHSFFKISYLTFLPSDDEIYDCKVEHWGLDEPLLKHW**GGGGSG**
**GGGSGGGGS**EDFVYQFKGMCYFTNGTERVRLVTRYIYNREEYARFDSDVGVYRAVTPLGPPDAEYWNSQK
EVLERTRAELDTVCRHNYQLELRTTLQRRVEPTVTISPSRTEALNHHNLLVCSVTDFYPAQIKVRWFRND
QEETTGVVSTPLIRNGDWTFQILVMLEMTPQRGDVYTCHVEHPSLQNPIIVEWEPKSCDKTHTCPPCPAP
ELLGGPSVFLFPPKPKDTLMISRTPEVTCVVVDVSHEDPEVKFNWYVDGVEVHNAKTKPREEQYNSTYRV
VSVLTVLHQDWLNGKEYKCKVSNKGLPSSIEKTISKAKGQPREPQVYTLPPSRDELTKNQVSLTCLVKGF
YPSDIAVEWESNGQPENNYKTTPPVLDSDGSFFLYSKLTVDKSRWQQGNVFSCSVMHEALHNHYTQKSLS
LSPGK**ITIFITLFLLSVCYSATVTFFKVKWIFSSVVDLKQTIIPDYRNMIGQGA**

HLA DP

>DPA1*01:03-DPB1*01:01(DP1)

**MRPEDRMFHIRAVILRALSLAFLLSLRGAGAIK**ADHVSTYAAFVQTHRPTGEFMFEFDEDEMFYVDLDKK
ETVWHLEEFGQAFSFEAQGGLANIAILNNNLNTLIQRSNHTQATNDPPEVTVFPKEPVELGQPNTLICHI

DKFFPPVLNVTWLCNGELVTEGVAESLFLPRTDYSFHKFHYLTFVPSAEDFYDCRVEHWGLDQPLLKHW**G**
**GGGSGGGGSGGGGS**ENYVYQGRQECYAFNGTQRFLERYIYNREEYARFDSDVGEFRAVTELGRPAAEYWN
SQKDILEEKRAVPDRVCRHNYELDEAVTLQRRVQPKVNVSPSKKGPLQHHNLLVCHVTDFYPGSIQVRWF
LNGQEETAGVVSTNLIRNGDWTFQILVMLEMTPQQGDVYICQVEHTSLDSPVTVEWEPKSCDKTHTCPPC
PAPELLGGPSVFLFPPKPKDTLMISRTPEVTCVVVDVSHEDPEVKFNWYVDGVEVHNAKTKPREEQYNST
YRVVSVLTVLHQDWLNGKEYKCKVSNKGLPSSIEKTISKAKGQPREPQVYTLPPSRDELTKNQVSLTCLV
KGFYPSDIAVEWESNGQPENNYKTTPPVLDSDGSFFLYSKLTVDKSRWQQGNVFSCSVMHEALHNHYTQK
SLSLSPGK**ITIFITLFLLSVCYSATVTFFKVKWIFSSVVDLKQTIIPDYRNMIGQGA**

>DPA1*02:01-DPB1*01:01(DP1)

**MRPEDRMFHIRAVILRALSLAFLLSLRGAGAIK**ADHVSTYAAFVQTHRPTGEFMFEFDEDEQFYVDLDKK
ETVWHLEEFGRAFSFEAQGGLANIAILNNNLNTLIQRSNHTQAANDPPEVTVFPKEPVELGQPNTLICHI
DRFFPPVLNVTWLCNGEPVTEGVAESLFLPRTDYSFHKFHYLTFVPSAEDVYDCRVEHWGLDQPLLKHW**G**
**GGGSGGGGSGGGGS**ENYVYQGRQECYAFNGTQRFLERYIYNREEYARFDSDVGEFRAVTELGRPAAEYWN
SQKDILEEKRAVPDRVCRHNYELDEAVTLQRRVQPKVNVSPSKKGPLQHHNLLVCHVTDFYPGSIQVRWF
LNGQEETAGVVSTNLIRNGDWTFQILVMLEMTPQQGDVYICQVEHTSLDSPVTVEWEPKSCDKTHTCPPC
PAPELLGGPSVFLFPPKPKDTLMISRTPEVTCVVVDVSHEDPEVKFNWYVDGVEVHNAKTKPREEQYNST
YRVVSVLTVLHQDWLNGKEYKCKVSNKGLPSSIEKTISKAKGQPREPQVYTLPPSRDELTKNQVSLTCLV
KGFYPSDIAVEWESNGQPENNYKTTPPVLDSDGSFFLYSKLTVDKSRWQQGNVFSCSVMHEALHNHYTQK
SLSLSPGK**ITIFITLFLLSVCYSATVTFFKVKWIFSSVVDLKQTIIPDYRNMIGQGA**

>DPA1*01:03-DPB1*02:01(DP2)

**MRPEDRMFHIRAVILRALSLAFLLSLRGAGAIK**ADHVSTYAAFVQTHRPTGEFMFEFDEDEMFYVDLDKK
ETVWHLEEFGQAFSFEAQGGLANIAILNNNLNTLIQRSNHTQATNDPPEVTVFPKEPVELGQPNTLICHI
DKFFPPVLNVTWLCNGELVTEGVAESLFLPRTDYSFHKFHYLTFVPSAEDFYDCRVEHWGLDQPLLKHW**G**
**GGGSGGGGSGGGGS**ENYLFQGRQECYAFNGTQRFLERYIYNREEFVRFDSDVGEFRAVTELGRPDEEYWN
SQKDILEEERAVPDRMCRHNYELGGPMTLQRRVQPRVNVSPSKKGPLQHHNLLVCHVTDFYPGSIQVRWF
LNGQEETAGVVSTNLIRNGDWTFQILVMLEMTPQQGDVYTCQVEHTSLDSPVTVEWEPKSCDKTHTCPPC
PAPELLGGPSVFLFPPKPKDTLMISRTPEVTCVVVDVSHEDPEVKFNWYVDGVEVHNAKTKPREEQYNST
YRVVSVLTVLHQDWLNGKEYKCKVSNKGLPSSIEKTISKAKGQPREPQVYTLPPSRDELTKNQVSLTCLV
KGFYPSDIAVEWESNGQPENNYKTTPPVLDSDGSFFLYSKLTVDKSRWQQGNVFSCSVMHEALHNHYTQK
SLSLSPGK**ITIFITLFLLSVCYSATVTFFKVKWIFSSVVDLKQTIIPDYRNMIGQGA**

>DPA1*01:03-DPB1*03:01(DP3)

**MRPEDRMFHIRAVILRALSLAFLLSLRGAGAIK**ADHVSTYAAFVQTHRPTGEFMFEFDEDEMFYVDLDKK
ETVWHLEEFGQAFSFEAQGGLANIAILNNNLNTLIQRSNHTQATNDPPEVTVFPKEPVELGQPNTLICHI
DKFFPPVLNVTWLCNGELVTEGVAESLFLPRTDYSFHKFHYLTFVPSAEDFYDCRVEHWGLDQPLLKHW**G**
**GGGSGGGGSGGGGS**ENYVYQLRQECYAFNGTQRFLERYIYNREEFVRFDSDVGEFRAVTELGRPDEDYWN
SQKDLLEEKRAVPDRVCRHNYELDEAVTLQRRVQPKVNVSPSKKGPLQHHNLLVCHVTDFYPGSIQVRWF
LNGQEETAGVVSTNLIRNGDWTFQILVMLEMTPQQGDVYICQVEHTSLDSPVTVEWEPKSCDKTHTCPPC
PAPELLGGPSVFLFPPKPKDTLMISRTPEVTCVVVDVSHEDPEVKFNWYVDGVEVHNAKTKPREEQYNST
YRVVSVLTVLHQDWLNGKEYKCKVSNKGLPSSIEKTISKAKGQPREPQVYTLPPSRDELTKNQVSLTCLV

KGFYPSDIAVEWESNGQPENNYKTTPPVLDSDGSFFLYSKLTVDKSRWQQGNVFSCSVMHEALHNHYTQK
SLSLSPGK**ITIFITLFLLSVCYSATVTFFKVKWIFSSVVDLKQTIIPDYRNMIGQGA**

>DPA1*01:05-DPB1*03:01(DP3)

**MRPEDRMFHIRAVILRALSLAFLLSLRGAGAIK**ADHVSTYAAFVQTHRPTGEFMFEFDEDEMFYVDLDKK
ETVWHLEEFGQAFSFEAQGGLANIAILNNNLNTLIQRSNHTQAANDPPEVTVFPKEPVELGQPNTLICHI
DKFFPPVLNVTWLCNGELVTEGVAESLFLPRTDYSFHKFHYLTFVPSAEDFYDCRVEHWGLDQPLLKHW**G**
**GGGSGGGGSGGGGS**ENYVYQLRQECYAFNGTQRFLERYIYNREEFVRFDSDVGEFRAVTELGRPDEDYWN
SQKDLLEEKRAVPDRVCRHNYELDEAVTLQRRVQPKVNVSPSKKGPLQHHNLLVCHVTDFYPGSIQVRWF
LNGQEETAGVVSTNLIRNGDWTFQILVMLEMTPQQGDVYICQVEHTSLDSPVTVEWEPKSCDKTHTCPPC
PAPELLGGPSVFLFPPKPKDTLMISRTPEVTCVVVDVSHEDPEVKFNWYVDGVEVHNAKTKPREEQYNST
YRVVSVLTVLHQDWLNGKEYKCKVSNKGLPSSIEKTISKAKGQPREPQVYTLPPSRDELTKNQVSLTCLV
KGFYPSDIAVEWESNGQPENNYKTTPPVLDSDGSFFLYSKLTVDKSRWQQGNVFSCSVMHEALHNHYTQK
SLSLSPGK**ITIFITLFLLSVCYSATVTFFKVKWIFSSVVDLKQTIIPDYRNMIGQGA**

>DPA1*02:01-DPB1*03:01(DP3)

**MRPEDRMFHIRAVILRALSLAFLLSLRGAGAIK**ADHVSTYAAFVQTHRPTGEFMFEFDEDEQFYVDLDKK
ETVWHLEEFGRAFSFEAQGGLANIAILNNNLNTLIQRSNHTQAANDPPEVTVFPKEPVELGQPNTLICHI
DRFFPPVLNVTWLCNGEPVTEGVAESLFLPRTDYSFHKFHYLTFVPSAEDVYDCRVEHWGLDQPLLKHW**G**
**GGGSGGGGSGGGGS**ENYVYQLRQECYAFNGTQRFLERYIYNREEFVRFDSDVGEFRAVTELGRPDEDYWN
SQKDLLEEKRAVPDRVCRHNYELDEAVTLQRRVQPKVNVSPSKKGPLQHHNLLVCHVTDFYPGSIQVRWF
LNGQEETAGVVSTNLIRNGDWTFQILVMLEMTPQQGDVYICQVEHTSLDSPVTVEWEPKSCDKTHTCPPC
PAPELLGGPSVFLFPPKPKDTLMISRTPEVTCVVVDVSHEDPEVKFNWYVDGVEVHNAKTKPREEQYNST
YRVVSVLTVLHQDWLNGKEYKCKVSNKGLPSSIEKTISKAKGQPREPQVYTLPPSRDELTKNQVSLTCLV
KGFYPSDIAVEWESNGQPENNYKTTPPVLDSDGSFFLYSKLTVDKSRWQQGNVFSCSVMHEALHNHYTQK
SLSLSPGK**ITIFITLFLLSVCYSATVTFFKVKWIFSSVVDLKQTIIPDYRNMIGQGA**

>DPA1*01:03-DPB1*04:01(DP4)

**MRPEDRMFHIRAVILRALSLAFLLSLRGAGAIK**ADHVSTYAAFVQTHRPTGEFMFEFDEDEMFYVDLDKK
ETVWHLEEFGQAFSFEAQGGLANIAILNNNLNTLIQRSNHTQATNDPPEVTVFPKEPVELGQPNTLICHI
DKFFPPVLNVTWLCNGELVTEGVAESLFLPRTDYSFHKFHYLTFVPSAEDFYDCRVEHWGLDQPLLKHW**G**
**GGGSGGGGSGGGGS**ENYLFQGRQECYAFNGTQRFLERYIYNREEFARFDSDVGEFRAVTELGRPAAEYWN
SQKDILEEKRAVPDRMCRHNYELGGPMTLQRRVQPRVNVSPSKKGPLQHHNLLVCHVTDFYPGSIQVRWF
LNGQEETAGVVSTNLIRNGDWTFQILVMLEMTPQQGDVYTCQVEHTSLDSPVTVEWEPKSCDKTHTCPPC
PAPELLGGPSVFLFPPKPKDTLMISRTPEVTCVVVDVSHEDPEVKFNWYVDGVEVHNAKTKPREEQYNST
YRVVSVLTVLHQDWLNGKEYKCKVSNKGLPSSIEKTISKAKGQPREPQVYTLPPSRDELTKNQVSLTCLV
KGFYPSDIAVEWESNGQPENNYKTTPPVLDSDGSFFLYSKLTVDKSRWQQGNVFSCSVMHEALHNHYTQK
SLSLSPGK**ITIFITLFLLSVCYSATVTFFKVKWIFSSVVDLKQTIIPDYRNMIGQGA**

>DPA1*01:03-DPB1*04:02(DP4)

**MRPEDRMFHIRAVILRALSLAFLLSLRGAGAIK**ADHVSTYAAFVQTHRPTGEFMFEFDEDEMFYVDLDKK
ETVWHLEEFGQAFSFEAQGGLANIAILNNNLNTLIQRSNHTQATNDPPEVTVFPKEPVELGQPNTLICHI
DKFFPPVLNVTWLCNGELVTEGVAESLFLPRTDYSFHKFHYLTFVPSAEDFYDCRVEHWGLDQPLLKHW**G**
**GGGSGGGGSGGGGS**ENYLFQGRQECYAFNGTQRFLERYIYNREEFVRFDSDVGEFRAVTELGRPDEEYWN
SQKDILEEKRAVPDRMCRHNYELGGPMTLQRRVQPRVNVSPSKKGPLQHHNLLVCHVTDFYPGSIQVRWF
LNGQEETAGVVSTNLIRNGDWTFQILVMLEMTPQQGDVYTCQVEHTSMDSPVTVEWEPKSCDKTHTCPPC
PAPELLGGPSVFLFPPKPKDTLMISRTPEVTCVVVDVSHEDPEVKFNWYVDGVEVHNAKTKPREEQYNST
YRVVSVLTVLHQDWLNGKEYKCKVSNKGLPSSIEKTISKAKGQPREPQVYTLPPSRDELTKNQVSLTCLV
KGFYPSDIAVEWESNGQPENNYKTTPPVLDSDGSFFLYSKLTVDKSRWQQGNVFSCSVMHEALHNHYTQK
SLSLSPGK**ITIFITLFLLSVCYSATVTFFKVKWIFSSVVDLKQTIIPDYRNMIGQGA**

>DPA1*02:01-DPB1*05:01(DP5)

**MRPEDRMFHIRAVILRALSLAFLLSLRGAGAIK**ADHVSTYAAFVQTHRPTGEFMFEFDEDEQFYVDLDKK
ETVWHLEEFGRAFSFEAQGGLANIAILNNNLNTLIQRSNHTQAANDPPEVTVFPKEPVELGQPNTLICHI
DRFFPPVLNVTWLCNGEPVTEGVAESLFLPRTDYSFHKFHYLTFVPSAEDVYDCRVEHWGLDQPLLKHW**G**
**GGGSGGGGSGGGGS**ENYLFQGRQECYAFNGTQRFLERYIYNREELVRFDSDVGEFRAVTELGRPEAEYWN
SQKDILEEKRAVPDRMCRHNYELDEAVTLQRRVQPKVNVSPSKKGPLQHHNLLVCHVTDFYPGSIQVRWF
LNGQEETAGVVSTNLIRNGDWTFQILVMLEMTPQQGDVYICQVEHTSLDSPVTVEWEPKSCDKTHTCPPC
PAPELLGGPSVFLFPPKPKDTLMISRTPEVTCVVVDVSHEDPEVKFNWYVDGVEVHNAKTKPREEQYNST
YRVVSVLTVLHQDWLNGKEYKCKVSNKGLPSSIEKTISKAKGQPREPQVYTLPPSRDELTKNQVSLTCLV
KGFYPSDIAVEWESNGQPENNYKTTPPVLDSDGSFFLYSKLTVDKSRWQQGNVFSCSVMHEALHNHYTQK
SLSLSPGK**ITIFITLFLLSVCYSATVTFFKVKWIFSSVVDLKQTIIPDYRNMIGQGA**

>DPA1*02:02-DPB1*05:01(DP5)

**MRPEDRMFHIRAVILRALSLAFLLSLRGAGAIK**ADHVSTYAMFVQTHRPTGEFMFEFDEDEQFYVDLDKK
ETVWHLEEFGRAFSFEAQGGLANIAILNNNLNTLIQRSNHTQAANDPPEVTVFPKEPVELGQPNTLICHI
DRFFPPVLNVTWLCNGEPVTEGVAESLFLPRTDYSFHKFHYLTFVPSAEDVYDCRVEHWGLDQPLLKHW**G**
**GGGSGGGGSGGGGS**ENYLFQGRQECYAFNGTQRFLERYIYNREELVRFDSDVGEFRAVTELGRPEAEYWN
SQKDILEEKRAVPDRMCRHNYELDEAVTLQRRVQPKVNVSPSKKGPLQHHNLLVCHVTDFYPGSIQVRWF
LNGQEETAGVVSTNLIRNGDWTFQILVMLEMTPQQGDVYICQVEHTSLDSPVTVEWEPKSCDKTHTCPPC
PAPELLGGPSVFLFPPKPKDTLMISRTPEVTCVVVDVSHEDPEVKFNWYVDGVEVHNAKTKPREEQYNST
YRVVSVLTVLHQDWLNGKEYKCKVSNKGLPSSIEKTISKAKGQPREPQVYTLPPSRDELTKNQVSLTCLV
KGFYPSDIAVEWESNGQPENNYKTTPPVLDSDGSFFLYSKLTVDKSRWQQGNVFSCSVMHEALHNHYTQK
SLSLSPGK**ITIFITLFLLSVCYSATVTFFKVKWIFSSVVDLKQTIIPDYRNMIGQGA**

>DPA1*01:03-DPB1*06:01(DP6)

**MRPEDRMFHIRAVILRALSLAFLLSLRGAGAIK**ADHVSTYAAFVQTHRPTGEFMFEFDEDEMFYVDLDKK
ETVWHLEEFGQAFSFEAQGGLANIAILNNNLNTLIQRSNHTQATNDPPEVTVFPKEPVELGQPNTLICHI
DKFFPPVLNVTWLCNGELVTEGVAESLFLPRTDYSFHKFHYLTFVPSAEDFYDCRVEHWGLDQPLLKHW**G**
**GGGSGGGGSGGGGS**ENYVYQLRQECYAFNGTQRFLERYIYNREEFVRFDSDVGEFRAVTELGRPDEDYWN
SQKDLLEEERAVPDRMCRHNYELDEAVTLQRRVQPKVNVSPSKKGPLQHHNLLVCHVTDFYPGSIQVRWF
LNGQEETAGVVSTNLIRNGDWTFQILVMLEMTPQQGDVYICQVEHTSLDSPVTVEWEPKSCDKTHTCPPC

PAPELLGGPSVFLFPPKPKDTLMISRTPEVTCVVVDVSHEDPEVKFNWYVDGVEVHNAKTKPREEQYNST
YRVVSVLTVLHQDWLNGKEYKCKVSNKGLPSSIEKTISKAKGQPREPQVYTLPPSRDELTKNQVSLTCLV
KGFYPSDIAVEWESNGQPENNYKTTPPVLDSDGSFFLYSKLTVDKSRWQQGNVFSCSVMHEALHNHYTQK
SLSLSPGK**ITIFITLFLLSVCYSATVTFFKVKWIFSSVVDLKQTIIPDYRNMIGQGA**

>DPA1*02:01-DPB1*06:01(DP6)

**MRPEDRMFHIRAVILRALSLAFLLSLRGAGAIK**ADHVSTYAAFVQTHRPTGEFMFEFDEDEQFYVDLDKK
ETVWHLEEFGRAFSFEAQGGLANIAILNNNLNTLIQRSNHTQAANDPPEVTVFPKEPVELGQPNTLICHI
DRFFPPVLNVTWLCNGEPVTEGVAESLFLPRTDYSFHKFHYLTFVPSAEDVYDCRVEHWGLDQPLLKHW**G**
**GGGSGGGGSGGGGS**ENYVYQLRQECYAFNGTQRFLERYIYNREEFVRFDSDVGEFRAVTELGRPDEDYWN
SQKDLLEEERAVPDRMCRHNYELDEAVTLQRRVQPKVNVSPSKKGPLQHHNLLVCHVTDFYPGSIQVRWF
LNGQEETAGVVSTNLIRNGDWTFQILVMLEMTPQQGDVYICQVEHTSLDSPVTVEWEPKSCDKTHTCPPC
PAPELLGGPSVFLFPPKPKDTLMISRTPEVTCVVVDVSHEDPEVKFNWYVDGVEVHNAKTKPREEQYNST
YRVVSVLTVLHQDWLNGKEYKCKVSNKGLPSSIEKTISKAKGQPREPQVYTLPPSRDELTKNQVSLTCLV
KGFYPSDIAVEWESNGQPENNYKTTPPVLDSDGSFFLYSKLTVDKSRWQQGNVFSCSVMHEALHNHYTQK
SLSLSPGK**ITIFITLFLLSVCYSATVTFFKVKWIFSSVVDLKQTIIPDYRNMIGQGA**

>DPA1*02:01-DPB1*09:01(DP9)

**MRPEDRMFHIRAVILRALSLAFLLSLRGAGAIK**ADHVSTYAAFVQTHRPTGEFMFEFDEDEQFYVDLDKK
ETVWHLEEFGRAFSFEAQGGLANIAILNNNLNTLIQRSNHTQAANDPPEVTVFPKEPVELGQPNTLICHI
DRFFPPVLNVTWLCNGEPVTEGVAESLFLPRTDYSFHKFHYLTFVPSAEDVYDCRVEHWGLDQPLLKHW**G**
**GGGSGGGGSGGGGS**ENYVHQLRQECYAFNGTQRFLERYIYNREEFVRFDSDVGEFRAVTELGRPDEDYWN
SQKDILEEERAVPDRVCRHNYELDEAVTLQRRVQPKVNVSPSKKGPLQHHNLLVCHVTDFYPGSIQVRWF
LNGQEETAGVVSTNLIRNGDWTFQILVMLEMTPQQGDVYICQVEHTSLDSPVTVEWEPKSCDKTHTCPPC
PAPELLGGPSVFLFPPKPKDTLMISRTPEVTCVVVDVSHEDPEVKFNWYVDGVEVHNAKTKPREEQYNST
YRVVSVLTVLHQDWLNGKEYKCKVSNKGLPSSIEKTISKAKGQPREPQVYTLPPSRDELTKNQVSLTCLV
KGFYPSDIAVEWESNGQPENNYKTTPPVLDSDGSFFLYSKLTVDKSRWQQGNVFSCSVMHEALHNHYTQK
SLSLSPGK**ITIFITLFLLSVCYSATVTFFKVKWIFSSVVDLKQTIIPDYRNMIGQGA**

>DPA1*02:02-DPB1*10:01(DP10)

**MRPEDRMFHIRAVILRALSLAFLLSLRGAGAIK**ADHVSTYAMFVQTHRPTGEFMFEFDEDEQFYVDLDKK
ETVWHLEEFGRAFSFEAQGGLANIAILNNNLNTLIQRSNHTQAANDPPEVTVFPKEPVELGQPNTLICHI
DRFFPPVLNVTWLCNGEPVTEGVAESLFLPRTDYSFHKFHYLTFVPSAEDVYDCRVEHWGLDQPLLKHW**G**
**GGGSGGGGSGGGGS**ENYVHQLRQECYAFNGTQRFLERYIYNREEFVRFDSDVGEFRAVTELGRPDEEYWN
SQKDILEEERAVPDRVCRHNYELDEAVTLQRRVQPKVNVSPSKKGPLQHHNLLVCHVTDFYPGSIQVRWF
LNGQEETAGVVSTNLIRNGDWTFQILVMLEMTPQQGDVYICQVEHTSLDSPVTVEWEPKSCDKTHTCPPC
PAPELLGGPSVFLFPPKPKDTLMISRTPEVTCVVVDVSHEDPEVKFNWYVDGVEVHNAKTKPREEQYNST
YRVVSVLTVLHQDWLNGKEYKCKVSNKGLPSSIEKTISKAKGQPREPQVYTLPPSRDELTKNQVSLTCLV
KGFYPSDIAVEWESNGQPENNYKTTPPVLDSDGSFFLYSKLTVDKSRWQQGNVFSCSVMHEALHNHYTQK
SLSLSPGK**ITIFITLFLLSVCYSATVTFFKVKWIFSSVVDLKQTIIPDYRNMIGQGA**

>DPA1*01:03-DPB1*11:01(DP11)

**MRPEDRMFHIRAVILRALSLAFLLSLRGAGAIK**ADHVSTYAAFVQTHRPTGEFMFEFDEDEMFYVDLDKK
ETVWHLEEFGQAFSFEAQGGLANIAILNNNLNTLIQRSNHTQATNDPPEVTVFPKEPVELGQPNTLICHI
DKFFPPVLNVTWLCNGELVTEGVAESLFLPRTDYSFHKFHYLTFVPSAEDFYDCRVEHWGLDQPLLKHW**G**
**GGGSGGGGSGGGGS**ENYVYQLRQECYAFNGTQRFLERYIYNRQEYARFDSDVGEFRAVTELGRPAAEYWN
SQKDLLEERRAVPDRMCRHNYELDEAVTLQRRVQPKVNVSPSKKGPLQHHNLLVCHVTDFYPGSIQVRWF
LNGQEETAGVVSTNLIRNGDWTFQILVMLEMTPQQGDVYICQVEHTSLDSPVTVEWEPKSCDKTHTCPPC
PAPELLGGPSVFLFPPKPKDTLMISRTPEVTCVVVDVSHEDPEVKFNWYVDGVEVHNAKTKPREEQYNST
YRVVSVLTVLHQDWLNGKEYKCKVSNKGLPSSIEKTISKAKGQPREPQVYTLPPSRDELTKNQVSLTCLV
KGFYPSDIAVEWESNGQPENNYKTTPPVLDSDGSFFLYSKLTVDKSRWQQGNVFSCSVMHEALHNHYTQK
SLSLSPGK**ITIFITLFLLSVCYSATVTFFKVKWIFSSVVDLKQTIIPDYRNMIGQGA**

>DPA1*02:02-DPB1*11:01(DP11)

**MRPEDRMFHIRAVILRALSLAFLLSLRGAGAIK**ADHVSTYAMFVQTHRPTGEFMFEFDEDEQFYVDLDKK
ETVWHLEEFGRAFSFEAQGGLANIAILNNNLNTLIQRSNHTQAANDPPEVTVFPKEPVELGQPNTLICHI
DRFFPPVLNVTWLCNGEPVTEGVAESLFLPRTDYSFHKFHYLTFVPSAEDVYDCRVEHWGLDQPLLKHW**G**
**GGGSGGGGSGGGGS**ENYVYQLRQECYAFNGTQRFLERYIYNRQEYARFDSDVGEFRAVTELGRPAAEYWN
SQKDLLEERRAVPDRMCRHNYELDEAVTLQRRVQPKVNVSPSKKGPLQHHNLLVCHVTDFYPGSIQVRWF
LNGQEETAGVVSTNLIRNGDWTFQILVMLEMTPQQGDVYICQVEHTSLDSPVTVEWEPKSCDKTHTCPPC
PAPELLGGPSVFLFPPKPKDTLMISRTPEVTCVVVDVSHEDPEVKFNWYVDGVEVHNAKTKPREEQYNST
YRVVSVLTVLHQDWLNGKEYKCKVSNKGLPSSIEKTISKAKGQPREPQVYTLPPSRDELTKNQVSLTCLV
KGFYPSDIAVEWESNGQPENNYKTTPPVLDSDGSFFLYSKLTVDKSRWQQGNVFSCSVMHEALHNHYTQK
SLSLSPGK**ITIFITLFLLSVCYSATVTFFKVKWIFSSVVDLKQTIIPDYRNMIGQGA**

>DPA1*02:01-DPB1*13:01(DP13)

**MRPEDRMFHIRAVILRALSLAFLLSLRGAGAIK**ADHVSTYAAFVQTHRPTGEFMFEFDEDEQFYVDLDKK
ETVWHLEEFGRAFSFEAQGGLANIAILNNNLNTLIQRSNHTQAANDPPEVTVFPKEPVELGQPNTLICHI
DRFFPPVLNVTWLCNGEPVTEGVAESLFLPRTDYSFHKFHYLTFVPSAEDVYDCRVEHWGLDQPLLKHW**G**
**GGGSGGGGSGGGGS**ENYVYQLRQECYAFNGTQRFLERYIYNREEYARFDSDVGEFRAVTELGRPAAEYWN
SQKDILEEERAVPDRICRHNYELDEAVTLQRRVQPKVNVSPSKKGPLQHHNLLVCHVTDFYPGSIQVRWF
LNGQEETAGVVSTNLIRNGDWTFQILVMLEMTPQQGDVYICQVEHTSLDSPVTVEWEPKSCDKTHTCPPC
PAPELLGGPSVFLFPPKPKDTLMISRTPEVTCVVVDVSHEDPEVKFNWYVDGVEVHNAKTKPREEQYNST
YRVVSVLTVLHQDWLNGKEYKCKVSNKGLPSSIEKTISKAKGQPREPQVYTLPPSRDELTKNQVSLTCLV
KGFYPSDIAVEWESNGQPENNYKTTPPVLDSDGSFFLYSKLTVDKSRWQQGNVFSCSVMHEALHNHYTQK
SLSLSPGK**ITIFITLFLLSVCYSATVTFFKVKWIFSSVVDLKQTIIPDYRNMIGQGA**

>DPA1*02:02-DPB1*13:01(DP13)

**MRPEDRMFHIRAVILRALSLAFLLSLRGAGAIK**ADHVSTYAMFVQTHRPTGEFMFEFDEDEQFYVDLDKK
ETVWHLEEFGRAFSFEAQGGLANIAILNNNLNTLIQRSNHTQAANDPPEVTVFPKEPVELGQPNTLICHI
DRFFPPVLNVTWLCNGEPVTEGVAESLFLPRTDYSFHKFHYLTFVPSAEDVYDCRVEHWGLDQPLLKHW**G**
**GGGSGGGGSGGGGS**ENYVYQLRQECYAFNGTQRFLERYIYNREEYARFDSDVGEFRAVTELGRPAAEYWN
SQKDILEEERAVPDRICRHNYELDEAVTLQRRVQPKVNVSPSKKGPLQHHNLLVCHVTDFYPGSIQVRWF
LNGQEETAGVVSTNLIRNGDWTFQILVMLEMTPQQGDVYICQVEHTSLDSPVTVEWEPKSCDKTHTCPPC

PAPELLGGPSVFLFPPKPKDTLMISRTPEVTCVVVDVSHEDPEVKFNWYVDGVEVHNAKTKPREEQYNST
YRVVSVLTVLHQDWLNGKEYKCKVSNKGLPSSIEKTISKAKGQPREPQVYTLPPSRDELTKNQVSLTCLV
KGFYPSDIAVEWESNGQPENNYKTTPPVLDSDGSFFLYSKLTVDKSRWQQGNVFSCSVMHEALHNHYTQK
SLSLSPGK**ITIFITLFLLSVCYSATVTFFKVKWIFSSVVDLKQTIIPDYRNMIGQGA**

>DPA1*03:01-DPB1*13:01(DP13)

**MRPEDRMFHIRAVILRALSLAFLLSLRGAGAIK**ADHVSTYAMFVQTHRPTGEFMFEFDEDEMFYVDLDKK
ETVWHLEEFGQAFSFEAQGGLANIAISNNNLNTLIQRSNHTQATNDPPEVTVFPKEPVELGQPNTLICHI
DKFFPPVLNVTWLCNGELVTEGVAESLFLPRTDYSFHKFHYLTFVPSAEDFYDCRVEHWGLDQPLLKHW**G**
**GGGSGGGGSGGGGS**ENYVYQLRQECYAFNGTQRFLERYIYNREEYARFDSDVGEFRAVTELGRPAAEYWN
SQKDILEEERAVPDRICRHNYELDEAVTLQRRVQPKVNVSPSKKGPLQHHNLLVCHVTDFYPGSIQVRWF
LNGQEETAGVVSTNLIRNGDWTFQILVMLEMTPQQGDVYICQVEHTSLDSPVTVEWEPKSCDKTHTCPPC
PAPELLGGPSVFLFPPKPKDTLMISRTPEVTCVVVDVSHEDPEVKFNWYVDGVEVHNAKTKPREEQYNST
YRVVSVLTVLHQDWLNGKEYKCKVSNKGLPSSIEKTISKAKGQPREPQVYTLPPSRDELTKNQVSLTCLV
KGFYPSDIAVEWESNGQPENNYKTTPPVLDSDGSFFLYSKLTVDKSRWQQGNVFSCSVMHEALHNHYTQK
SLSLSPGK**ITIFITLFLLSVCYSATVTFFKVKWIFSSVVDLKQTIIPDYRNMIGQGA**

>DPA1*02:01-DPB1*14:01(DP14)

**MRPEDRMFHIRAVILRALSLAFLLSLRGAGAIK**ADHVSTYAAFVQTHRPTGEFMFEFDEDEQFYVDLDKK
ETVWHLEEFGRAFSFEAQGGLANIAILNNNLNTLIQRSNHTQAANDPPEVTVFPKEPVELGQPNTLICHI
DRFFPPVLNVTWLCNGEPVTEGVAESLFLPRTDYSFHKFHYLTFVPSAEDVYDCRVEHWGLDQPLLKHW**G**
**GGGSGGGGSGGGGS**ENYVHQLRQECYAFNGTQRFLERYIYNREEFVRFDSDVGEFRAVTELGRPDEDYWN
SQKDLLEEKRAVPDRVCRHNYELDEAVTLQRRVQPKVNVSPSKKGPLQHHNLLVCHVTDFYPGSIQVRWF
LNGQEETAGVVSTNLIRNGDWTFQILVMLEMTPQQGDVYICQVEHTSLDSPVTVEWEPKSCDKTHTCPPC
PAPELLGGPSVFLFPPKPKDTLMISRTPEVTCVVVDVSHEDPEVKFNWYVDGVEVHNAKTKPREEQYNST
YRVVSVLTVLHQDWLNGKEYKCKVSNKGLPSSIEKTISKAKGQPREPQVYTLPPSRDELTKNQVSLTCLV
KGFYPSDIAVEWESNGQPENNYKTTPPVLDSDGSFFLYSKLTVDKSRWQQGNVFSCSVMHEALHNHYTQK
SLSLSPGK**ITIFITLFLLSVCYSATVTFFKVKWIFSSVVDLKQTIIPDYRNMIGQGA**

>DPA1*02:01-DPB1*15:01(DP15)

**MRPEDRMFHIRAVILRALSLAFLLSLRGAGAIK**ADHVSTYAAFVQTHRPTGEFMFEFDEDEQFYVDLDKK
ETVWHLEEFGRAFSFEAQGGLANIAILNNNLNTLIQRSNHTQAANDPPEVTVFPKEPVELGQPNTLICHI
DRFFPPVLNVTWLCNGEPVTEGVAESLFLPRTDYSFHKFHYLTFVPSAEDVYDCRVEHWGLDQPLLKHW**G**
**GGGSGGGGSGGGGS**ENYVYQGRQECYAFNGTQRFLERYIYNRQEYARFDSDVGEFRAVTELGRPAAEYWN
SQKDLLEERRAVPDRMCRHNYELVGPMTLQRRVQPKVNVSPSKKGPLQHHNLLVCHVTDFYPGSIQVRWF
LNGQEETAGVVSTNLIRNGDWTFQILVMLEMTPQQGDVYICQVEHTSLDSPVTVEWEPKSCDKTHTCPPC
PAPELLGGPSVFLFPPKPKDTLMISRTPEVTCVVVDVSHEDPEVKFNWYVDGVEVHNAKTKPREEQYNST
YRVVSVLTVLHQDWLNGKEYKCKVSNKGLPSSIEKTISKAKGQPREPQVYTLPPSRDELTKNQVSLTCLV
KGFYPSDIAVEWESNGQPENNYKTTPPVLDSDGSFFLYSKLTVDKSRWQQGNVFSCSVMHEALHNHYTQK
SLSLSPGK**ITIFITLFLLSVCYSATVTFFKVKWIFSSVVDLKQTIIPDYRNMIGQGA**

>DPA1*02:01-DPB1*17:01(DP17)

**MRPEDRMFHIRAVILRALSLAFLLSLRGAGAIK**ADHVSTYAAFVQTHRPTGEFMFEFDEDEQFYVDLDKK
ETVWHLEEFGRAFSFEAQGGLANIAILNNNLNTLIQRSNHTQAANDPPEVTVFPKEPVELGQPNTLICHI
DRFFPPVLNVTWLCNGEPVTEGVAESLFLPRTDYSFHKFHYLTFVPSAEDVYDCRVEHWGLDQPLLKHW**G**
**GGGSGGGGSGGGGS**ENYVHQLRQECYAFNGTQRFLERYIYNREEFVRFDSDVGEFRAVTELGRPDEDYWN
SQKDILEEERAVPDRMCRHNYELDEAVTLQRRVQPRVNVSPSKKGPLQHHNLLVCHVTDFYPGSIQVRWF
LNGQEETAGVVSTNLIRNGDWTFQILVMLEMTPQQGDVYTCQVEHTSLDSPVTVEWEPKSCDKTHTCPPC
PAPELLGGPSVFLFPPKPKDTLMISRTPEVTCVVVDVSHEDPEVKFNWYVDGVEVHNAKTKPREEQYNST
YRVVSVLTVLHQDWLNGKEYKCKVSNKGLPSSIEKTISKAKGQPREPQVYTLPPSRDELTKNQVSLTCLV
KGFYPSDIAVEWESNGQPENNYKTTPPVLDSDGSFFLYSKLTVDKSRWQQGNVFSCSVMHEALHNHYTQK
SLSLSPGK**ITIFITLFLLSVCYSATVTFFKVKWIFSSVVDLKQTIIPDYRNMIGQGA**

>DPA1*01:04-DPB1*18:01(DP18)

**MRPEDRMFHIRAVILRALSLAFLLSLRGAGAIK**ADHVSTYAAFVQTHRPTGEFMFEFDDDEMFYVDLDKK
ETVWHLEEFGQAFSFEAQGGLANIAILNNNLNTLIQRSNHTQATNDPPEVTVFPKEPVELGQPNTLICHI
DKFFPPVLNVTWLCNGELVTEGVAESLFLPRTDYSFHKFHYLTFVPSAEDFYDCRVEHWGLDQPLLKHW**G**
**GGGSGGGGSGGGGS**ENYVYQGRQECYAFNGTQRFLERYIYNREEFVRFDSDVGEFRAVTELGRPDEEYWN
SQKDILEEKRAVPDRMCRHNYELVGPMTLQRRVQPKVNVSPSKKGPLQHHNLLVCHVTDFYPGSIQVRWF
LNGQEETAGVVSTNLIRNGDWTFQILVMLEMTPQQGDVYICQVEHTSLDSPVTVEWEPKSCDKTHTCPPC
PAPELLGGPSVFLFPPKPKDTLMISRTPEVTCVVVDVSHEDPEVKFNWYVDGVEVHNAKTKPREEQYNST
YRVVSVLTVLHQDWLNGKEYKCKVSNKGLPSSIEKTISKAKGQPREPQVYTLPPSRDELTKNQVSLTCLV
KGFYPSDIAVEWESNGQPENNYKTTPPVLDSDGSFFLYSKLTVDKSRWQQGNVFSCSVMHEALHNHYTQK
SLSLSPGK**ITIFITLFLLSVCYSATVTFFKVKWIFSSVVDLKQTIIPDYRNMIGQGA**

>DPA1*01:05-DPB1*18:01(DP18)

**MRPEDRMFHIRAVILRALSLAFLLSLRGAGAIK**ADHVSTYAAFVQTHRPTGEFMFEFDEDEMFYVDLDKK
ETVWHLEEFGQAFSFEAQGGLANIAILNNNLNTLIQRSNHTQAANDPPEVTVFPKEPVELGQPNTLICHI
DKFFPPVLNVTWLCNGELVTEGVAESLFLPRTDYSFHKFHYLTFVPSAEDFYDCRVEHWGLDQPLLKHW**G**
**GGGSGGGGSGGGGS**ENYVYQGRQECYAFNGTQRFLERYIYNREEFVRFDSDVGEFRAVTELGRPDEEYWN
SQKDILEEKRAVPDRMCRHNYELVGPMTLQRRVQPKVNVSPSKKGPLQHHNLLVCHVTDFYPGSIQVRWF
LNGQEETAGVVSTNLIRNGDWTFQILVMLEMTPQQGDVYICQVEHTSLDSPVTVEWEPKSCDKTHTCPPC
PAPELLGGPSVFLFPPKPKDTLMISRTPEVTCVVVDVSHEDPEVKFNWYVDGVEVHNAKTKPREEQYNST
YRVVSVLTVLHQDWLNGKEYKCKVSNKGLPSSIEKTISKAKGQPREPQVYTLPPSRDELTKNQVSLTCLV
KGFYPSDIAVEWESNGQPENNYKTTPPVLDSDGSFFLYSKLTVDKSRWQQGNVFSCSVMHEALHNHYTQK
SLSLSPGK**ITIFITLFLLSVCYSATVTFFKVKWIFSSVVDLKQTIIPDYRNMIGQGA**

>DPA1*02:01-DPB1*18:01(DP18)

**MRPEDRMFHIRAVILRALSLAFLLSLRGAGAIK**ADHVSTYAAFVQTHRPTGEFMFEFDEDEQFYVDLDKK
ETVWHLEEFGRAFSFEAQGGLANIAILNNNLNTLIQRSNHTQAANDPPEVTVFPKEPVELGQPNTLICHI
DRFFPPVLNVTWLCNGEPVTEGVAESLFLPRTDYSFHKFHYLTFVPSAEDVYDCRVEHWGLDQPLLKHW**G**
**GGGSGGGGSGGGGS**ENYVYQGRQECYAFNGTQRFLERYIYNREEFVRFDSDVGEFRAVTELGRPDEEYWN
SQKDILEEKRAVPDRMCRHNYELVGPMTLQRRVQPKVNVSPSKKGPLQHHNLLVCHVTDFYPGSIQVRWF
LNGQEETAGVVSTNLIRNGDWTFQILVMLEMTPQQGDVYICQVEHTSLDSPVTVEWEPKSCDKTHTCPPC

PAPELLGGPSVFLFPPKPKDTLMISRTPEVTCVVVDVSHEDPEVKFNWYVDGVEVHNAKTKPREEQYNST
YRVVSVLTVLHQDWLNGKEYKCKVSNKGLPSSIEKTISKAKGQPREPQVYTLPPSRDELTKNQVSLTCLV
KGFYPSDIAVEWESNGQPENNYKTTPPVLDSDGSFFLYSKLTVDKSRWQQGNVFSCSVMHEALHNHYTQK
SLSLSPGK**ITIFITLFLLSVCYSATVTFFKVKWIFSSVVDLKQTIIPDYRNMIGQGA**

>DPA1*01:03-DPB1*19:01(DP19)

**MRPEDRMFHIRAVILRALSLAFLLSLRGAGAIK**ADHVSTYAAFVQTHRPTGEFMFEFDEDEMFYVDLDKK
ETVWHLEEFGQAFSFEAQGGLANIAILNNNLNTLIQRSNHTQATNDPPEVTVFPKEPVELGQPNTLICHI
DKFFPPVLNVTWLCNGELVTEGVAESLFLPRTDYSFHKFHYLTFVPSAEDFYDCRVEHWGLDQPLLKHW**G**
**GGGSGGGGSGGGGS**ENYLFQGRQECYAFNGTQRFLERYIYNREEFVRFDSDVGEFRAVTELGRPEAEYWN
SQKDILEEERAVPDRICRHNYELDEAVTLQRRVQPKVNVSPSKKGPLQHHNLLVCHVTDFYPGSIQVRWF
LNGQEETAGVVSTNLIRNGDWTFQILVMLEMTPQQGDVYICQVEHTSLDSPVTVEWEPKSCDKTHTCPPC
PAPELLGGPSVFLFPPKPKDTLMISRTPEVTCVVVDVSHEDPEVKFNWYVDGVEVHNAKTKPREEQYNST
YRVVSVLTVLHQDWLNGKEYKCKVSNKGLPSSIEKTISKAKGQPREPQVYTLPPSRDELTKNQVSLTCLV
KGFYPSDIAVEWESNGQPENNYKTTPPVLDSDGSFFLYSKLTVDKSRWQQGNVFSCSVMHEALHNHYTQK
SLSLSPGK**ITIFITLFLLSVCYSATVTFFKVKWIFSSVVDLKQTIIPDYRNMIGQGA**

>DPA1*03:01-DPB1*20:01(DP20)

**MRPEDRMFHIRAVILRALSLAFLLSLRGAGAIK**ADHVSTYAMFVQTHRPTGEFMFEFDEDEMFYVDLDKK
ETVWHLEEFGQAFSFEAQGGLANIAISNNNLNTLIQRSNHTQATNDPPEVTVFPKEPVELGQPNTLICHI
DKFFPPVLNVTWLCNGELVTEGVAESLFLPRTDYSFHKFHYLTFVPSAEDFYDCRVEHWGLDQPLLKHW**G**
**GGGSGGGGSGGGGS**ENYVYQLRQECYAFNGTQRFLERYIYNREEFVRFDSDVGEFRAVTELGRPDEDYWN
SQKDLLEEKRAVPDRMCRHNYELDEAVTLQRRVQPKVNVSPSKKGPLQHHNLLVCHVTDFYPGSIQVRWF
LNGQEETAGVVSTNLIRNGDWTFQILVMLEMTPQQGDVYICQVEHTSLDSPVTVEWEPKSCDKTHTCPPC
PAPELLGGPSVFLFPPKPKDTLMISRTPEVTCVVVDVSHEDPEVKFNWYVDGVEVHNAKTKPREEQYNST
YRVVSVLTVLHQDWLNGKEYKCKVSNKGLPSSIEKTISKAKGQPREPQVYTLPPSRDELTKNQVSLTCLV
KGFYPSDIAVEWESNGQPENNYKTTPPVLDSDGSFFLYSKLTVDKSRWQQGNVFSCSVMHEALHNHYTQK
SLSLSPGK**ITIFITLFLLSVCYSATVTFFKVKWIFSSVVDLKQTIIPDYRNMIGQGA**

>DPA1*01:03-DPB1*23:01(DP23)

**MRPEDRMFHIRAVILRALSLAFLLSLRGAGAIK**ADHVSTYAAFVQTHRPTGEFMFEFDEDEMFYVDLDKK
ETVWHLEEFGQAFSFEAQGGLANIAILNNNLNTLIQRSNHTQATNDPPEVTVFPKEPVELGQPNTLICHI
DKFFPPVLNVTWLCNGELVTEGVAESLFLPRTDYSFHKFHYLTFVPSAEDFYDCRVEHWGLDQPLLKHW**G**
**GGGSGGGGSGGGGS**ENYLFQGRQECYAFNGTQRFLERYIYNREEFVRFDSDVGEFRAVTELGRPAAEYWN
SQKDILEEKRAVPDRMCRHNYELGGPMTLQRRVQPRVNVSPSKKGPLQHHNLLVCHVTDFYPGSIQVRWF
LNGQEETAGVVSTNLIRNGDWTFQILVMLEMTPQQGDVYTCQVEHTSLDSPVTVEWEPKSCDKTHTCPPC
PAPELLGGPSVFLFPPKPKDTLMISRTPEVTCVVVDVSHEDPEVKFNWYVDGVEVHNAKTKPREEQYNST
YRVVSVLTVLHQDWLNGKEYKCKVSNKGLPSSIEKTISKAKGQPREPQVYTLPPSRDELTKNQVSLTCLV
KGFYPSDIAVEWESNGQPENNYKTTPPVLDSDGSFFLYSKLTVDKSRWQQGNVFSCSVMHEALHNHYTQK
SLSLSPGK**ITIFITLFLLSVCYSATVTFFKVKWIFSSVVDLKQTIIPDYRNMIGQGA**

>DPA1*01:03-DPB1*28:01(DP28)

**MRPEDRMFHIRAVILRALSLAFLLSLRGAGAIK**ADHVSTYAAFVQTHRPTGEFMFEFDEDEMFYVDLDKK
ETVWHLEEFGQAFSFEAQGGLANIAILNNNLNTLIQRSNHTQATNDPPEVTVFPKEPVELGQPNTLICHI
DKFFPPVLNVTWLCNGELVTEGVAESLFLPRTDYSFHKFHYLTFVPSAEDFYDCRVEHWGLDQPLLKHW**G**
**GGGSGGGGSGGGGS**ENYLFQGRQECYAFNGTQRFLERYIYNREEFARFDSDVGEFRAVTELGRPDEEYWN
SQKDLLEEKRAVPDRMCRHNYELVGPMTLQRRVQPRVNVSPSKKGPLQHHNLLVCHVTDFYPGSIQVRWF
LNGQEETAGVVSTNLIRNGDWTFQILVMLEMTPQQGDVYTCQVEHTSLDSPVTVEWEPKSCDKTHTCPPC
PAPELLGGPSVFLFPPKPKDTLMISRTPEVTCVVVDVSHEDPEVKFNWYVDGVEVHNAKTKPREEQYNST
YRVVSVLTVLHQDWLNGKEYKCKVSNKGLPSSIEKTISKAKGQPREPQVYTLPPSRDELTKNQVSLTCLV
KGFYPSDIAVEWESNGQPENNYKTTPPVLDSDGSFFLYSKLTVDKSRWQQGNVFSCSVMHEALHNHYTQK
SLSLSPGK**ITIFITLFLLSVCYSATVTFFKVKWIFSSVVDLKQTIIPDYRNMIGQGA**

>DPA1*01:05-DPB1*28:01(DP28)

**MRPEDRMFHIRAVILRALSLAFLLSLRGAGAIK**ADHVSTYAAFVQTHRPTGEFMFEFDEDEMFYVDLDKK
ETVWHLEEFGQAFSFEAQGGLANIAILNNNLNTLIQRSNHTQAANDPPEVTVFPKEPVELGQPNTLICHI
DKFFPPVLNVTWLCNGELVTEGVAESLFLPRTDYSFHKFHYLTFVPSAEDFYDCRVEHWGLDQPLLKHW**G**
**GGGSGGGGSGGGGS**ENYLFQGRQECYAFNGTQRFLERYIYNREEFARFDSDVGEFRAVTELGRPDEEYWN
SQKDLLEEKRAVPDRMCRHNYELVGPMTLQRRVQPRVNVSPSKKGPLQHHNLLVCHVTDFYPGSIQVRWF
LNGQEETAGVVSTNLIRNGDWTFQILVMLEMTPQQGDVYTCQVEHTSLDSPVTVEWEPKSCDKTHTCPPC
PAPELLGGPSVFLFPPKPKDTLMISRTPEVTCVVVDVSHEDPEVKFNWYVDGVEVHNAKTKPREEQYNST
YRVVSVLTVLHQDWLNGKEYKCKVSNKGLPSSIEKTISKAKGQPREPQVYTLPPSRDELTKNQVSLTCLV
KGFYPSDIAVEWESNGQPENNYKTTPPVLDSDGSFFLYSKLTVDKSRWQQGNVFSCSVMHEALHNHYTQK
SLSLSPGK**ITIFITLFLLSVCYSATVTFFKVKWIFSSVVDLKQTIIPDYRNMIGQGA**

>DPA1*04:01-DPB1*28:01(DP28)

**MRPEDRMFHIRAVILRALSLAFLLSLRGAGAIK**ADHVSTYAAFVQTHRTTGEFMFEFDDDEMFYVDLDKK
ETVWHLEEFGRAFSFEAQGGLANIAILNNNLNIAIQRSNHTQAANDPPEVTVFPKEAVELGQPNTLICHI
DKFFPPVLNVTWLCNGEPVTEGVAESLFLPRTDYSFHKFHYLTFVPSAEDVYDCRVEHWGLDQPLLKHW**G**
**GGGSGGGGSGGGGS**ENYLFQGRQECYAFNGTQRFLERYIYNREEFARFDSDVGEFRAVTELGRPDEEYWN
SQKDLLEEKRAVPDRMCRHNYELVGPMTLQRRVQPRVNVSPSKKGPLQHHNLLVCHVTDFYPGSIQVRWF
LNGQEETAGVVSTNLIRNGDWTFQILVMLEMTPQQGDVYTCQVEHTSLDSPVTVEWEPKSCDKTHTCPPC
PAPELLGGPSVFLFPPKPKDTLMISRTPEVTCVVVDVSHEDPEVKFNWYVDGVEVHNAKTKPREEQYNST
YRVVSVLTVLHQDWLNGKEYKCKVSNKGLPSSIEKTISKAKGQPREPQVYTLPPSRDELTKNQVSLTCLV
KGFYPSDIAVEWESNGQPENNYKTTPPVLDSDGSFFLYSKLTVDKSRWQQGNVFSCSVMHEALHNHYTQK
SLSLSPGK**ITIFITLFLLSVCYSATVTFFKVKWIFSSVVDLKQTIIPDYRNMIGQGA**
